# Female lineages and changing kinship patterns in Neolithic Çatalhöyük

**DOI:** 10.1101/2024.06.23.600259

**Authors:** Eren Yüncü, Ayça Küçükakdağ Doğu, Damla Kaptan, Muhammed Sıddık Kılıç, Camilla Mazzucato, Merve N. Güler, Elifnaz Eker, Büşra Katırcıoğlu, Maciej Chyleński, Kıvılcım Başak Vural, Arda Sevkar, Gözde Atağ, N. Ezgi Altınışık, Fatma Küçük Baloğlu, Defne Bozkurt, Jessica Pearson, Marco Milella, Cansu Karamurat, Şevval Aktürk, Ekin Sağlıcan, Nisan Yıldız, Dilek Koptekin, Sevgi Yorulmaz, Duygu Deniz Kazancı, Ayça Aydoğan, Nergis Bilge Karabulut, Kanat Gürün, Eline M.J. Schotsmans, Jana Anvari, Eva Rosenstock, Jennifer Byrnes, Peter F. Biehl, David Orton, Vendela Kempe Lagerholm, Hasan Can Gemici, Milena Vasic, Arkadiusz Marciniak, Çiğdem Atakuman, Yılmaz Selim Erdal, Emrah Kırdök, Marin Pilloud, Clark Spencer Larsen, Scott D. Haddow, Anders Götherström, Christopher J. Knüsel, Füsun Özer, Ian Hodder, Mehmet Somel

## Abstract

Arguments have long suggested that the advent of early farming in the Near East and Anatolia was linked to a ‘Mother Goddess’ cult. However, evidence for a dominant female role in these societies has been scarce. We studied social organisation, mobility patterns and gendered practices in Neolithic Southwest Asia using 131 paleogenomes from Çatalhöyük East Mound (7100-5950 BCE), a major settlement in Central Anatolia with an uninterrupted occupation and an apparent egalitarian structure. In contrast to widespread genetic evidence for patrilocality in Neolithic Europe, the Çatalhöyük individuals revealed no indication of patrilocal mobility. Analysing genetic kin ties among individuals buried in the same house (co-burials) across 35 Çatalhöyük buildings, we identified close ties concentrated within buildings and among neighbours in Çatalhöyük’s Early period, akin to those in the preceding Pre-Pottery Neolithic in Southwest Asia. This pattern weakened over time: by the late 7th millennium BCE, subadults buried in the same building were rarely closely genetically related, despite sharing similar diets. Still, throughout the site’s occupation, genetic connections within Çatalhöyük buildings were much more frequently connected via the maternal than the paternal line. We also identified differential funerary treatment of female subadults compared to those of males, with a higher frequency of grave goods associated with females. Our results reveal how kinship practices changed while key female roles persisted over one thousand years in a large Neolithic community in western Eurasia.

## Introduction

Social organisation and gender differentiation in prehistoric societies can be difficult to discern from material culture data and are often subjects of controversy. One such long-standing debate has revolved around the existence of a “Mother Goddess” cult in early food-producing societies. This theory was originally inspired by dominant female figurines found at Pottery Neolithic sites across Anatolia and the Aegean, interpreted as deities symbolising fertility or as representatives of matriarchal organisation (*1*, *2*). However, neither material culture nor bioarchaeology has provided additional evidence for female-biased roles in these societies (*3*, *4*).

Recently, genetic data has emerged as a novel source of evidence to study social organisation and mobility patterns in prehistoric societies. The majority of such studies have thus far focused on European Neolithic societies, revealing a predominant picture of genetic kin-based and patrilocal/patrilineal organisation [reviewed in (*5*)], such as generations within village cemeteries frequently connected through the paternal genetic line (*6–8*) or close genetic connections among mainly male burials in megalithic tombs (*9*, *10*). It is unclear, however, if these organisational patterns apply to the Neolithic Near East, where food-producing cultures originally emerged. The limited genetic data published to date from Anatolia has suggested that genetic kinship and sex-biased mobility in these societies may have differed from those in Neolithic European groups, although the results have remained inconclusive due to limited sample sizes (*11*).

Here, we tackle these questions using a comprehensive paleogenomic dataset from Çatalhöyük, a major Pottery Neolithic site in Central Anatolia known for its elaborate symbolism, including vivid wall paintings and diverse array of female figurines (*1*, *12*) (**Figure 1A-C**). The main Çatalhöyük East Mound was occupied uninterrupted through the 7th millennium BCE (7150-5900 BCE) (*13*) and has been divided into Early (7100-6700 BCE), Middle (6700-6500 BCE), Late (6500-6300 BCE), and Final (6300-5950 BCE) periods based on typological changes. The settlement had a relatively large population for its time, with estimations of 500-800 individuals at its peak (*14*) [3500-8000 in earlier work(*15*)]. The neighbouring Çatalhöyük West Mound dates to the early Chalcolithic of Anatolia, 6100-5500 BCE (*16*, *17*). Agriculture and animal husbandry were the main sources of subsistence in both mounds, but hunting of wild animals also continued (*18*, *19*), also reflected in the symbolism (*20*). Neolithic Çatalhöyük has been described as a house-based and relatively egalitarian society, with evidence neither for public buildings nor for systematic socioeconomic inequality among houses, despite the apparent presence of private food storage (*18*, *21*, *22*). Dominant social rules can be observed in the shared patterns of internal architecture within buildings (*21*) and intramural burial patterns, where the dead were buried under house floors while the buildings were still in use, with the adults usually under raised platforms of the central room, and young subadults (e.g. neonates and infants) more frequently near hearths or in storage rooms (*23–26*). The bioarchaeological evidence indicates a generally healthy population, with the presence of large numbers of subadults suggesting high fertility (*27*). Dietary differentiation between sexes was limited, and while interpersonal violence is documented, lethal aggression is not (in contrast to the European Neolithic evidence) (*27*, *28*). Despite this wealth of knowledge about Çatalhöyük, significant questions have remained open, such as its demographic connections with neighbouring populations, possible sex-bias in mobility patterns (*29*), and the nature and extent of genetic kin ties among individuals buried in the same building, which we term co-burials, including the possible roles of maternal versus paternal connections (*30*).

**Figure 1:**
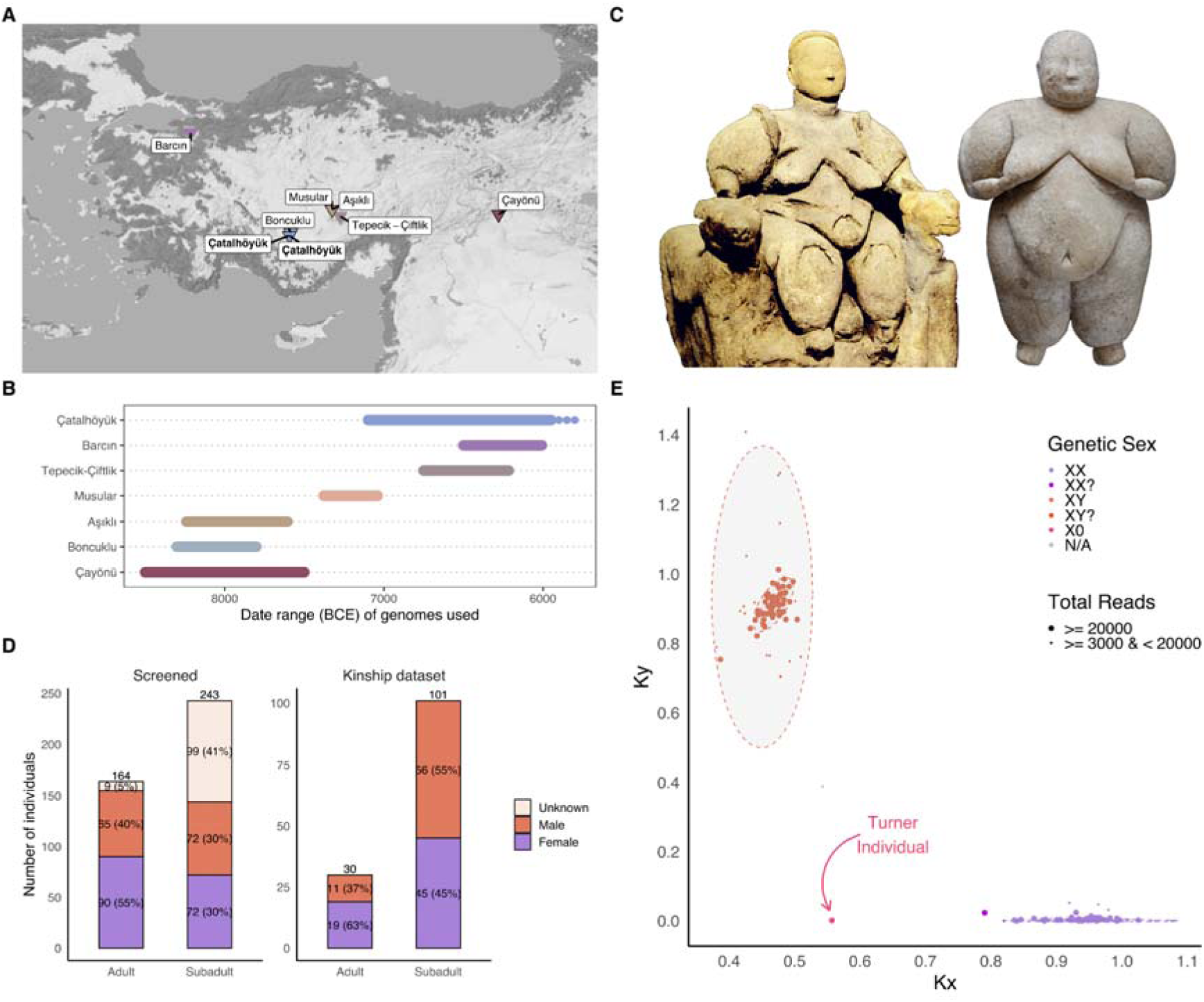
Çatalhöyük, its figurines, and the genomic sample. **A)** Map of Çatalhöyük and Neolithic Anatolian settlements genetically analysed in this study. **B)** Timeline showing the dates of the genetically sampled individuals used in this study. The date range of Çatalhöyük West Mound genomes is shown as dots. **C)** Two renowned adult female figurines from Çatalhöyük. **D)** The distribution of genetically analysed individuals with respect to age and sex. The number of individuals in each category and their percentages are shown inside the bars. **E)** Genetic sex assessment of 179 individuals comparing *K_Y_*(observed/expected chrY reads) and *K_X_* (observed/expected chrX reads) statistics. The neonate with Turner syndrome is indicated with an arrow. The size of points indicates the total number of DNA sequencing reads available per individual, as shown in the figure.

## Results

We genetically screened the skeletal remains of 395 individuals from Çatalhöyük by shallow aDNA sequencing (Methods). Despite poor organic preservation and low endogenous human DNA proportions (median 0.03%), we further shotgun sequenced libraries from 113 new individuals (Methods) and 4 previously published individuals (*31*); together with 17 previously published genomes (*31*) (Koptekin et al. forthcoming; Doğu et al. forthcoming) our final dataset (used in population genetic or kinship analyses) comprised 131 individuals from Çatalhöyük East Mound (c.7100-5900 BCE) and two individuals from the West Mound (c.5900-5800 BCE), with final genomic coverages of 0.001-5.5x (median 0.06x) (**Figure 1D, Figure S1**, **Table S1**). About 60% of the genetically available sample consists of subadults owing to the superior aDNA preservation in subadult skeletal remains, a unique observation we discuss later in this article. We identified the genetic sex of 179 individuals (**Table S1**) in a new approach, *K_XY_* (Methods), with the sex of most subadults determined for the first time (**Figure 1E**). Among the adults, we documented a slightly higher frequency of females (58%) but similar sex ratios among subadults (**Figure 1D-E**). One neonate carried a Turner syndrome (X0) karyotype (**Figure 1E**) (**Supplementary Information**). We used 67 unrelated genomes with >0.03x coverage from the East Mound for demographic analyses (**Table S1**), and 123 for estimating genetic kinship among 5535 pairs (**Table S1; Table S2**). We also imputed two sets of genomes to perform haplotype-based analyses and diploid kinship estimates, 18 with >0.25x coverage, and 49 with >0.1x coverage, respectively (**Table S1**), while keeping track of possible biases that may arise from imputation on 0.1x genomes using downsampling experiments (Methods). We screened all libraries, including 25 from teeth, for pathogenic microbes, but did not identify reliable signatures (Methods; **Supplementary Information**).

### Regional mobility and genetic interactions with neighbours

A central question about Çatalhöyük pertains to mobility dynamics that shaped the community through the 7^th^ millennium BCE as reflected in its gene pool. Whether the site was demographically insular given its conspicuous and persistent cultural characteristics (e.g. the internal architecture of its houses) and also given the lack of large contemporaneous settlements in the region has been a long-standing question (*29*, *32*). The presence of raw materials and products, e.g. from W and E Anatolia, might be considered evidence for wide regional interaction networks (*33*, *34*), although such interactions could have happened without genetic mixing. To investigate evidence for regional gene flow into Çatalhöyük through its occupation, we first explored possible shifts in its average genetic profile over time. MDS and PCA-based clustering of the 67 unrelated Çatalhöyük genomes suggested a relatively homogeneous group similar to other Pottery Neolithic West and Central Anatolian sites (**Figure 2A; Figure S2**); qpAdm modelling also revealed an average Çatalhöyük genetic profile similar to its Anatolian contemporaries (**Figure 2B; Table S3**). This profile, which likely arose through gene flow incomers from U Mesopotamia into local Aceramic/Pre-Pottery Neolithic (PPN) populations by the mid-8th millennium BCE(*35*), appeared stable across Çatalhöyük’s 1000-year occupation (**Figure 2B**). Allelic or haplotype diversity levels were also stable over time, implying no major gene flow event involving genetically distant populations (**Figure S3; Table S2; Supplemental Information**). Meanwhile, individuals from Çatalhöyük’s Early period were genetically more closely related to other Early period Çatalhöyük individuals than those from other periods, as measured using *f*_4_-statistics; this pattern was substantially weaker in the Middle or Late periods (**Figure 2C; Table S4**). This apparent loss of genetic homogeneity over time could be driven by immigration, a pattern we replicate in population genetic simulations (**Supplemental Information**). Indeed, using *f*_4_-statistics, we found that Çayönü genomes (8th mil. BCE U Mesopotamia) show higher genetic affinity to Early period Çatalhöyük than to later periods, whereas Barcın genomes (7th mil. BCE NW Anatolia) show higher affinity to Late period Çatalhöyük than to earlier periods (**Figure S4; Table S4; Supplemental Information**). Together, these results suggest subtle temporal shifts in the Çatalhöyük gene pool through gene flow.

**Figure 2:**
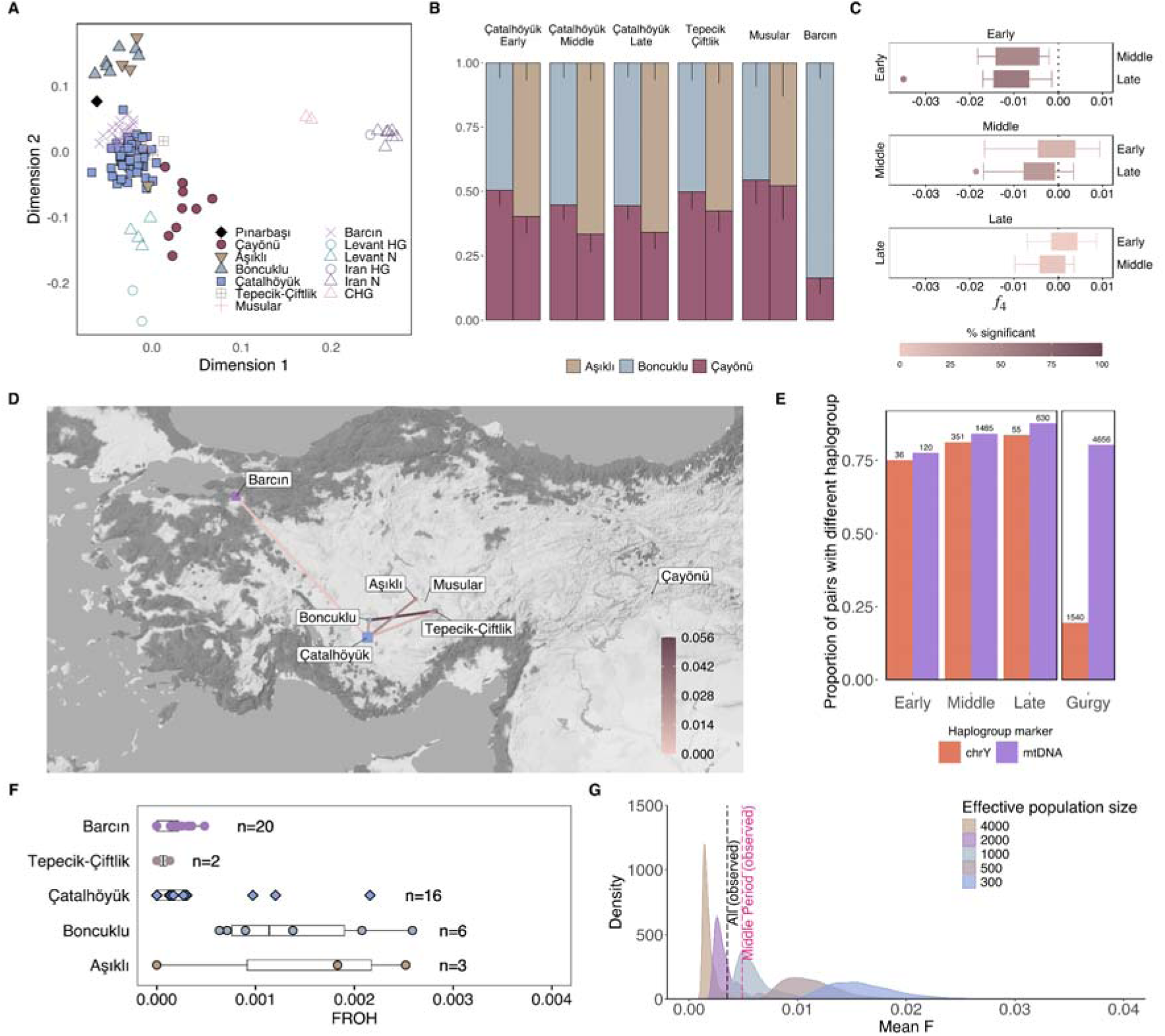
Characterising the Çatalhöyük East Mound gene pool and temporal change. **A)** Multidimensional scaling (MDS) plot summarising outgroup *f*_3_-based genetic distances among late Upper Pleistocene and early Holocene genomes from Southwest Asia, including 67 unrelated Çatalhöyük genomes. **B)** qpAdm modelling of ancestry sources (shown in colours) 8th and 7th millennium Neolithic Anatolian genomes from Çatalhöyük (three periods), Tepecik-Çiftlik, Musular, and Barcın. Each column indicates a feasible model, with the y-axis showing admixture proportions. **C)** *f*_4_-statistics between groups of genomes from the three Çatalhöyük periods. **D)** IBD-sharing with genetically sampled PPN and PN settlements from Anatolia. The colour represents the relative strength of IBD-sharing between two settlements, calculated as the total number of segments shared between all pairwise comparisons divided by the total number of comparisons and the maximum sharing between any pairs in the full sample. **E)** Mitochondrial DNA and Y chromosome haplogroup diversity in Çatalhöyük and French Neolithic Gurgy (*8*), the latter representing a gene pool shaped by patrilocal practices. The numbers of total pairs in each category is indicated on the bars. The difference in chrY diversities between Gurgy and Çatalhöyük is significant as measured by a random subsampling experiment (**Figure S7**). **F)** Comparison of the F_ROH_ values, the inbreeding coefficient estimated using runs of homozygosity (ROH) >4cM, among different Neolithic sites in Anatolia. **G)** The distribution of mean F (inbreeding coefficient) values in a sample of 16 individuals estimated using genealogy simulations under various breeding population sizes shown in the key and the observed mean F_ROH_ values for the 16 Çatalhöyük individuals tested (vertical grey dashed line) and for the 10 individuals from the Çatalhöyük Middle period only (vertical pink dashed line).

We next investigated regional mobility at the individual level, seeking genetic outliers in Çatalhöyük and other Anatolian Neolithic sites using both clustering methods (MDS and PCA) (**Figure 2A; Figure S2**) and formal tests (*f*_4_ and qpWave) (**Figure S4, Table S4, Table S5, Supplementary Information**). Although 28 of 67 Çatalhöyük individuals showed divergent trends in one of five different tests, none displayed consistent affinity to populations of distant regions (e.g. the Levant, Zagros, or Balkans) across multiple analyses **(Figure S5, Table S6**) (for the sake of comparison: one Aşıklı (*31*) individual showed higher affinity to the Levant than to its compatriots in all five tests). However, possible mobility into Çatalhöyük could have involved genetically similar groups from the region. We indeed found that 5 in 11 Middle and 1 in 3 Late period Çatalhöyük individuals with imputed genomes shared 12-16cM identical-by-descent (IBD) segments (*36*) with various genetically sampled Neolithic Anatolian communities, representing distant (e.g. >10 generations) relatedness (**Figure 2D, Figure S6; Table S7, Table S8**). More interestingly, in *f*_4_-statistics, 5/67 Çatalhöyük individuals (subadults from Middle and Late periods) appeared more closely related to contemporaneous Barcın than to other Çatalhöyük genomes **(Table S4)**, again suggesting mobility into Çatalhöyük from beyond Central Anatolia. Our results are compatible with the strontium isotopic data which reported distinct profiles in 7 out of 77 individuals sampled, attesting to inter-regional mobility (*29*). Overall, Çatalhöyük was likely not insular but received non-negligible levels of incomers mainly from proximate and hence genetically alike communities, e.g. those in Central and West Anatolia (Koptekin et al. 2024), who mixed with the village population.

### No evidence for patrilocal mobility in Çatalhöyük

The observation that Neolithic Çatalhöyük received notable gene flow during its occupation raises the question of whether this might be sex-biased. Ethnographic data suggests farming societies, on average, tend to be more patrilocal than foragers (*37*). Strontium and genetic evidence from European Neolithic societies have also frequently reported patrilocality, also termed female exogamy (*6–8*, *38–40*) [reviewed by Bentley(*5*); also see Hrncir and colleagues (*41*) for alternative interpretations].

If mobility into Çatalhöyük was also shaped by female exogamy, this should cause divergent haplogroup diversity levels between chromosome Y (chrY) and mitochondrial DNA (mtDNA). We found comparable chrY and mtDNA diversity levels in all three Çatalhöyük periods (**Figure 2E**). In contrast, the same analysis on a comparable-sized archaeogenetic dataset from Gurgy, a Middle Neolithic burial group in France (*8*), revealed much lower chrY than mtDNA diversity (**Figure 2E**). This is in line with pedigree reconstructions from Gurgy, indicating patrilocality. The Gurgy chrY pattern was also significantly different (p<0.001) from the Çatalhöyük profiles, as we demonstrate in a randomisation test (**Figure S7**). Together with evidence of adult females buried with putative nephews in Çatalhöyük (see below) and in other Neolithic Anatolian sites (*11*, *31*, *42*), our results imply that patrilocal-like traditions observed in the later-coming European may have been inconspicuous, if not absent, in Neolithic Anatolia.

### Low consanguinity in Çatalhöyük suggests active inbreeding avoidance

Comparisons of modern-day mobile forager versus food-producing groups indicate a tendency towards a higher frequency of consanguinity among food producers, possibly owing to social dynamics shaped by property relationships (*43*, *44*). Genetic data from past societies also supports this general conclusion (*45*). We asked if the Çatalhöyük community also practised consanguinity. We estimated ROH >4cM in 16 Çatalhöyük genomes with >0.3x coverage with hapROH (*46*) (Methods). Çatalhöyük homozygosity levels measured with the F_ROH_ statistic were overall lower than estimates from PPN Anatolian communities but comparable with contemporaneous Pottery Neolithic (PN) groups (**Figure 2F, Table S9**). This PPN-PN difference can be attributed to increased genetic diversity in the Neolithic Anatolian gene pool over time owing to the 8th millennium BCE gene flow event mentioned earlier (**Figure 2B**). More interestingly, of the tested Çatalhöyük individuals, only 9 of 16 carried ROH >4cM, with the most extreme case a possible offspring of second cousin mating (**Figure S8**). This indicates relatively low levels of inbreeding in Çatalhöyük, as in other Neolithic Anatolian groups.

The finding of low inbreeding raises the possibility of active inbreeding avoidance in Çatalhöyük. We asked whether the observed F_ROH_ levels might be compatible with random mating under various breeding population sizes (N_b_) without active inbreeding avoidance. We performed genealogy simulations and estimated the inbreeding coefficient (F) assuming a range of N_b_ values (300-4,000) and possible reproductive skew, allowing random mating within generations but excluding sibling and half-sibling unions (Methods). We found that the observed F_ROH_ levels could be attained without active inbreeding avoidance as long as N_b_ was >1000, and more likely, ≥2000 (**Figure 2G)**. Our results can therefore be explained by either Çatalhöyük N_b_ being >1000 and not actively avoiding consanguinity, or, Çatalhöyük N_b_ being <1000 and the community engaging in exogamous reproductive practices (*29*), e.g. by choosing reproductive partners from outside and/or avoiding consanguinity by tracing biological family lineages (**Figure S9, Figure S10**). Given recent estimates of a maximum of c.300 breeding-age adults in Çatalhöyük (*14*), the latter scenario may be more likely.

### Çatalhöyük genetic kin networks

The common practice of intramural subfloor burials in Neolithic SW Asia presents a unique opportunity to study the genetic relationships among individuals connected by a shared social attribute, namely, having been interred in the same building during its occupation. These individuals, while living, may have also been socially and genetically linked with the building inhabitants, as implied by the memory of their burials (*47–50*). Our earlier work in 9th millennium BCE PPN Anatolian settlements suggested that co-burials were frequently genetically closely related (*31*, *42*). In contrast, samples from the PN sites of Çatalhöyük and Barcın found that close genetic ties between spatially connected burials were fewer, suggesting changing practices over time, albeit with much-limited data (*31*); analysis of dental features and mtDNA had also pointed in the same direction (*30*, *51*).

We estimated genetic relatedness between 123 individuals (7503 pairs, 5535 of which had sufficient overlapping SNPs) using multiple tools and approaches simultaneously to maximise the data without compromising accuracy (e.g. using an expanded SNP panel, using imputed diploid genotypes for 49 genomes in genetic kin estimation, and a multi-tiered genetic kin classification system) (Methods). We also analysed the data using different thresholds and included fuzzy categories (e.g. “second- or third-degree”) to ensure the robustness of these results and reflect the limitations of our inference (Supplementary Information). This yielded a network of 133 first- to third-degree kin pairs (2.9% of all tested pairs), connecting 108 individuals across 30 buildings in total (23 of these buildings with multiple burials) (**Figure 3, Figure S11**). Both adult-adult and subadult-subadult pairs were connected at similar rates within buildings (∼30%), although adults were slightly more connected across buildings (5% vs. 2%) (**Table S10**).

**Figure 3:**
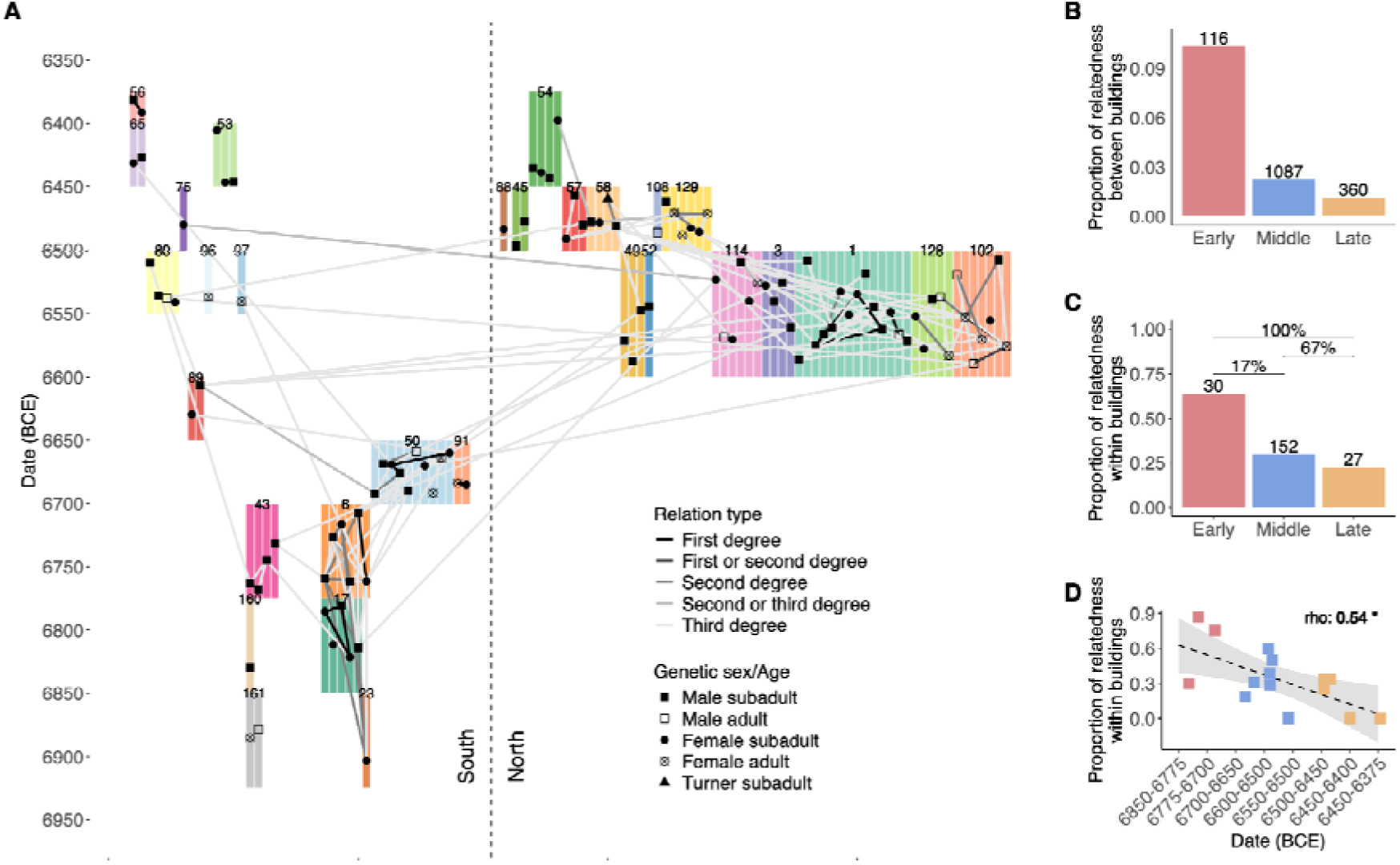
A network of genetic kin across Çatalhöyük buildings and the changing frequency of genetic ties among co-burials. **A)** The figure shows genetic relatedness among intramural burials with common SNPs >3000, shown as dots. The lines show close to more distant relationships from dark to light colours. The coloured blocks show buildings, with building numbers assigned by the excavation team indicated adjacent to the blocks. The height of the blocks is proportional to the number of genetically represented burials in that building. North and South refer to the two main excavation areas on the mound separated by ∼200m. **B)** The proportion of genetic kin (up to third-degree) identified between individuals buried in different buildings in the three Çatalhöyük periods. **C)** The proportion of genetic kin (up to third-degree) identified in co-burials within the same building, separated into three Çatalhöyük periods. See also **Figure S9** for the same proportion calculated for different sets of relatives and age groups. The analysis involves 23 buildings. Percentages indicated on the horizontal bars show the percentage of Monte Carlo simulations where the null hypothesis of no difference between a pair of periods was rejected (p<0.05) out of 24 scenarios involving various assumptions/conditions. The overall rate of rejection among all 108 comparisons was 69% (see **Figure S12** for details). **D)** The proportion of genetic kin within co-burials in 15 buildings (only including buildings with a minimum of 2 burials). The inset shows the Spearman correlation coefficient (p=0.03).

All first-degree pairs were found within buildings (**Figure 3**). Second- and third-degree connections also tended to be spatially concentrated, either within buildings, between successive buildings in the same location, or between buildings in the same area and period, albeit not exclusively (**Figure 3; Table S11**). Inter-building connections were most frequent in the Early period (15%) but declined in the Middle and Late periods (5% and 2%, respectively), consistent with *f*_4_-statistics indicating higher homogeneity in Early period Çatalhöyük (**Figure 3B**). The intensity of interbuilding genetic kin networks also appears correlated with material culture similarity (*52*) (Mazzucato et al., forthcoming). Despite the sparsity of the network, we reconstructed several multi-generational pedigrees, including adult female pairs with possible mother-sibling or avuncular relationships (**Figure 4A-B; Figure S11**).

**Figure 4:**
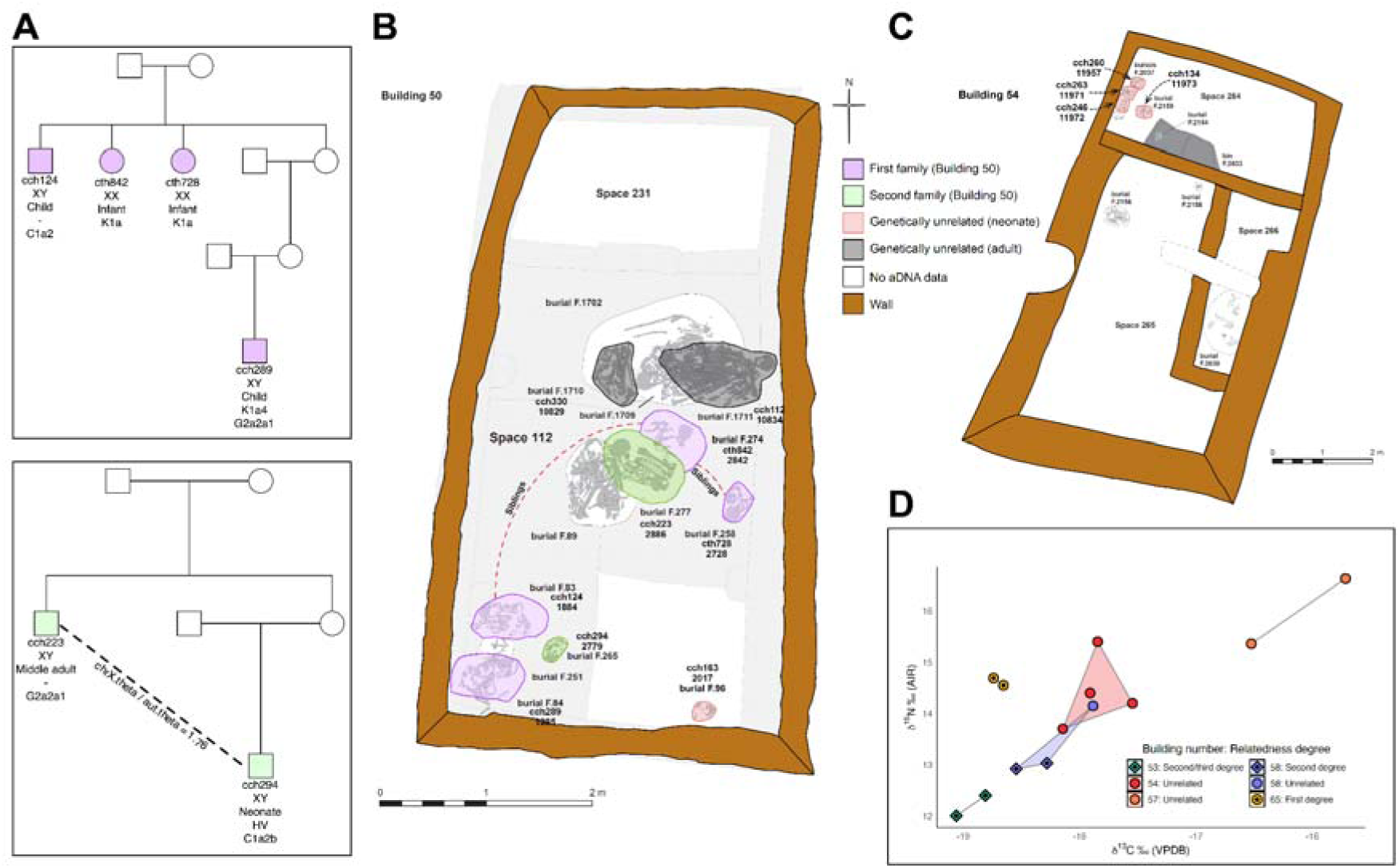
Genetic kin ties and diet among Çatalhöyük co-burials. **A)** Two reconstructed pedigrees of burials inside Building 50 (Middle period), and **B)** their burial location inside the building. Circles show female and squares show male individuals. In the pedigrees, the lab ID, sex, approximate age at death, mitochondrial haplogroup, and Y chromosomal haplogroup, of each genetically studied individual is indicated, respectively. (-) indicates no reliable haplogroup information. On the building plan, individuals are indicated with their lab and excavation unit IDs; grey colouring indicates the individuals were not close genetic kin with the other individuals. The observations that the ratio between chrX and autosomal kinship coefficients (θ) between individuals with IDs 2779 and 2886 was ∼1.8 and that these individuals are carried different Y haplogroups suggests a maternal genetic connection. **C)** Genetically unrelated burials in Building 54 (Late period). **D)** Comparison of carbon and nitrogen stable isotope ratios among neonates interred in Çatalhöyük’s Late period buildings.

### Co-burial of genetically unrelated individuals increases over time

Interestingly, we noticed a temporal change in the density of genetic connections among burials in the same building. Specifically, co-buried pairs inside Early-period buildings were frequently third-degree (e.g., cousin) or closer genetic kin (63%) (**Figure 3**), a pattern similar to those of PPN Anatolian settlements such as Aşıklı, Boncuklu, and Çayönü(*31*, *42*). In contrast, genetic kin among co-buried pairs was only at 30% and 22% frequencies in the Middle and Late periods, respectively (**Figure 3, Figure S12**). We also calculated the correlation between co-burial genetic kin frequencies per building and the building ages and found a similar decrease in time (Spearman correlation r=0.58, p=0.03) (**Figure 3D**). Notably, the only individuals recovered from Chalcolithic Çatalhöyük West, two neonates interred in the infill of the same building (Byrnes and Anvari, forthcoming), were similarly genetically unrelated (Doğu et al., forthcoming). Closer inspection revealed that the Middle and Late-period buildings frequently either accommodated multiple biological families interred together (e.g., Building 50 in **Figure 4A-B**) or burials with no genetic kin ties (e.g. **Figure 4C**) (also see **Figure S12**). The predominance of subadults in the sample renders this observation even more intriguing because if buildings were strictly used by nuclear family groups for burial, then co-burials could include unrelated adults but children should all be related.

We asked whether technical factors could account for the observed temporal change in co-burial genetic ties, such as time variation in DNA preservation, the number of total burials or genetically sampled burials, or the frequency of subadults per building. There was no difference in endogenous DNA proportion or genome coverage among periods (Kruskal-Wallis test p>0.25). To account for variation in sample sizes, we performed Monte-Carlo simulations where we assigned burials within 23 buildings to hypothetical biological families of fixed or variable sizes (n=2-6) under the null hypothesis of no difference among periods, leading to 108 scenarios depending on the genetic kin definitions, hypothetical family sizes used, and whether we included all individuals or only subadults (**Figure S13)**. Under each scenario, we randomly sampled burials from the simulated sets 10,000 times each, and determined if the observed co-burial genetic kin frequency differences among periods could be replicated in the simulations (Supplementary Methods). This revealed that the difference among the periods was overall unexpected, with p<0.05 in 74 out of 108 scenarios (69%) **(Table S12; Figure S13**).

### Share dietary signatures among unrelated neonates

The co-burial of genetically unrelated individuals, the majority of whom were neonates and infants, is intriguing. These individuals may have been unconnected during their lifetime but chosen for burial in the same houses for another reason. Indeed, the frequency of non-primary burials increases over time in Çatalhöyük (**Figure S14**) (Methods) (**Supplementary Information**), representing a practice that may have served to construct or consolidate inter-community ties in Neolithic communities (*53*). Alternatively, these co-buried but genetically unrelated pairs may also have been connected during their lifetimes, such as sharing diets or being raised by foster families (*47–50*).

In order to study this question in depth, we leveraged dietary isotope data collected from neonates (*54*), focusing on five buildings from the Late period with multiple neonate burials (Methods) (**Figure S15, Table S13**). Neonate isotopic profiles could reflect the diet of the individuals carrying the child during pregnancy and providing breast milk or even catabolism in its absence, as well as the duration of feeding (*54*). We found significant differences in isotopic signature among neonates buried in different buildings (Kruskal-Wallis test p<0.02). However, only three of the buildings contained genetic kin, whereas neonates buried in two buildings were genetically unrelated, and one building contained both relatives and an unrelated individual (**Figure 4D**). This record suggests that fostering or other social mechanisms created shared environments for genetically unrelated individuals, and such practices gained prominence over time in Çatalhöyük.

### Predominance of maternal genetic connections among Çatalhöyük co-burials

The pedigrees we constructed revealed an interesting trend: intergenerational connections within buildings frequently running through female lines (**Figure 4; Figure S11**). To study this systematically, we compared mtDNA and chrY haplogroup similarity within buildings (Methods). mtDNA haplogroup homogeneity within buildings was conspicuously higher than between buildings, a pattern maintained across the site’s occupation (**Figure 5A**). In contrast, the chrY homogeneity was relatively low and comparable within and between buildings. We further studied this result using comparisons of autosomal and chrX kinship coefficients (θ). We found that chrX kinship between pairs from the same building was higher than autosomal kinship, while there was no difference between chrX and autosomal kinship coefficients when considering pairs from different buildings.

**Figure 5:**
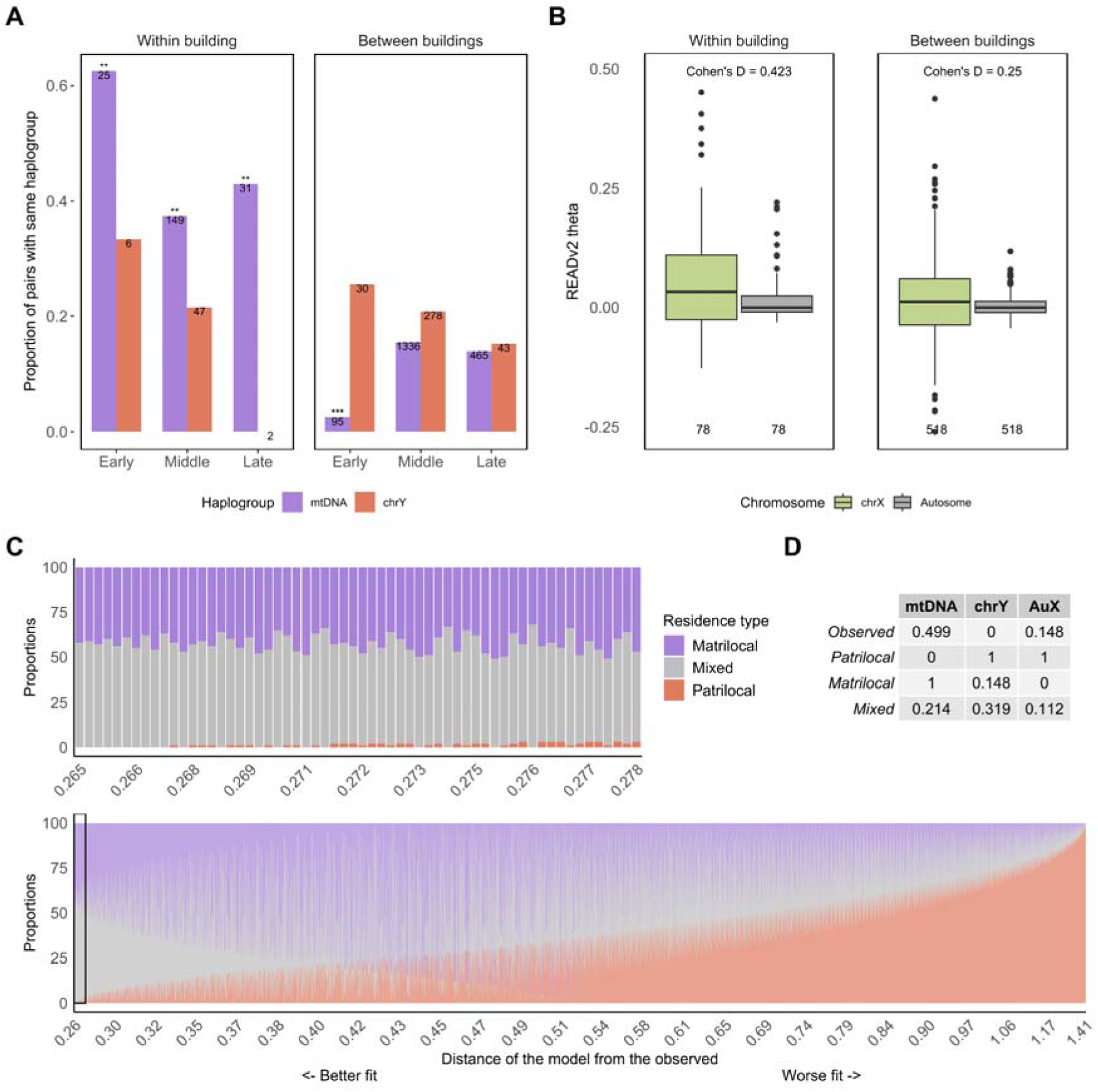
Maternal genetic connections in Çatalhöyük buildings. **A)** mtDNA and chrY haplogroup homogeneity estimates for all pairs buried within the same building (left) and between different buildings (right). The analysis was limited to pairs from the same building. The numbers inside the bars show the total number of pairs in that category. (***) indicates that the observed homogeneity was significantly (p<0.001) different from random expectation, (**) indicates that the observed homogeneity was significantly (p<0.01) different from random expectation, tested by randomly assigning individuals to buildings within the same period. **B)** Autosomal and chrX kinship coefficient (θ) estimates for all pairs buried within the same building (left) and between different buildings (right). The Cohen’s D statistic shows the effect size between the two sets. **C)** Modelling the observed sex-biased diversity patterns using residence model simulations of matrilocal, patrilocal and mixed mobility. Matrilocal here describes the maternal line remaining within buildings across generations and males moving (analogous to matrilineality). We calculated all possible weighted averages of the mean summary statistics (i.e. chrY homogeneity, mtDNA homogeneity, autosomal vs. chrX θ differences). The weights (or proportions) are shown on the y-axis. We then calculated the Euclidean distance (x-axis) between each vector of weighted averages and the observed values. Each bar represents one set of combinations. The model’s fit to the observed genetic data increases towards the left. The upper part of Panel C is a zoomed-in version of the section within the rectangle in the lower part. **D)** The mean summary statistics calculated for each scenario and the observed values are shown in the inset table; each vector was normalised to ensure an equal contribution of each summary statistic to the final result.

To evaluate this evidence further, we performed pedigree simulations of villages and 4-6 generation households with 4-6 offspring per generation, under strict matrilocality, strict patrilocality, and mixed residence (note that here we study mobility among buildings, not among settlements; matrilocality thus corresponds to matrilineality) (Methods). We generated summary statistics (mtDNA and chrY homogeneity, and autosomal vs. chrX kinship coefficient differences) (**Figure S16A-C**). Comparing the summary statistics from the observed data with those from the simulations, we could best explain the observed data as ∼50% mixed and ∼50% matrilocal residence but inferred practically no contribution of strictly patrilocal residence (**Figure 5C**). These results suggest that whenever co-burials were genetically connected, the connections were biased towards the matriline.

### An age-specific burial practice in Çatalhöyük

We had previously reported a trend towards higher aDNA preservation in juvenile temporal bones versus adult temporal bones at Çatalhöyük(*31*). We replicated this observation using 248 subadult and 166 adult samples (Mann-Whitney U test p<0.001). Separating the sample into further age subcategories revealed a remarkable monotonic decline in endogenous proportion with individual age, with two orders of magnitude difference in median endogenous human DNA proportion between newborns and adults (Spearman correlation r=-1, p<0.001) (**Figure 6A**). This result is surprising, given the naive expectation that mature and thus less porous bones of adults may better preserve DNA. We detected a similar but non-significant trend using published and unpublished data in Aşıklı Höyük, while not in other settlements such as Çayönü or Gurgy, denoting that this was not a universal pattern (**Figure S17**). Meanwhile, a scan of organic matter preservation across 111 Çatalhöyük bone samples using Fourier Transform Infrared Spectroscopy (FTIR) identified the same pattern of age-related decline in preservation (Methods) (**Figure 6B; Table S1; Table S14**).

**Figure 6:**
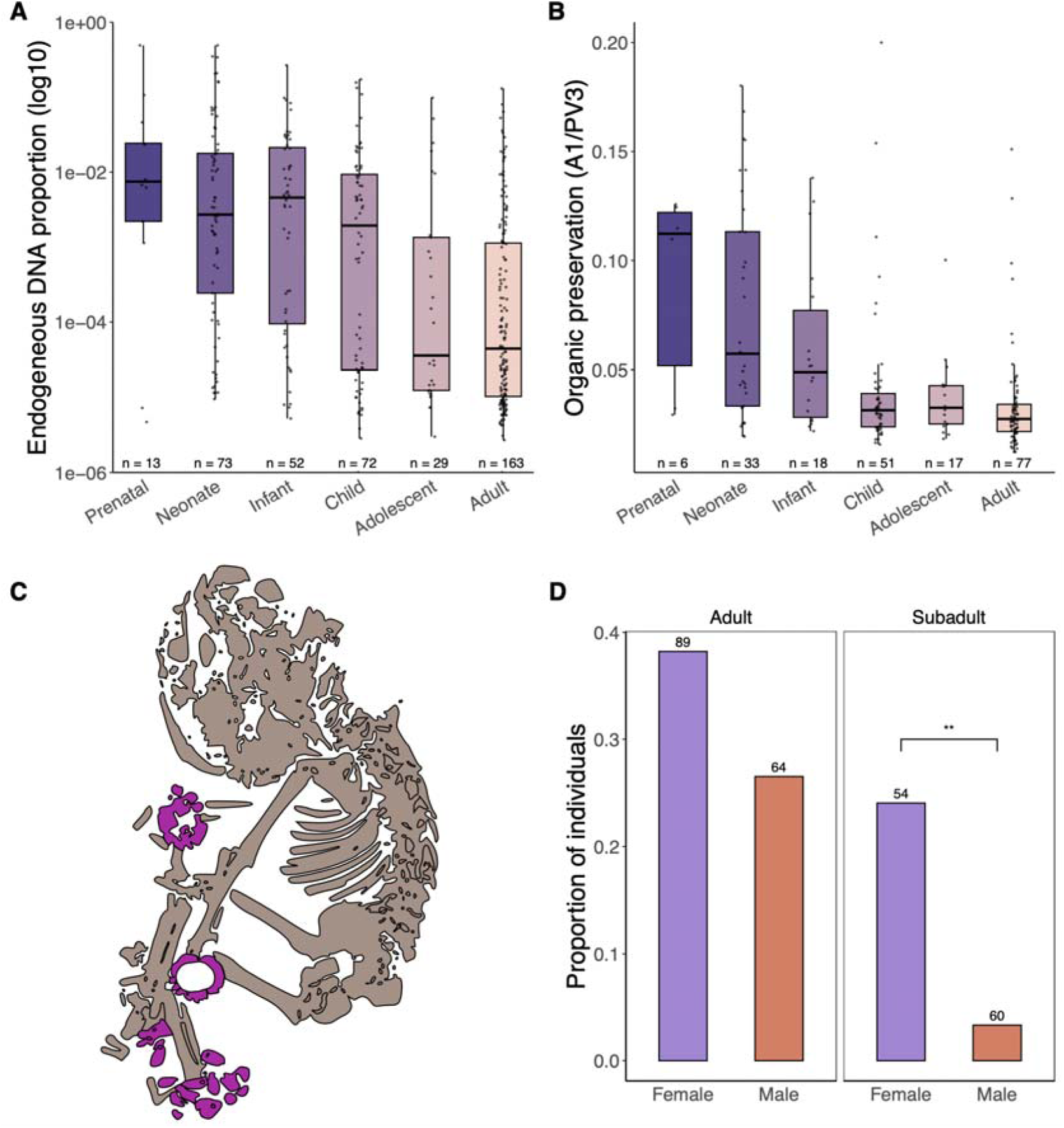
Age- and sex-specific burial practices. **A)** Endogenous DNA yield difference between all age subcategories. **B)** Amide I (A1)/Phosphate (PV3) ratios for assessing the relative organic content preserved in the bones between all age subcategories. **C)** Representation of an infant burial from Building 6 with multiple bone and coloured stone bead ornaments (artificially coloured purple in this image). The individual was not genetically sampled. **D)** Comparison of the inclusion frequencies of burial objects for male and female burials among adults and subadults. The numbers on the bars indicate the total sample size (**: Fisher’s exact test p<0.01).

We hypothesised that this age effect could reflect the outcome of a specific treatment of the corpse or burial practices affecting skeletal preservation, which was differentially applied to the dead of different ages. The effect was observed across all Çatalhöyük periods (**Figure S18A**), suggesting that the practice persisted across centuries. We sought possible correlates of this differential preservation pattern, asking whether the burials with low versus highly preserved aDNA, a) were identified as primary, disturbed, or secondary burials, b) showed signs of extreme flexion, and c) differed in their burial location (**Figure S18B-C-D**) (**Supplementary Information**). We found a correlation between aDNA preservation only with burial location, with lower preservation among subadult burials in the platform area (Mann-Whitney U test p<0.001) (**Figure S18D**). Still, burial location may not be the main reason for the preservation difference; we instead speculate that a body treatment procedure confounded with burial location choices, such as defleshing, drying or similar treatments of selected bodies could have expedited decomposition and DNA degradation (*55*), e.g. by boosting microbial activity. A previous histotaphonomic study carried out on a subset of individuals from Çatalhöyük had failed to detect any bacterial bioerosion (*56*). We nonetheless performed a comparative metagenomic analysis of 40 Çatalhöyük aDNA libraries with the best and worst human DNA preservation (Supplementary Information), which did identify systematic differences in microbial DNA abundance between the two groups. We further found a higher frequency of aerobic microbes in the low-preserved (likely more air-exposed) bones as opposed to a higher frequency of anaerobes in high-preserved (likely immediately interred) bones (**Figures S19**). It is not yet possible to strictly identify whether these microbial composition differences reflect causal or secondary effects. Still, together with multiple reports on the defleshing and processing of adult Çatalhöyük corpses (*1*, *24*, *57*), our results suggest that the age-specific application of funerary treatments on bodies were behind the observed pattern of DNA preservation differences.

### Gender differentiation among subadult burials

Among organic preservation differences within Çatalhöyük age groups, we noted non-significant trends for female children and adolescents to have lower endogenous DNA proportions than male peers (**Figure S18E**), which, if true, could suggest their treatment was more akin to those of adults. We further investigated possible sex differences involving another burial practice, the placement of artefacts such as beads, shells, pigments or stone tools in grave pits (*58*, *59*) (**Figure 6C).** Previous work had found a slightly higher frequency of burial objects associated with adult female burials, but the difference was not statistically significant. In contrast, using subadults genetically sexed in this study, we found a five-fold higher frequency of female burials containing objects than male burials (Fisher’s exact test p<0.01) **(Figure 6D; Table S15**). This result suggests that Çatalhöyük female subadults may have received more elaborate treatment than their male counterparts upon death, which could be related to the linking role of the female lineages across generations.

## Conclusion

The new genetic evidence unveils novel features of Çatalhöyük funerary practices and social organisation, some likely shared with other Neolithic Anatolian sites and others that might be unique to Çatalhöyük. For instance, we find no indication of patrilocal residence in Çatalhöyük. The sparse evidence from other Neolithic Anatolian settlements also points in the same direction, both genetically (*11*) and isotopically (*29*). Hence, mobility might have been bilateral or matrilocal in Neolithic Anatolia, which could be a continuation of earlier forager traditions in this region (*60*). We further speculate that the patrilocal traditions identified in Europe after the 6th millennium BCE emerged after the farming expansion across that continent, either through cultural drift or in response to social stress (*61*). Meanwhile, a putative burial treatment identified at Çatalhöyük reserved for adults (and perhaps some female subadults) that influenced body decomposition appears (for now) without parallel. Our data also revealed a surprising shift in Çatalhöyük funerary traditions and social relations during its occupation: in the Early period co-burials were frequently genetic kin, similar to Pre-Pottery Neolithic Anatolian settlements, while burials (mostly subadults) in Middle and Late-period buildings were frequently not close genetic kin. The choice of burial of genetically unrelated individuals in the same space is reminiscent of practices identified in some Upper Palaeolithic and Mesolithic societies (*62*, *63*) but unknown for the majority of Neolithic contexts from W Eurasia genetically analysed [with the possible exception of contemporaneous Barcın Höyük (*31*)]. Given the dietary similarities between genetically unrelated Çatalhöyük neonates buried in the same building, it is tempting to propose that the widespread fostering of newborns could partly explain the sparse genetic ties among Middle and Late Çatalhöyük co-burials. Such practices could be related to houses becoming economically more independent over time in Çatalhöyük (e.g. grazing sheep in different areas and including larger storage areas) (*64*, *65*), which may have led households to recruit pregnant mothers or newborns from other biological family groups. Fostering could also have served to construct new social kin ties within the community (*30*), perhaps helping maintain egalitarian relationships.

Arguably, the most striking set of observations relates to gender roles. Whether the prominent female adult figurines found in Çatalhöyük indeed represented a “Mother Goddess” cult remains a point of contention (*1*, *2*, *4*). Still, our findings, including the predominance of the matriline connecting intramural burials and the special burial treatment accorded to subadult females, highlight the role of the maternal lineage in Çatalhöyük society. We do not know whether these practices reflected strict matrilineality or gender status differences (i.e. matriarchy), whether they might have been connected with the apparent egalitarian relationships in Çatalhöyük, or whether other Neolithic settlements of the 7th millennium BCE in Anatolia and the Aegean with similar female figurine symbolism had adopted similar practices. Irrespectively, Çatalhöyük social organisation and symbolism stand in stark contrast with the predominantly male-focused societies that emerged in West Eurasia in subsequent millennia.

## Data and code availability

All FASTQ and BAM files produced in this study have been deposited at the European Nucleotide Archive.

Code and scripts used in the analyses will be made available at https://github.com/CompEvoMetu.

## Acknowledgements

We thank Özgür Taşkent, Sabina Cveček, Eva-Maria Geigl, all members of the CompEvo (METU) group and the NEOMATRIX Collective for helpful discussion and suggestions, Reyhan Yaka and Elif Sürer for assistance, Swapan ‘Shop’ Mallick and David Reich for sharing Allen Ancient Genome Diversity Project shotgun-sequenced genomes.

This work was supported by the H2020 ERC Consolidator grant (no. 772390 NEOGENE to M.S), the H2020-WIDESPREAD-05-2020 TWINNING grant (no. 952317 NEOMATRIX to M.S.), TUBITAK of Turkey (no. 117Z229 to M.S.), NSF (no. BCS-1827338 to M.P.). CJK, SDH, and EMJS’s participation in this research benefited from the scientific framework of the University of Bordeaux’s IdEx (Initiative d’Excellence) ‘Investments for the Future’ program / GPR (Grands Programmes de Recherche) ‘Human Past’.

Computations were performed on NEOGENE Clusters at Middle East Technical University.

## Author contributions

