## Supplementary Information for "Female lineages and changing kinship patterns in Neolithic Çatalhöyük"

**Supplementary Figures** **5**
**Methods and Supplementary Results** **28**
1. Laboratory procedures 28
    a. DNA extraction 28
    b. DNA sequencing library preparation and indexing 28
    c. Fourier Transform Infrared (FTIR) Spectroscopy 28
    d. Radiocarbon dating 29
2. Data preprocessing and quality control 29
3. Genomic imputation 31
4. Molecular sex assignment 31
    a. Kx and Ky approaches 31
    b. Comparison of Kx and Rx approaches 34
    c. PMD filtering for sex assignment 34
    d. Turner individual 35
5. Uniparental haplogroup determination 35
6. Genotyping 36
7. Population genetic analysis 37
    a. Çatalhöyük genomes used in population genetic analyses 37
    b. f3-statistics 37
    c. Testing change in pairwise f3 diversity over time 37
    d. Multidimensional Scaling (MDS) 37
    e. Principal Components Analysis (PCA) 38
    f. Admixture modelling with qpAdm 38
    g. Testing change in pairwise haplotype diversity over time 38
    h. f4-statistics 39
    i. qpWave analysis 40
    j. Identifying putative outliers 40
8. Coalescent simulations for interpreting diversity estimates 41
    a. Population genetic simulation framework 41
    b. Population genetic simulation results and discussion 42
9. Detecting IBD segments 46
10. Genetic kinship assignment 47
    a. Relatedness classification cutoffs 47
    b. Autosomal relatedness estimation 47
        i. The first round of kinship estimates 47
        ii. Downsampling experiments to test the effect of imputation on kinship estimates 48
        iii. Close relatedness estimation using ancIBD results 51
        iv. The final round of relatedness assignments 52

v. SNP count threshold based on false positives 54
c. Relatedness estimation based on X chromosome 54
d. Estimating variation in kinship estimates 54
11. Runs of Homozygosity (ROH) 56
12. Genealogy simulations for measuring inbreeding under random mating 56
a. Simulation framework 56
b. Implementation for comparison with observed data 57
13. Uniparental haplogroup homogeneity / diversity analyses 58
a. Determining major haplogroups 58
b. Haplogroup homogeneity calculations 59
c. Haplogroup assignment in the Gurgy dataset 59
14. Simulations of matrilocal, patrilocal and mixed residence 59
a. Simulation of founders 59
b. Simulation of pedigrees 60
c. Residence type simulations for haplogroup diversity 61
d. Residence type simulations for autosome vs chrX diversity 62
e. Modelling Çatalhöyük residence dynamics using simulation results 62
15. Burial and diet analyses 63
a. DNA preservation differences 63
b. Burial objects 63
c. Dietary isotopes 63
16. Simulations of within-building co-burial relatedness 64
17. Metagenomic analysis 66
a. Microbial screening 66
b. Testing differences in bone preservation-associated microbes using random forests 67
c. Testing aerobic versus anaerobic microbe abundance differences 70
**References** **71**

### Supplementary Figures

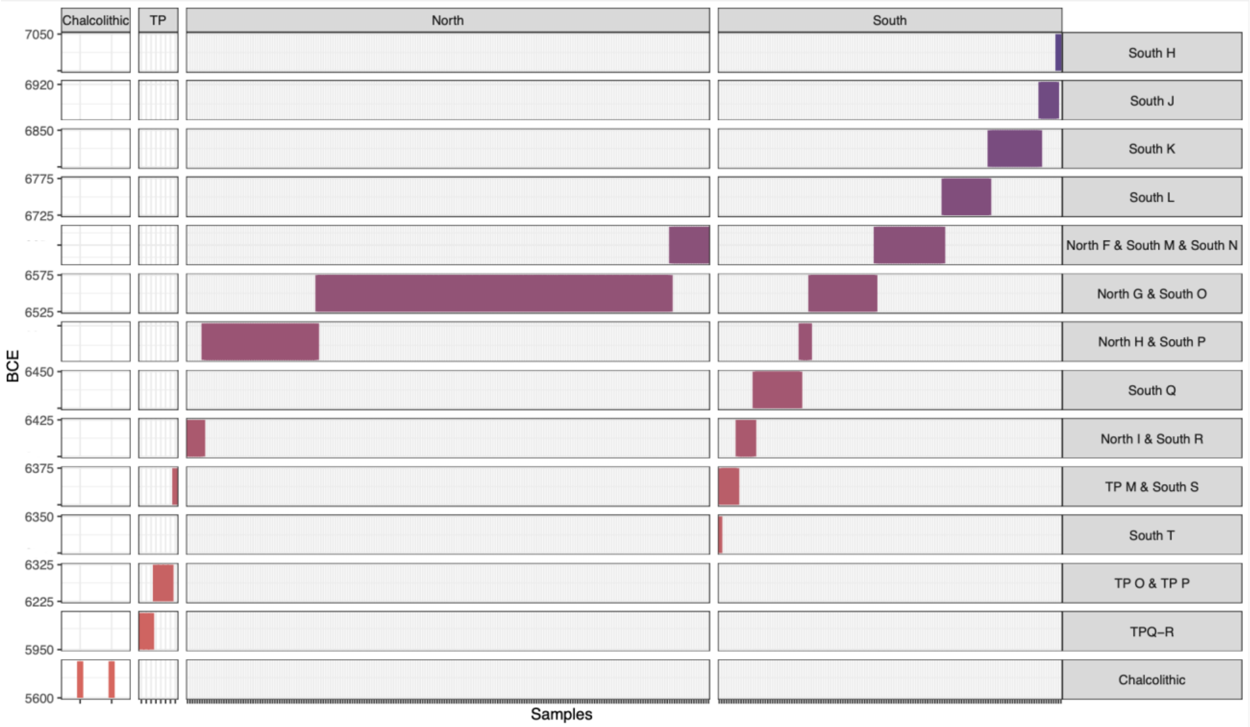


**Figure S1:** Temporal and regional distribution of all 411 genetically screened Çatalhöyük individuals. Each line represents an individual assigned to a specific archaeological level, shown on the left, and the date range of the levels (*1*), shown on the right (see Table S1). Note that the two West Mound (Chalcolithic) genomes have been dated to c.5900-5800 calBCE.

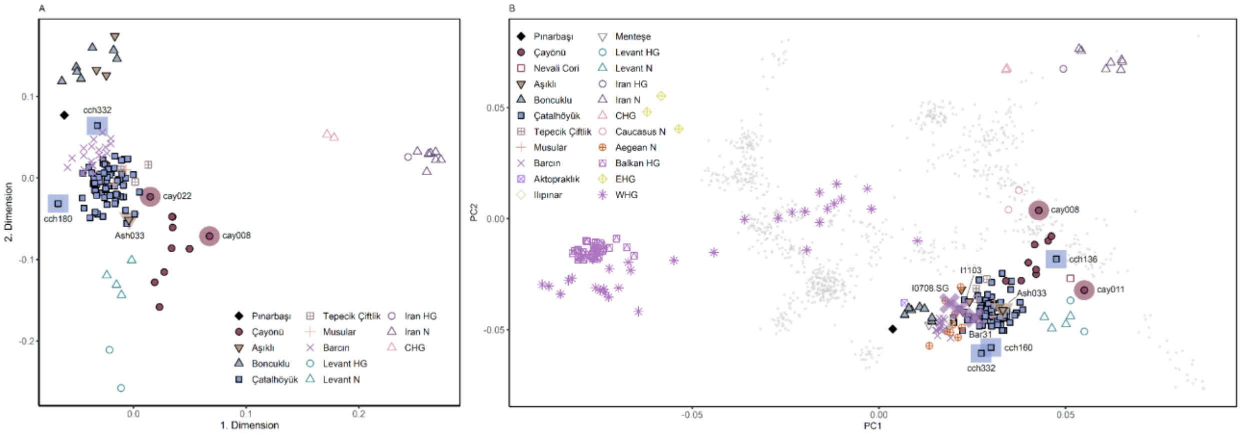


**Figure S2:** Genetic clustering of Çatalhöyük genomes with Upper Paleolithic and Early Holocene genomes from Southwest Asia. **A)** MDS and **B)** Principal Component Analysis (PCA) of ancient and modern-day genomes. The PCs were calculated using Western Eurasian modern-day genomes from the Human Origins (HO) dataset, and ancient Upper Paleolithic and Early Holocene genomes from Southwest Asia (data produced here and published genomes) were projected on this PC space. In both MDS and PCA outlier individuals are labeled and marked with larger semi-transparent symbols.

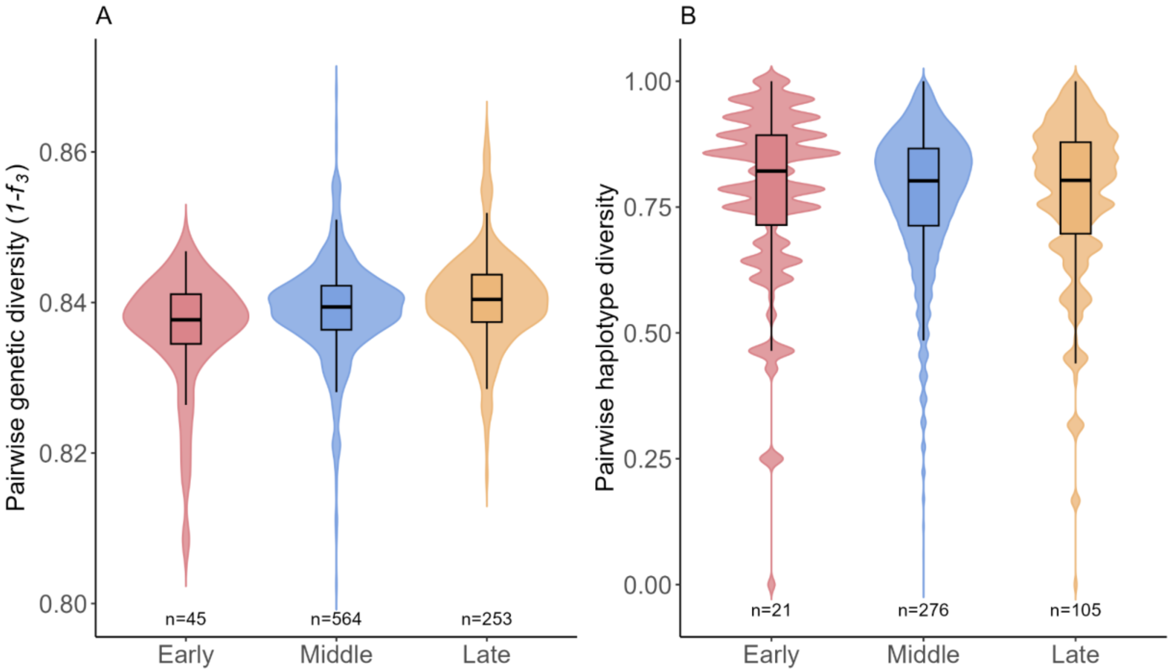


**Figure S3:** Genetic diversity in three main Çatalhöyük periods. Allelic diversity was estimated using **A)** pairwise outgroup $f_3$-based diversity [i.e. $f_3$*(Outgroup; ÇatalhöyükInd1, ÇatalhöyükInd2)*] and **B)** haplotype-based diversity, by comparing all available genomes assigned to that period. The latter was calculated using imputed genomes (see Supplementary Methods section “Testing change in pairwise haplotype diversity over time”). Numbers at the bottom of the graphs show number of pairs in that period .

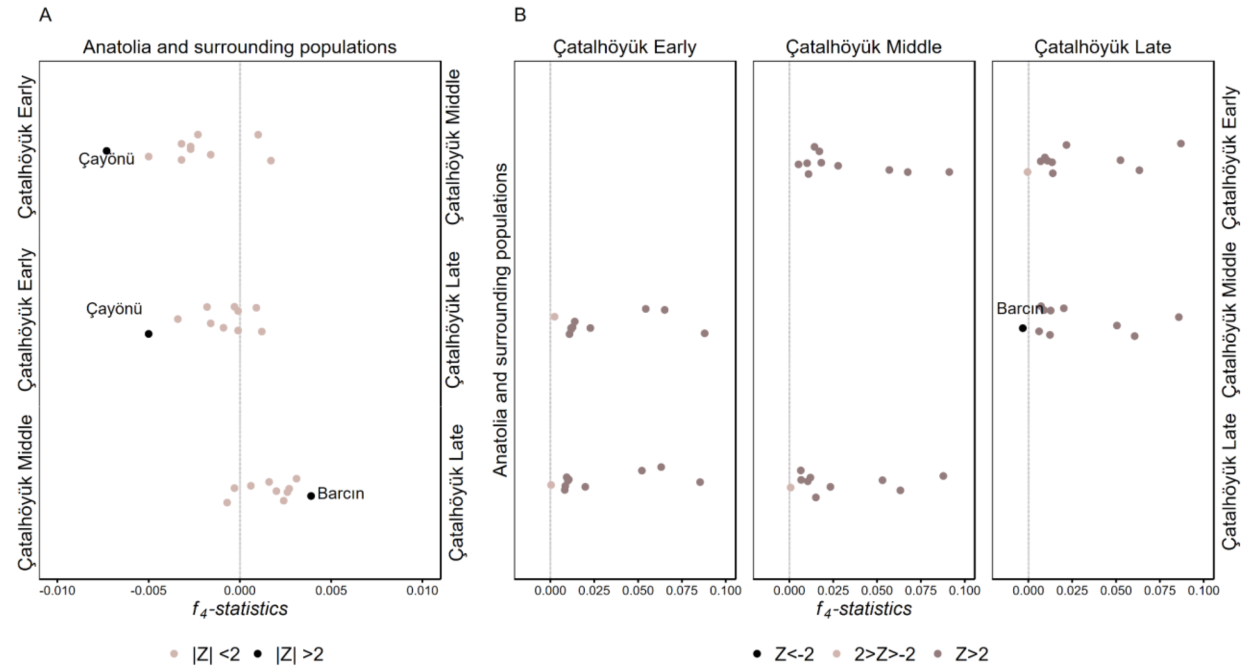


**Figure S4**: Genetic affinities between Çatalhöyük periods and other Anatolian Neolithic or surrounding regional populations measured in $f_4$-statistics. The statistics were computed in the form of **A)** *$f_4$(Outgroup, AnatSurrPop; ÇatalhöyükPeriod1, ÇatalhöyükPeriod2)* and **B)** *$f_4$(Outgroup, ÇatalhöyükPeriod1; AnatSurrPop, ÇatalhöyükPeriod2)*. The Anatolia Neolithic group comprises genomes from Aşıklı Höyük, Boncuklu, Çayönü, Musular, Tepecik Çiftlik and Barcın, meanwhile surrounding populations comprise Iran Neolithic, Levant Neolithic, Balkan hunter-gatherer (HG) and Caucasus hunter-gatherer (CHG). Each group is tested independently, represented by a single point in the graph. Comparisons which were nominally significant with |Z|>2 are indicated with their site names and as light coloured points.

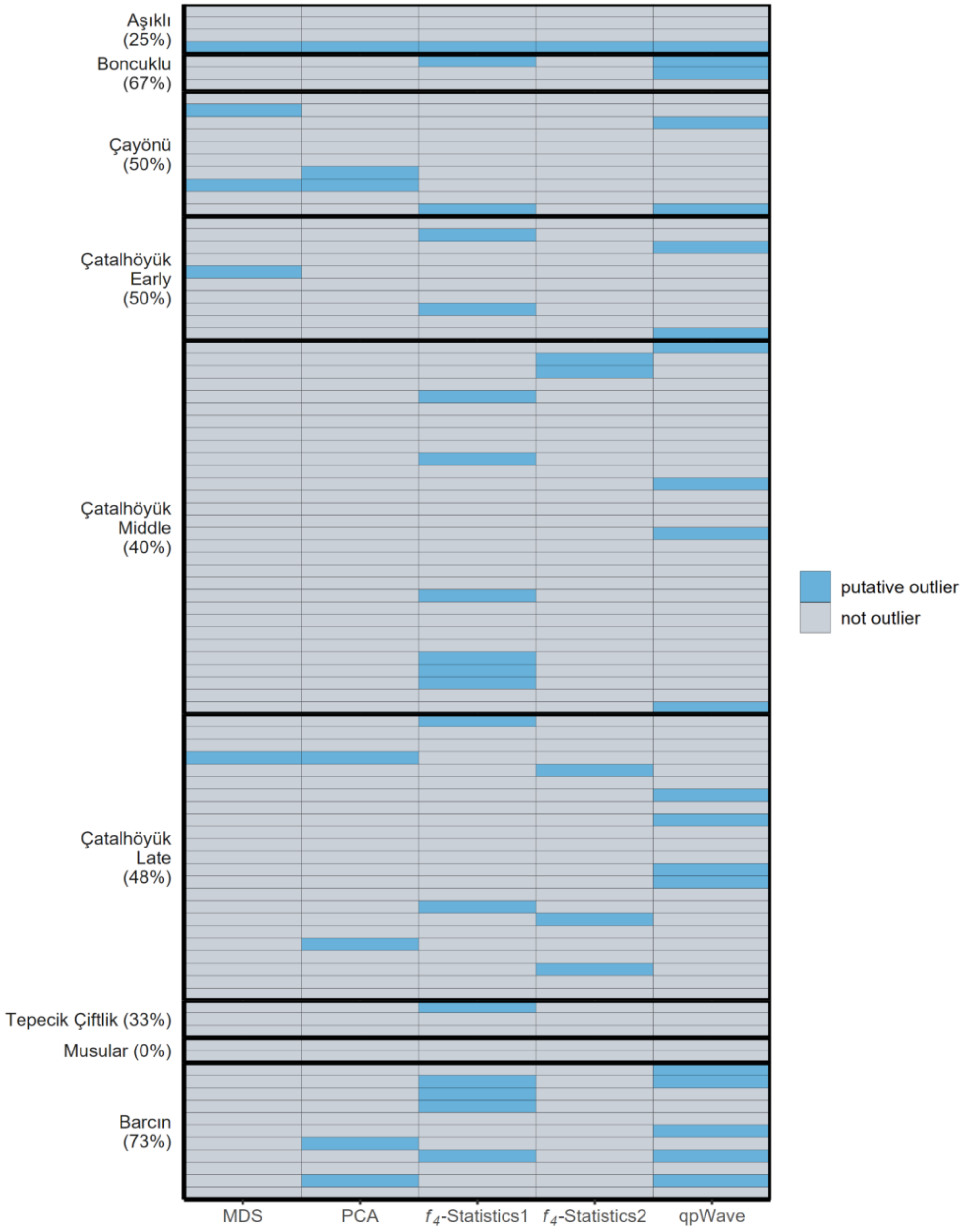


**Figure S5:** Screening for genetic outliers in Çatalhöyük and six other Anatolian Neolithic genomes. The figure summarises the results from Table S7, where we used two clustering methods (MDS and PCA) and three formal tests (two $f_4$-statistics and qpWave) to identify putative genetic outlier individuals (see Supplementary Information section 'Population genetic analysis'). Each row indicates a tested individual, and blue cells show positive results for a test. Çatalhöyük samples are grouped by their periods. The percentages of outliers per population are shown in parentheses.

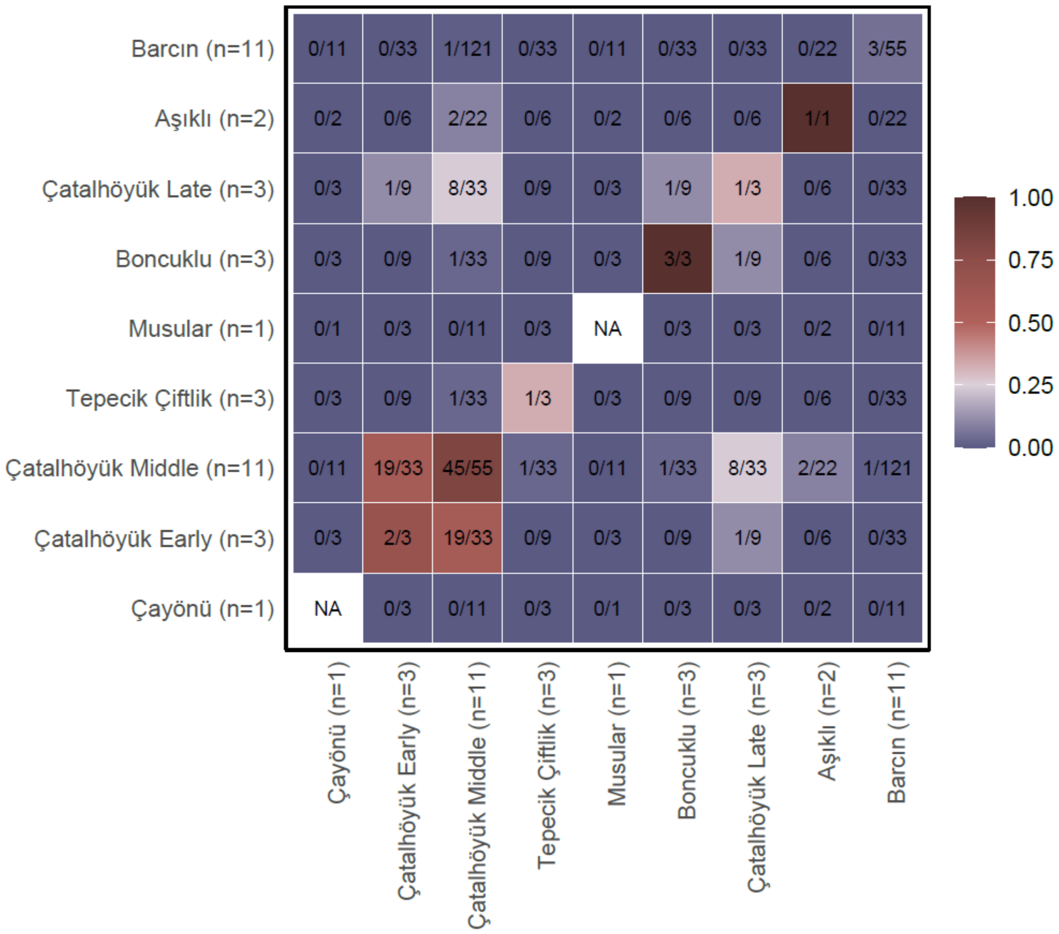


**Figure S6:** Heatmaps showing IBD-sharing among imputed Anatolian Neolithic genomes based on ancIBD results. The color coding indicates the proportion of sharing, as shown as a fraction within each cell. For instance, the number 0/11 in the top left cell indicates that Çayönü, represented by imputed 1 genome, was compared with 11 imputed Barcın late genomes. Out of 11 comparisons, none of the pairs shared any IBD segments of length > 12 cM.

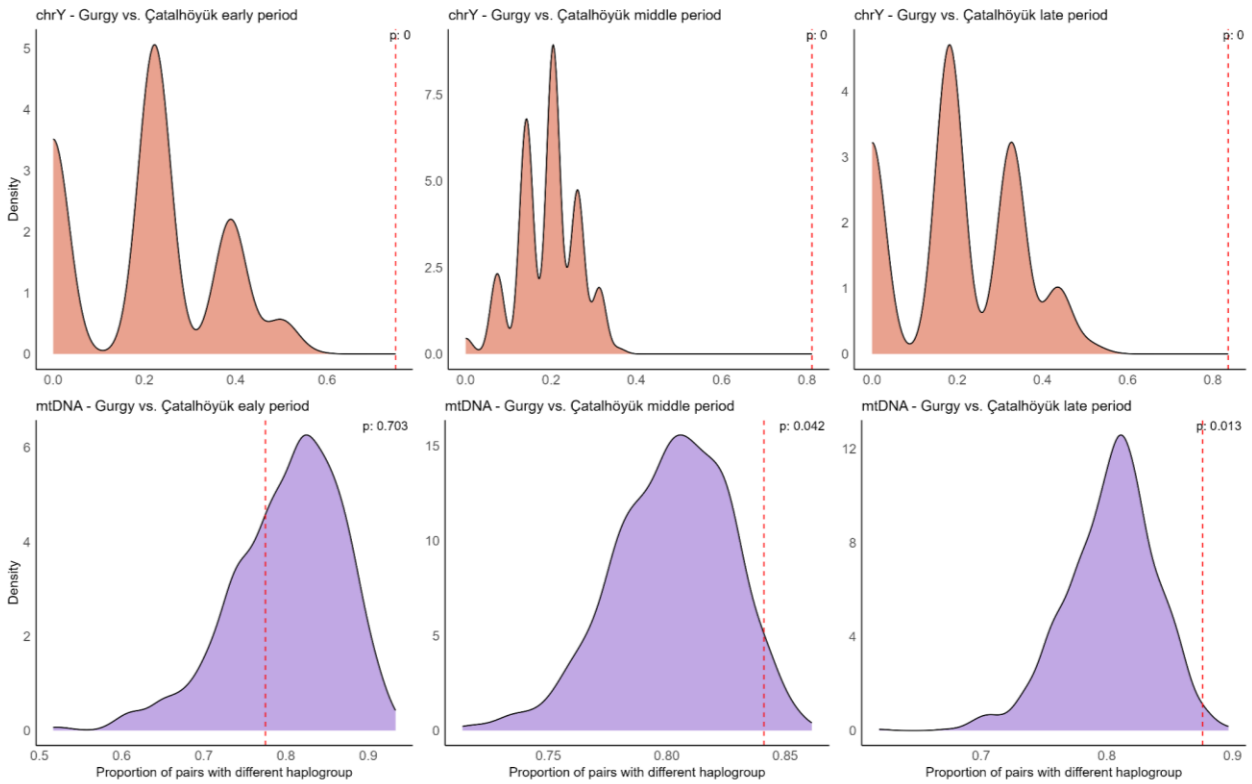


**Figure S7:** Random subsampling tests for Çatalhöyük-Gurgy uniparental haplogroup diversity differences (**Figure 2E**). Top panels: chrY diversity. Bottom panels: chrY diversity. Left to right: Early, Middle, and Late period Çatalhöyük. We randomly subsampled the same number of individuals as in each of the Çatalhöyük periods from the much larger Gurgy set, 1000 times, and calculated the proportion of pairs with different haplogroups, shown by the density plots. The observed proportions for each Çatalhöyük period is shown in vertical red dashed lines. The p-values are also indicated on the top right in each panel. All periods of Çatalhöyük diverge from Gurgy in having an incomparably higher chrY diversity (top panels). The Middle and Late periods of Çatalhöyük also differ from Gurgy in having slightly higher mtDNA diversity, which may be related to the fact that the latter two periods comprise fewer close relatives, hence their higher diversity. We note that the high frequency of H subclades in Gurgy and the fact that we cut the tree at a high level might partly explain high mtDNA homozygosity in Gurgy.

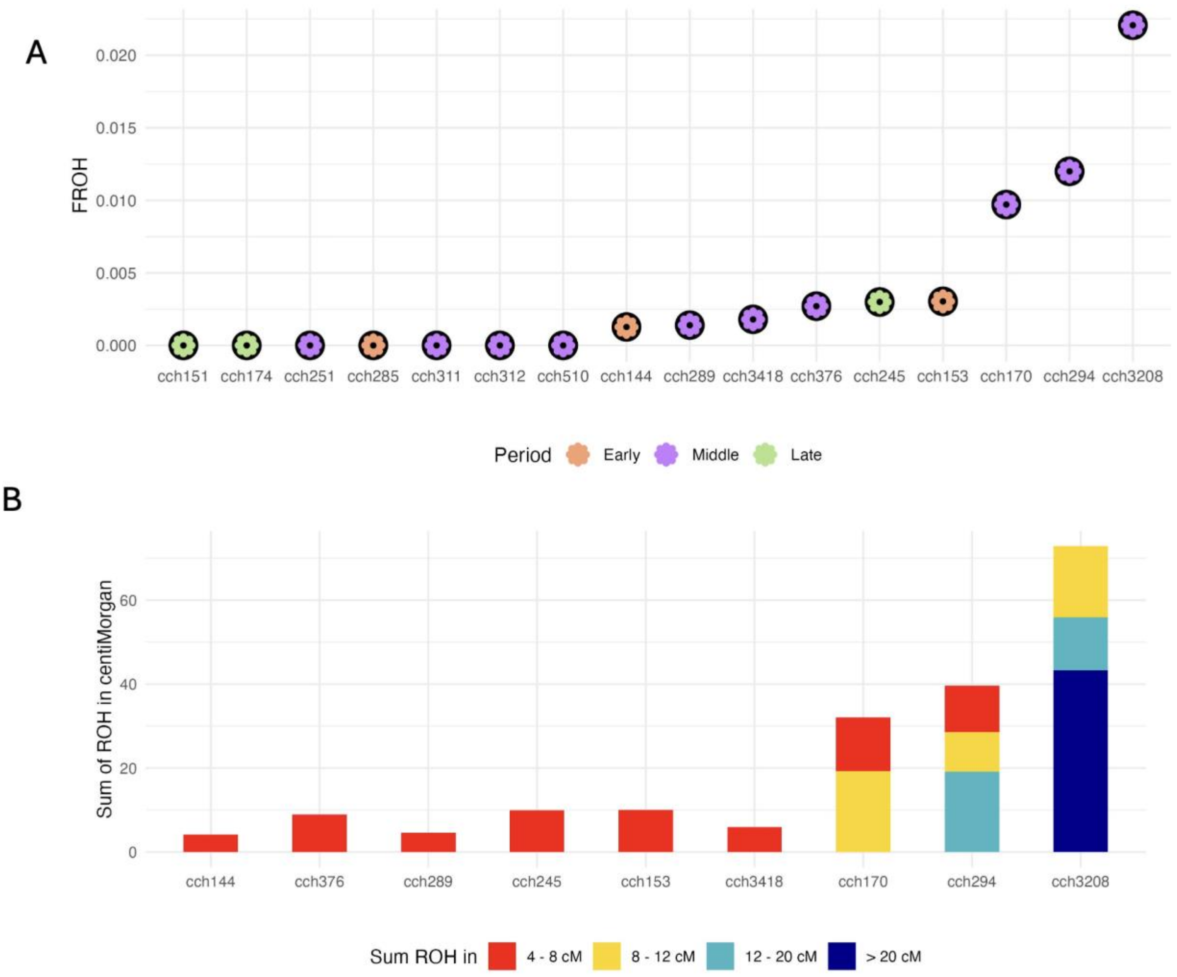


**Figure S8:** Consanguinity as measured by runs of homozygosity (ROH) in Çatalhöyük. **A)** The inbreeding coefficient ($F_{ROH}$) estimated based on hapROH results for 16 individuals with sufficient genomic coverage (>0.3x). **B)** The distribution of segments for genomes with minimum one ROH segment. Note that hapROH only identifies ROH >4cM.

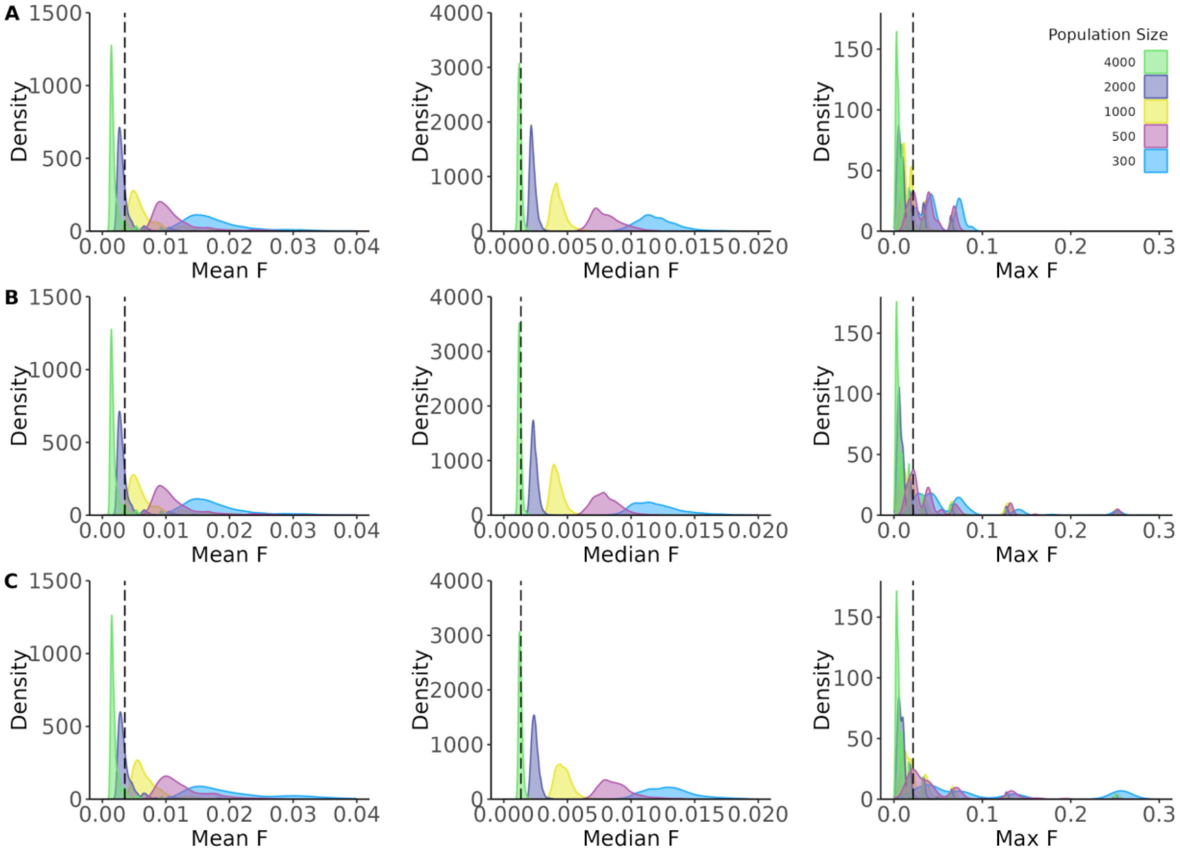


**Figure S9:** Distribution of mean, median, and maximum of simulated pedigree-based inbreeding coefficients ($F_{ped}$) under different scenarios and mean $F_{ROH}$ estimates for 16 genomes. The simulations were performed under three different conditions: **A)** sibling unions are not allowed, and the sex ratio in the breeding population is 1:1; **B)** sibling unions are allowed, and the sex ratio is 1:1; **C)** sibling unions are allowed, and there is reproductive skew with ~1:2 males vs females (see **Table S18**). We ran the algorithm 5 times for each distribution with the same parameters, taking 1000 subsets with n=16 individuals each time randomly, and calculating corresponding statistics. The dashed line represents corresponding statistics for observed $F_{ROH}$ levels in 16 Çatahöyük individuals from all periods.

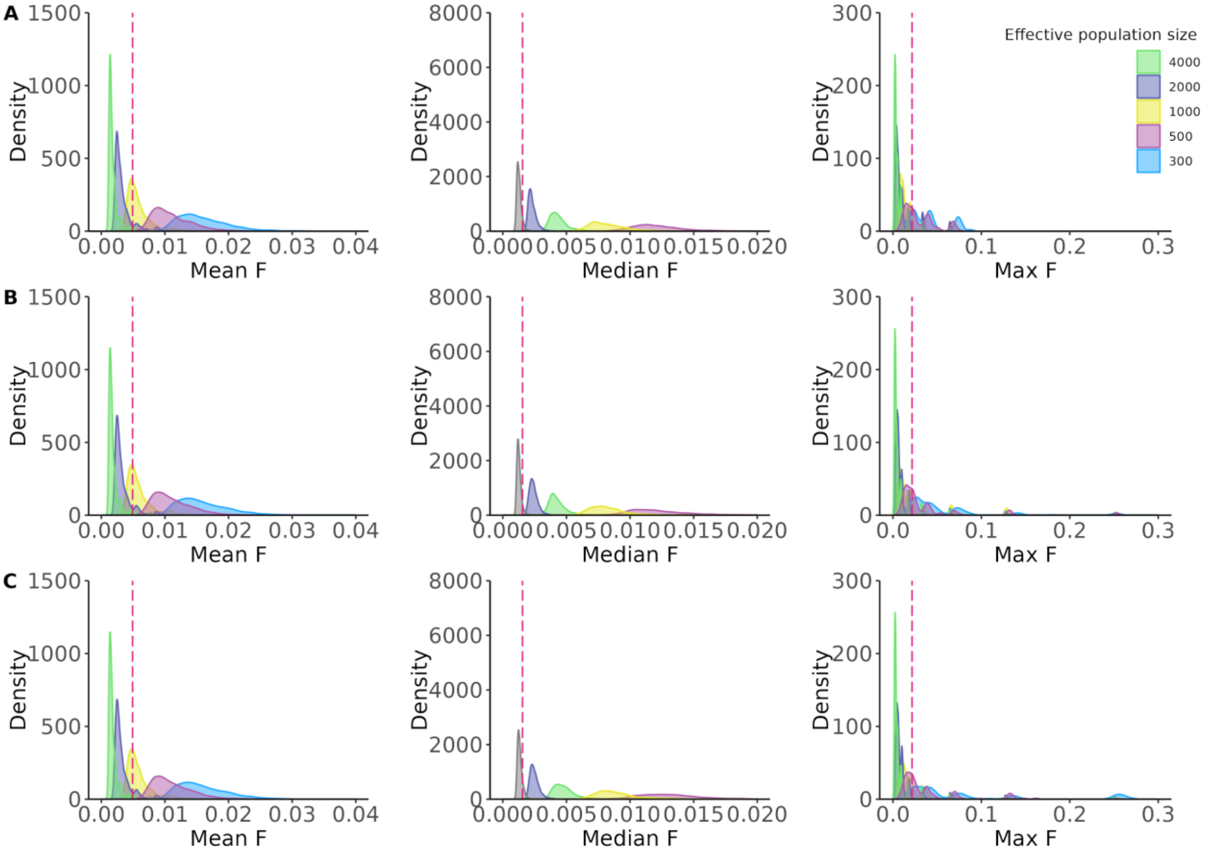


**Figure S10:** Distribution of mean, median, and maximum of simulated inbreeding coefficients ($F_{ped}$) under different scenarios and mean $F_{ROH}$ estimates for 10 genomes. The simulations were performed under three different conditions: **A)** sibling unions are not allowed, and the sex ratio in the breeding population is 1:1; **B)** sibling unions are allowed, and the sex ratio is 1:1; **C)** sibling unions are allowed, and there is reproductive skew with ~1:2 males vs females (see **Table S18**). We ran the algorithm 5 times for each distribution with the same parameters, taking 1000 subsets with n=10 individuals each time randomly, and calculating corresponding statistics. The dashed line represents corresponding statistics for observed $F_{ROH}$ levels in 10 Çatahöyük genomes from the Middle period.

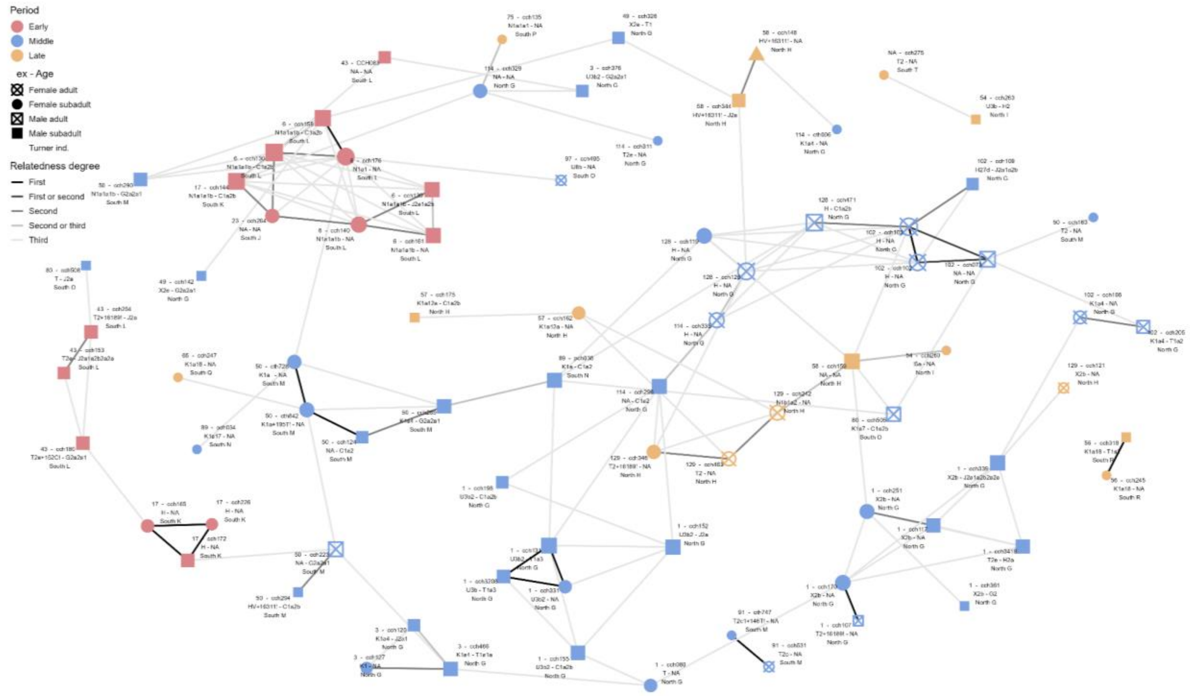


**Figure S11:** A network of estimated genetic kin relationships among 80 Çatalhöyük individuals with common SNPs >3000 (see section ‘Genetic kinship assignment’).

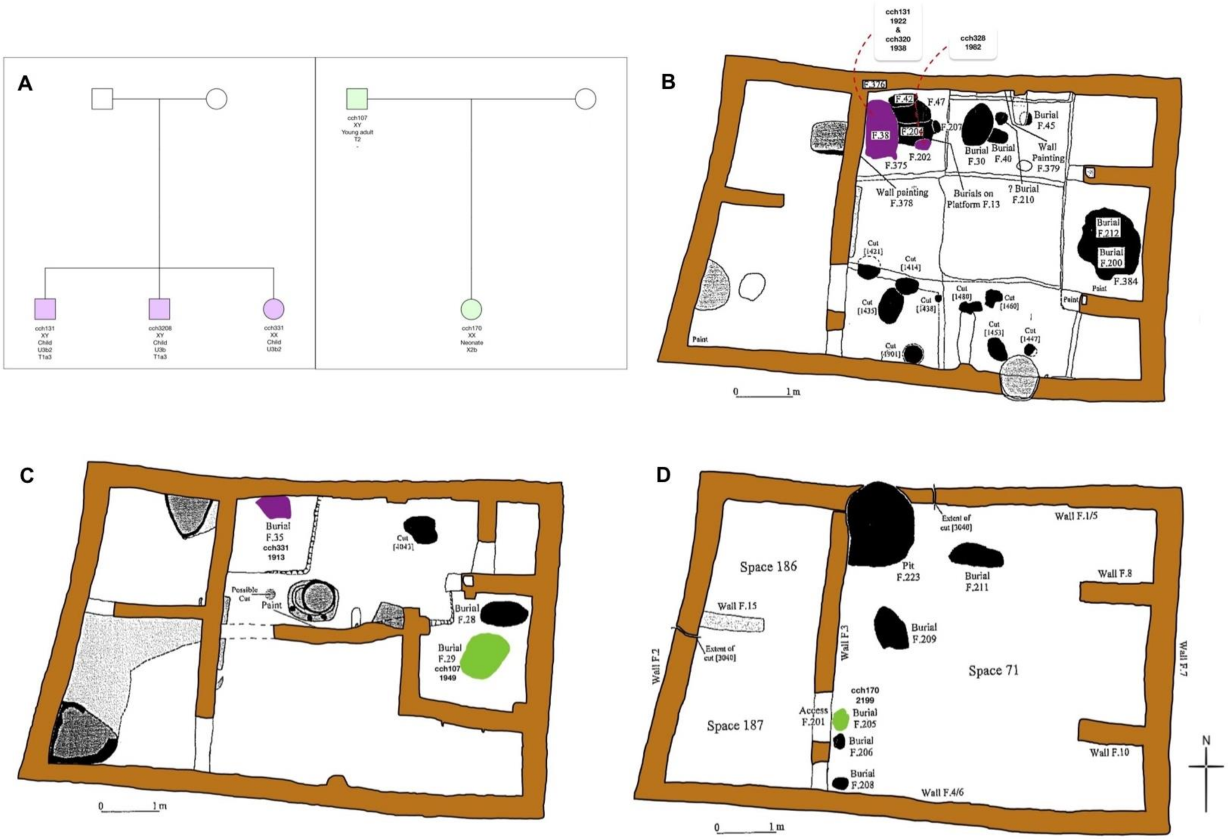


**Figure S12A:** Possible pedigrees for genetic kin pairs co-buried in Building 1. **A)** Pedigrees of identified first-degree related pairs from two genetic families, shown here as purple and green. Circles show female and squares show male individuals. In the pedigrees, the lab ID, sex, approximate age at death, mitochondrial haplogroup, and Y chromosomal haplogroup, of each genetically studied individual is indicated, respectively. Plans of building subphases **B)** B1.2B, **C)** B1.2C, and **D)** B1.1B showing burial locations. On the plan, labels show lab ID, excavation ID (unit number), and feature number of each burial, respectively.

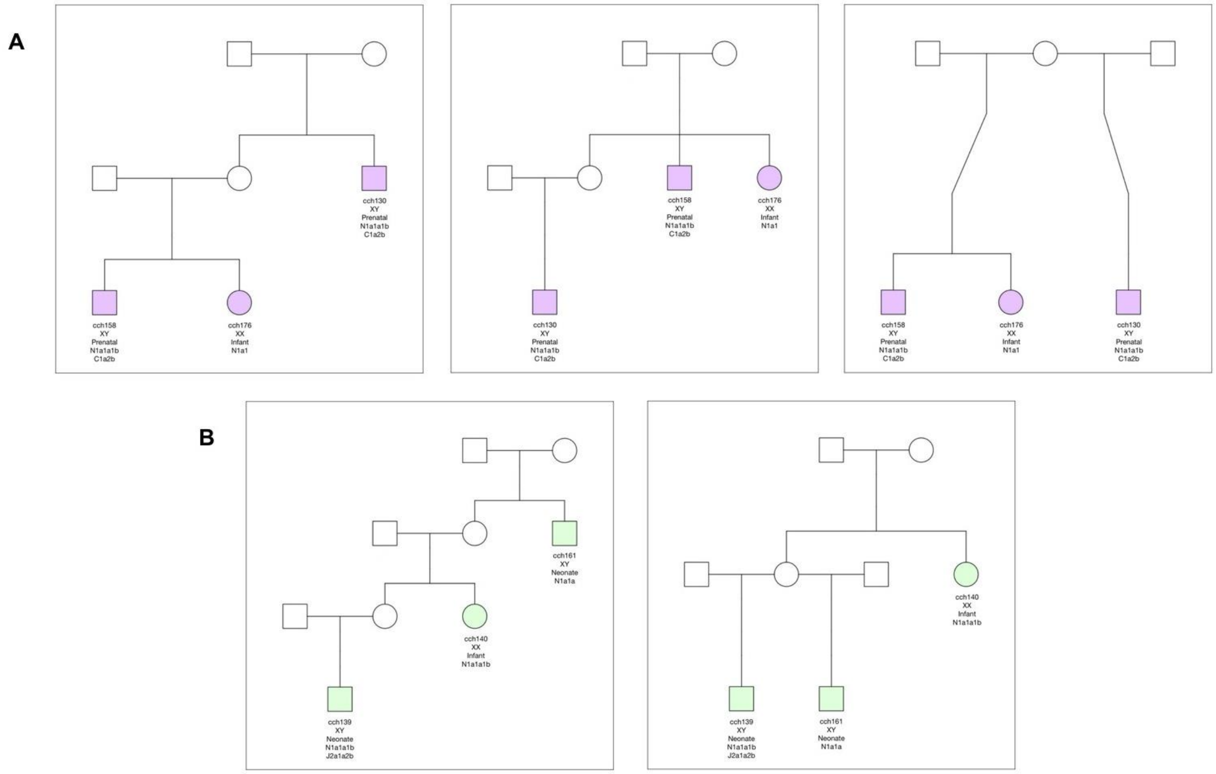


**Figure S12B:** Possible pedigrees for genetic kin pairs co-buried in Building 6. The panels show alternative possible pedigree configurations of two genetic families identified (shown here as purple and green). Circles show female and squares show male individuals. Colors indicate individuals belonging to different families. In the pedigrees, the lab ID, sex, approximate age at death, mitochondrial haplogroup, and Y chromosomal haplogroup, of each genetically studied individual is indicated, respectively. The building plan was not available.

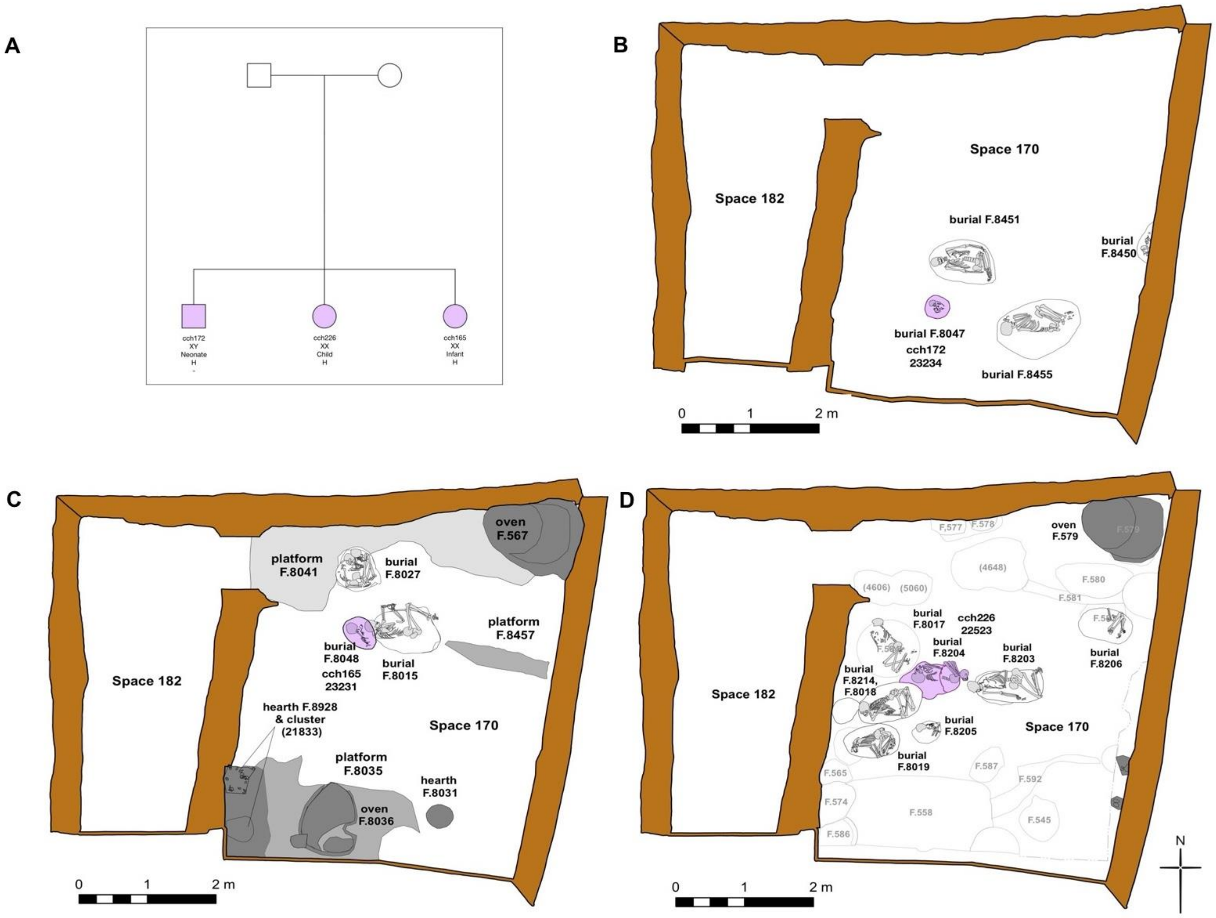


**Figure S12C:** Possible pedigrees for genetic kin pairs co-buried in Building 17. **A)** The pedigrees of three burials identified as a genetic family. Circles show female and squares show male individuals. In the pedigrees, the lab ID, sex, approximate age at death, mitochondrial haplogroup, and Y chromosomal haplogroup, of each genetically studied individual is indicated, respectively. (-) indicates no reliable haplogroup information. Plans of **B)** phase 1, **C)** phase 2.2, **D)** phase 2.3/E. Pink color indicates individuals belonging to the same family. In pedigrees, labels show lab ID, sex, age at death, mitochondrial haplogroup, and Y chromosomal haplogroup accordingly. On the plan, labels show lab ID, excavation ID (unit number), and feature number of each burial, respectively.

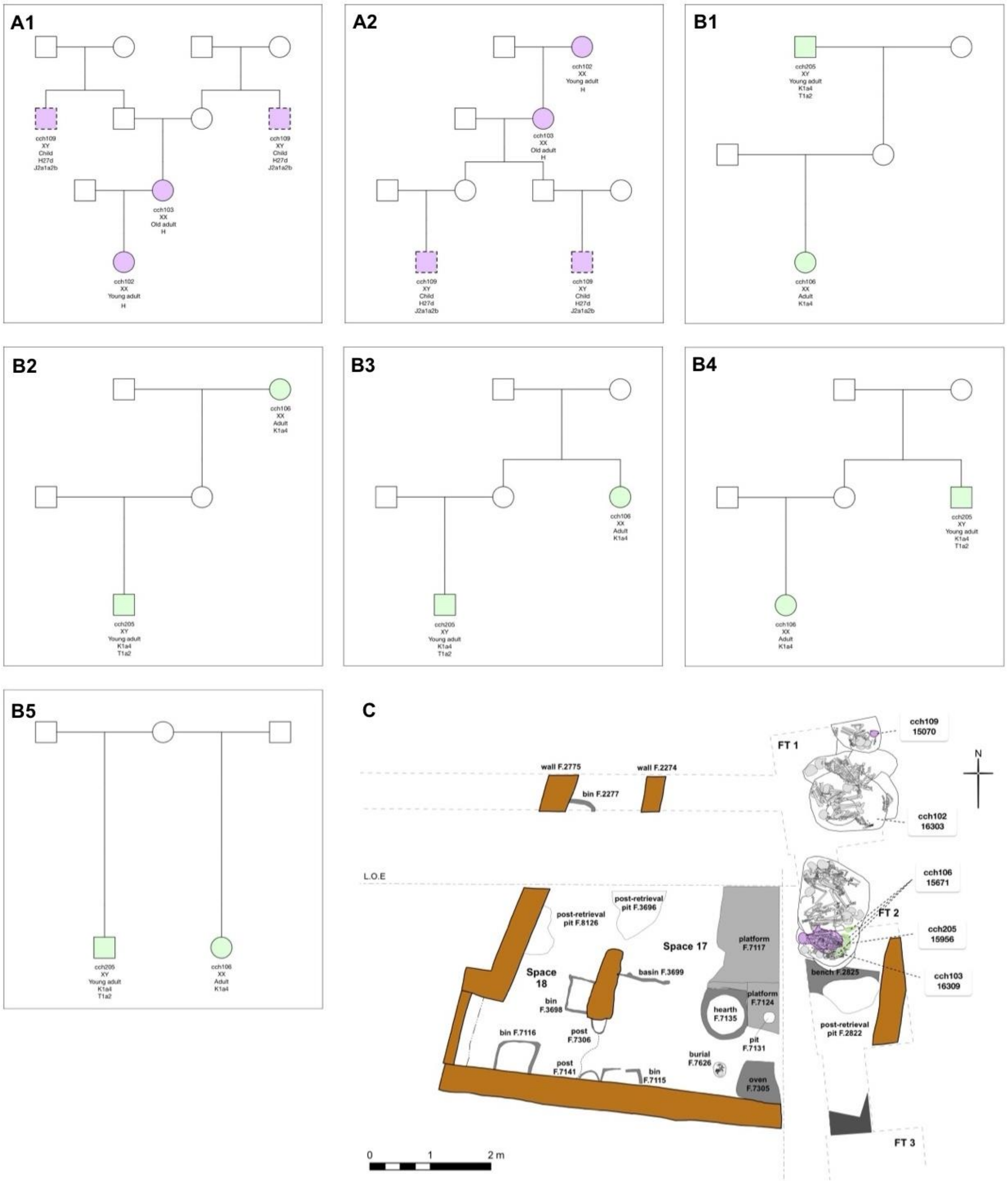


**Figure S12D:** Possible pedigrees for genetic kin pairs co-buried in Building 102. **A-B)** The panels show alternative possible pedigree configurations of two identified genetic families (purple and green). Circles show female and squares show male individuals. In the pedigrees, the lab ID, sex, approximate age at death, mitochondrial haplogroup, and Y chromosomal haplogroup, of each genetically studied individual is indicated, respectively. (-) indicates no reliable haplogroup information. In panel A, the dashed square shows alternative genetic kin ties for the cch109 individual. **C)** Plan of phase C of the building. On the plan, labels show lab ID, excavation ID (unit number), and feature number of each burial, respectively.

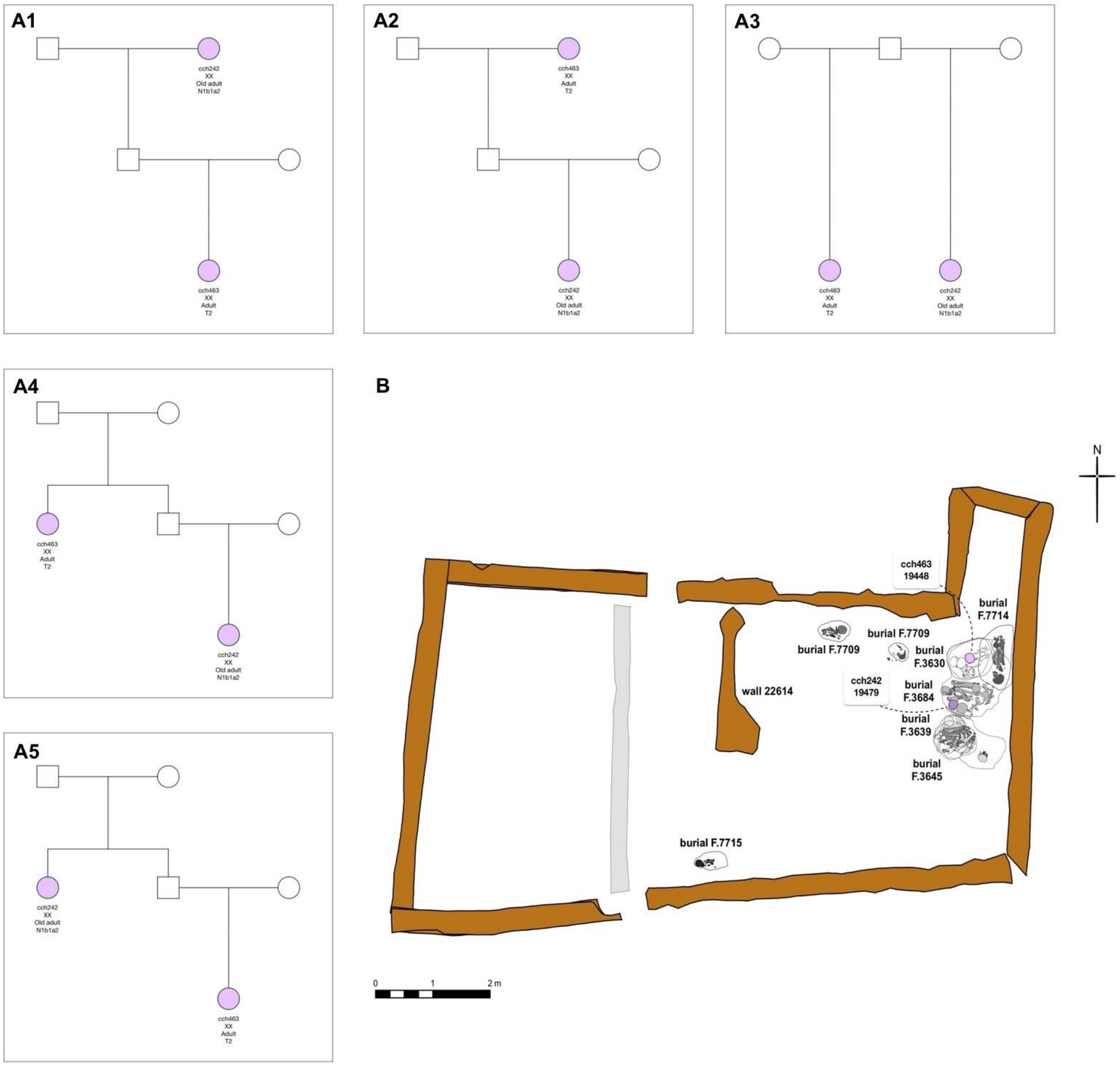


**Figure S12E:** Possible pedigrees for genetic kin pairs co-buried in Building 129. **A)** The panels show five alternative possible pedigree configurations of an identified genetic family. Circles show female and squares show male individuals. In the pedigrees, the lab ID, sex, approximate age at death, mitochondrial haplogroup, and Y chromosomal haplogroup, of each genetically studied individual is indicated, respectively. (-) indicates no reliable haplogroup information. **B)** Plan of Building 129. On the plan, labels show lab ID, excavation ID (unit number), and feature number of each burial, respectively.

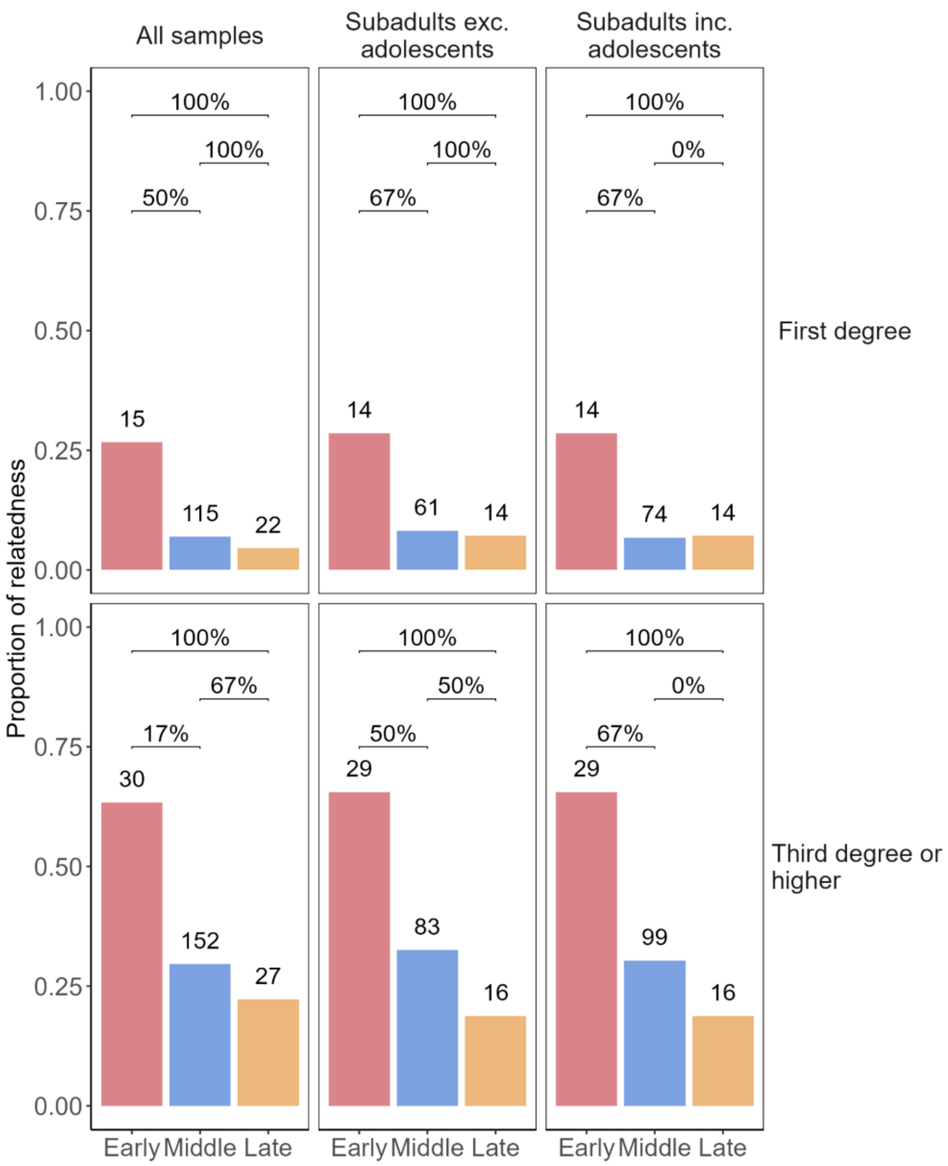


**Figure S13:** Proportion of genetic kin among co-buried pairs in each Çatalhöyük period. The same analysis was performed by including two different cutoffs for estimated genetic kin relations, shown in rows: "third-degree or higher": unambiguous third-degree and closer relationships; "first-degree": only first-degree relationships. The columns show different age groups included, from left to right: all samples, subadults excluding adolescents, and subadults including adolescents. The percentages above the horizontal bars show the percentage of Monte-Carlo simulations where the null hypothesis of no difference between a pair of periods was rejected out of 18 scenarios per comparison. The 18 different scenarios involved using fixed or variable family sizes (2-6) assigned to each building and using different age categories. The number reported in the main text is the the results of simulations of all samples with genetic kin "third degree or higher".

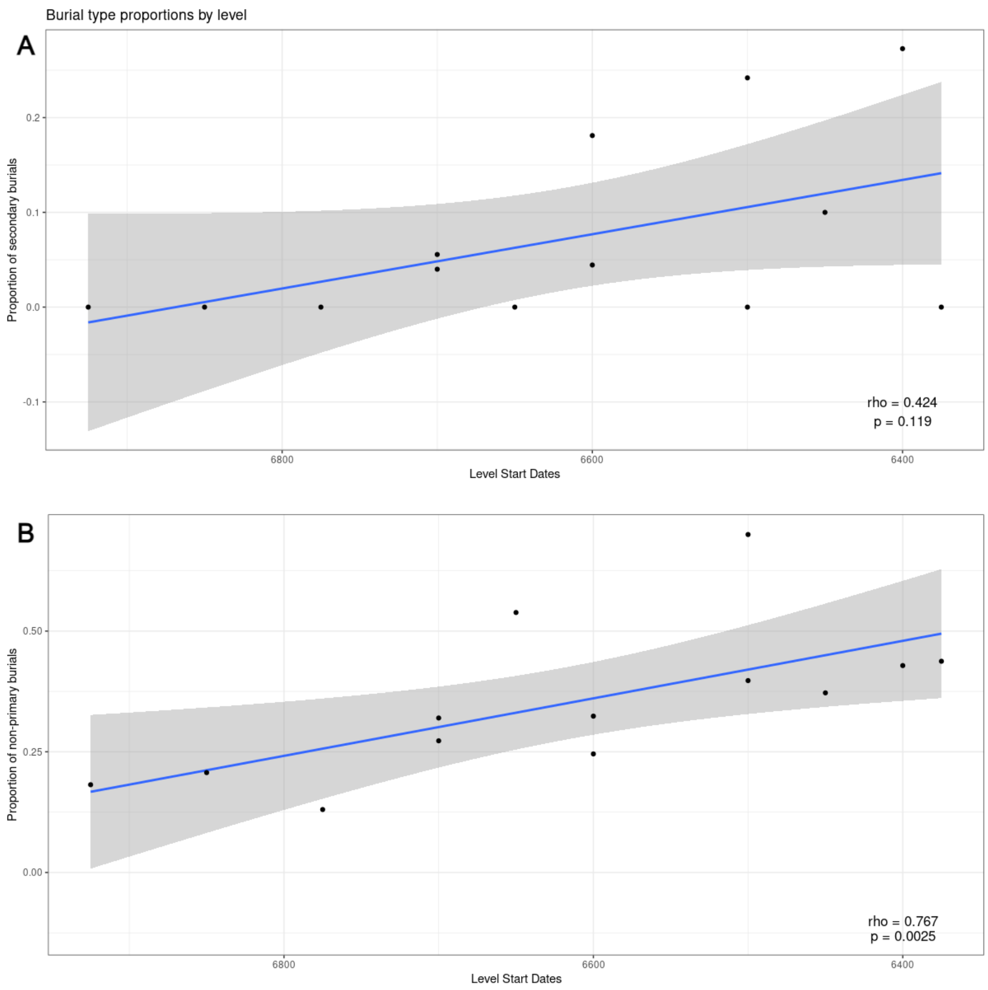


**Figure S14:** Increasing frequency of non-primary burials over time. Panel **A** shows the frequency of secondary burials among all primary and secondary burials (excluding tertiary burials), while Panel **B** shows the frequency of all non-primary burials among all burials within Çatalhöyük North and South levels (**Table S1**). Each point represents the proportion in one level. The change over time was measured using Spearman correlation; the coefficient and p-values are indicated within the panels.

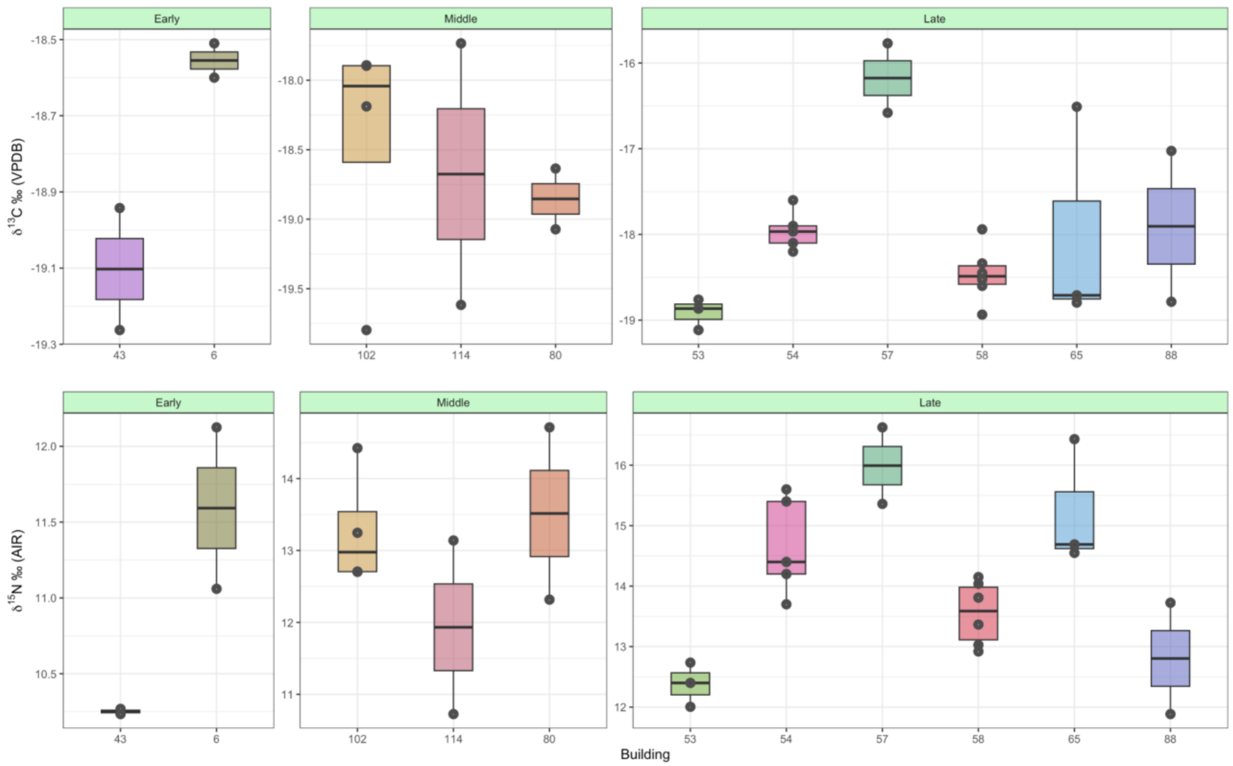


**Figure S15:** Distribution of stable carbon ($\delta^{13}C$) (upper panel) and nitrogen ($\delta^{15}N$) (lower panel) values measured in neonates from the Late period. The majority of neonates belong to the Late period and solely these Late period neonates show statistically significant differences among buildings (Kruskal-Wallis test $p<0.02$) (see Supplementary Information section "Dietary isotopes").

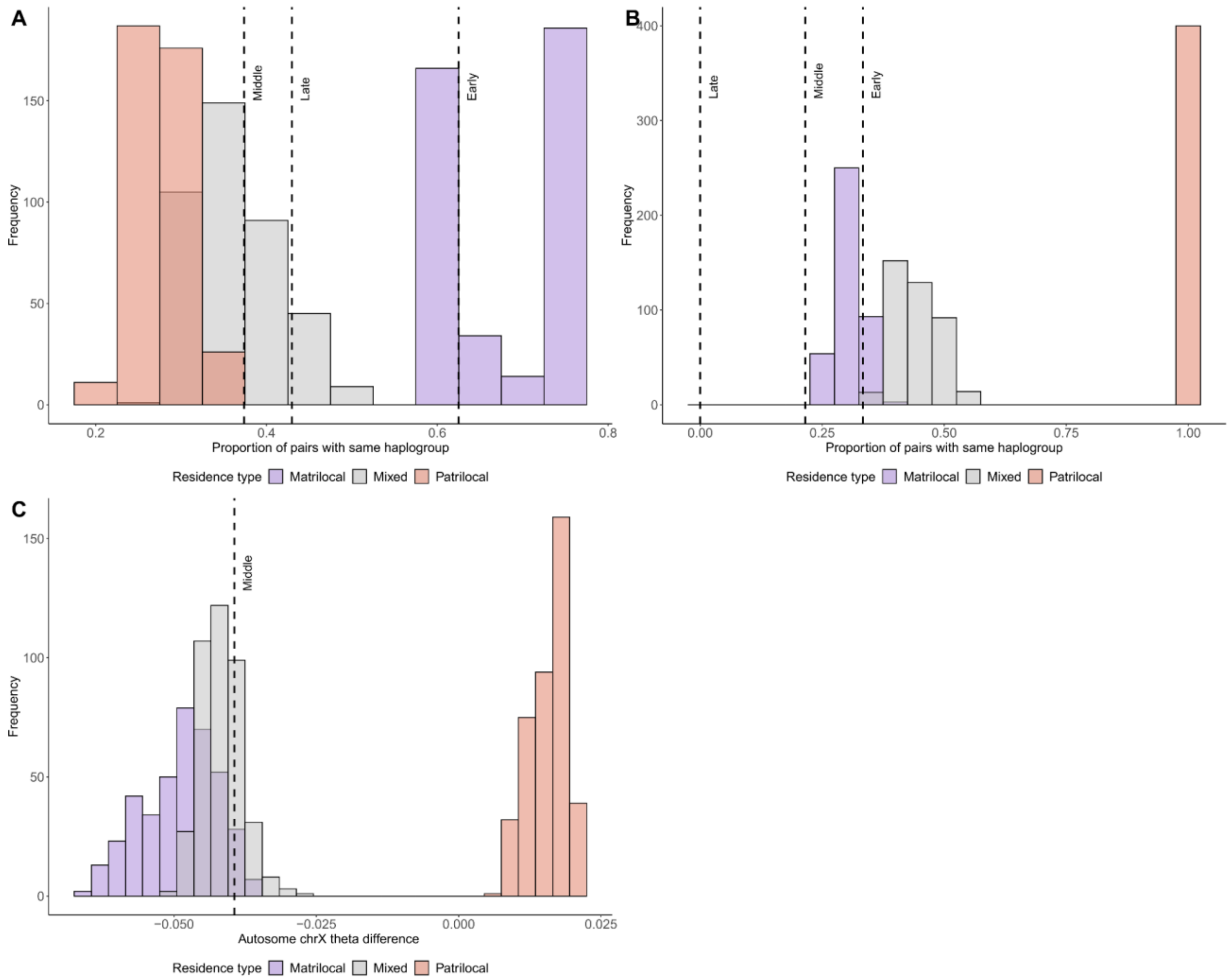


**Figure S16:** Modelling sex-biased genetic connections within Çatalhöyük buildings. We simulated genotype and haplogroup data under three scenarios: matrilocal residence where only males move among buildings, patrilocal residence where only females move among buildings, and the mixed model where both males and females move. We then measured **A)** chrY haplogroup homogeneity and **B)** mtDNA haplogroup homogeneity for pairs of individuals buried in the same building in the simulations. The x-axis shows the proportion of pairs with the same haplogroup. The observed mean homogeneity values are shown as dashed lines; these were calculated for co-buried pairs within Çatalhöyük buildings, separately for each of the three periods. **C)** We further calculated autosomal and X chromosomal (chrX) kinship coefficients (theta) within buildings. The x-axis shows the difference between autosomal and chrX thetas in simulated residence scenarios. The observed mean value for the Middle period is shown as dashed line.

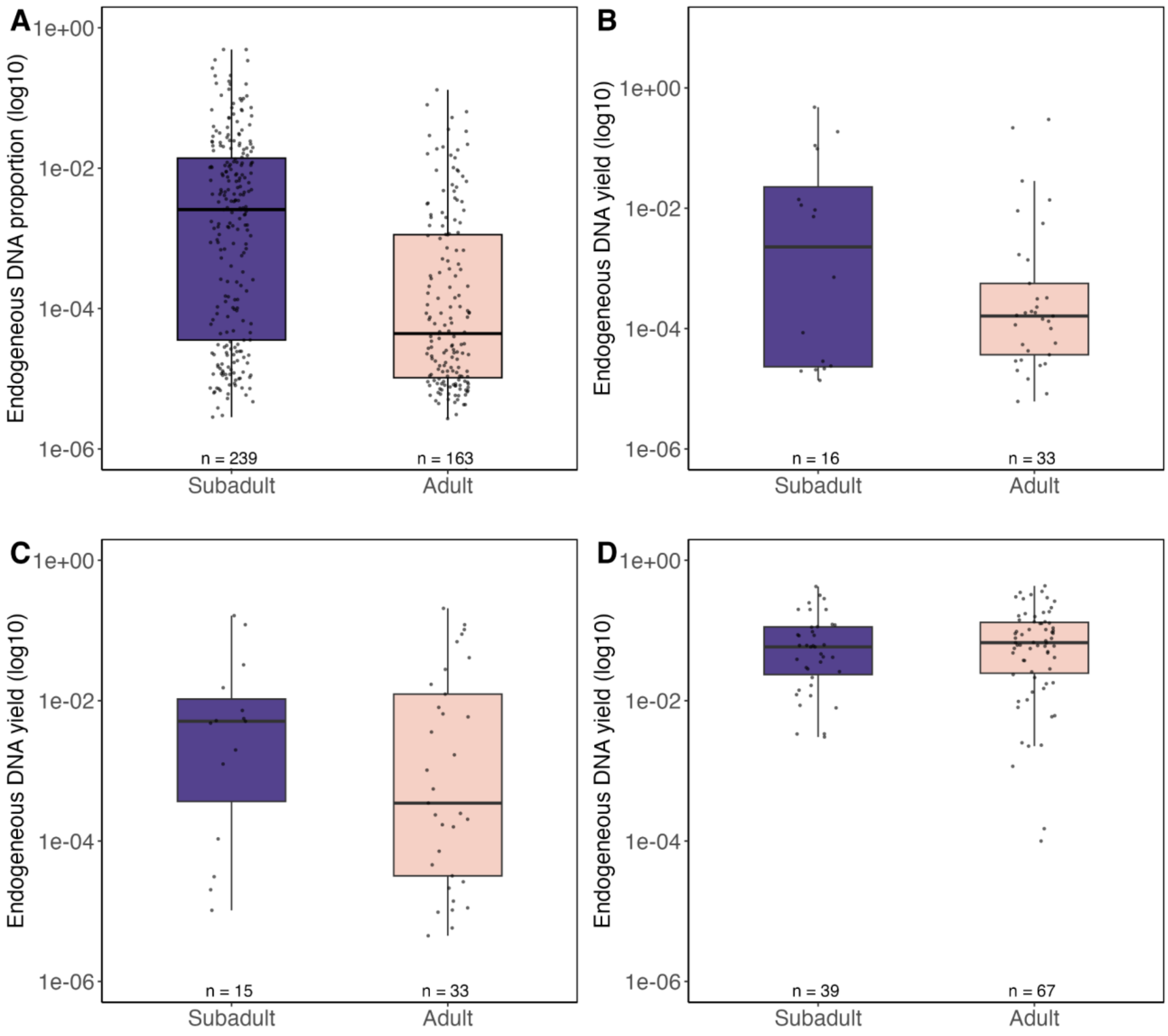


**Figure S17:** Endogenous aDNA proportions among adults and subadults compared in other Neolithic West Eurasian villages; **A)** Çatalhöyük from C Anatolia, **B)** Aşıklı Höyük from C Anatolia, **C)** Çayönü from U Mesopotamia, **D)** Gurgy from France. None of the comparisons other than Çatalhöyük were significant in Mann-Whitney U tests (p>0.05).

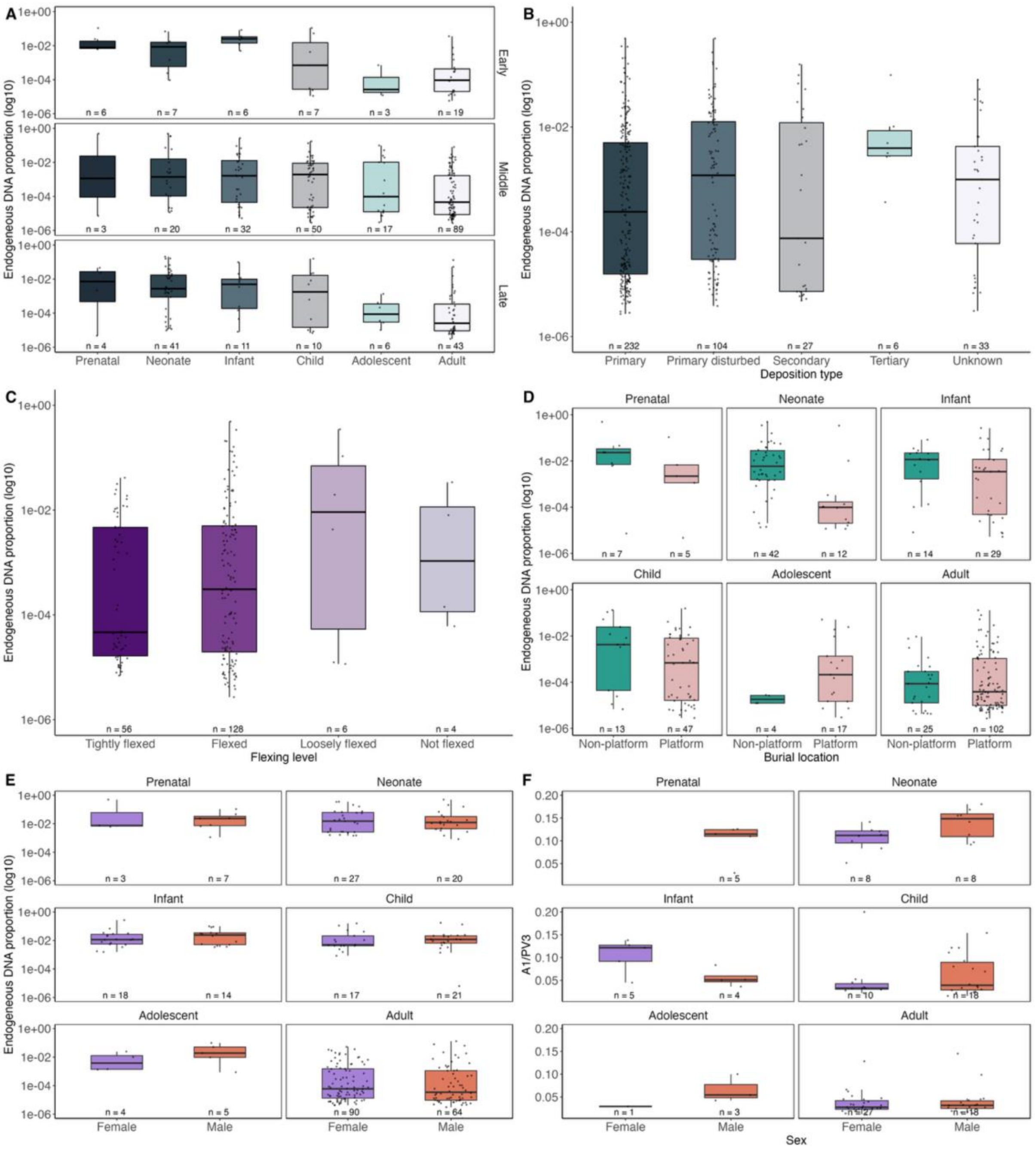


**Figure S18:** Distribution of endogenous DNA yield amount based on **A)** age groups within each period. **B)** deposition types (deposition types refer to how many times a burial opened. “Primary” means undisturbed state. **C)** flexing of Çatalhöyük individuals. **D**) burial location of all Çatalhöyük individuals within the house. In C, the non-platform versus platform distributions were significant (Mann-Whitney test $p<0.001$). The differences were also significant when limiting the sample to subadults ($p<0.001$). **E)** Comparison of male and female endogenous DNA yield for each age category. **F)** Comparison of male and female Amide I / Phosphate (A1/PV3) ratios for each age category.

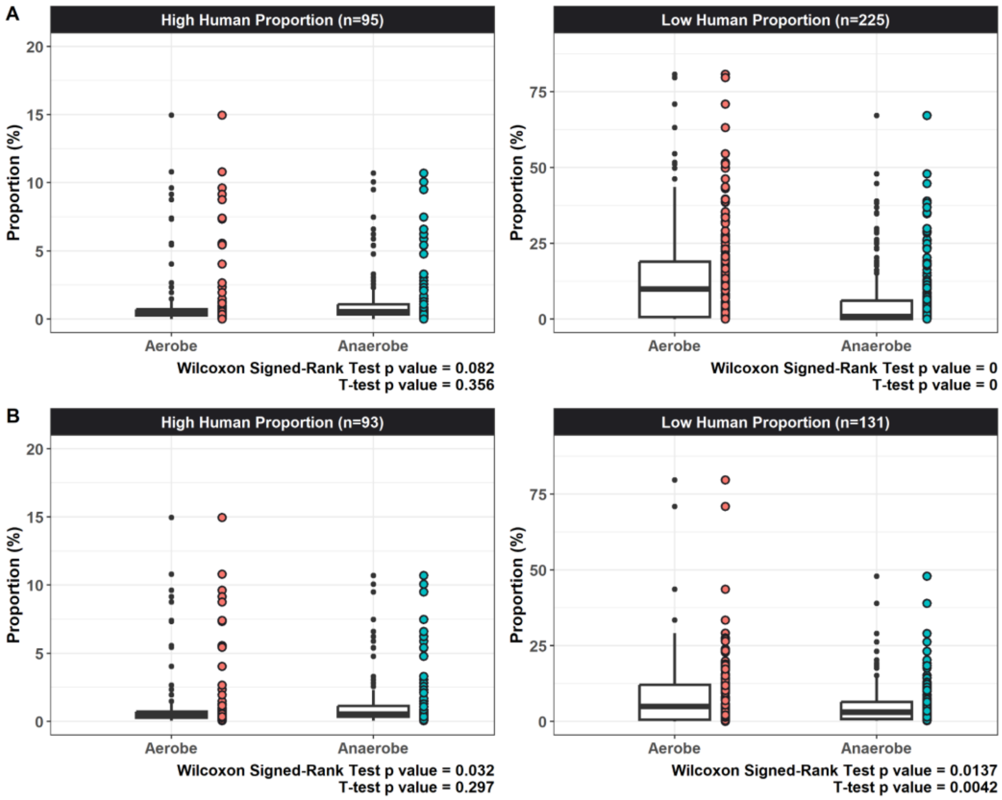


**Figure S19: A)** Unfiltered **B)** filtered proportions of aerobic and anaerobic species for libraries with high and low human endogenous DNA, based on the *KrakenUniq* abundance matrix. The total sample sizes for each level of human DNA proportion are indicated at the top of each plot. The p-values obtained from statistical tests are noted at the bottom right of each plot. See **Table S27**.

### Methods and Supplementary Results

#### 1. Laboratory procedures

##### a. DNA extraction

Experimental procedures were carried out in the dedicated ancient DNA laboratories at Middle East Technical University (METU) and Hacettepe University. To minimise contamination, the laboratories and the equipment were decontaminated with 5-10% NaOCl, DNAway solution and UV irradiation. The labelled bone samples were photographed, recorded, and then brought to the ancient DNA laboratories. During the examination process, the outer surfaces of the petrous bone were cleaned off by a Dremel hand motor. Cleaned bones were cut open to reach the otic capsule region, and a piece of approximately 200 mg was extracted from each sample. Each side of every bone piece was exposed to UV in a crosslinker at 254 nm for 15 minutes and grounded by the SPEX 6875 freezer mill. A fraction of petrous samples were also directly drilled through the otic capsule region, while all tooth samples were drilled through the pulp cavity to obtain c.80-120 mg of powder. All pulverised samples were transferred into 2 ml screw-cap tubes which were labelled by their sample IDs. Two hydroxyapatite negative controls were added during the powdering process for every set.

Extraction buffer (0.45M EDTA, 0.25 mg/ml Proteinase K, pH:8) was added to each powder, and the prepared mix was incubated at 37°C overnight (approximately 18-24h) two times. DNA extraction was performed with lysates by following the Dabney 2013(*2*) protocol. Two extraction blanks were carried through the extraction process. We did not perform UDG treatment on any extracts.

##### b. DNA sequencing library preparation and indexing

Blunt-end, double-stranded and double-indexed Illumina-compatible whole genome libraries were prepared using Meyer and Kircher (2010) (*3*) protocols with Kircher et al. (2012) (*4*) indexes. One negative control was included to the sets at the library preparation and indexing processes. The amplification cycle numbers were estimated using qPCR for each library. After amplification, the enriched libraries were purified using AMPure XP Beads (Beckman Coulter) and then quality and concentration were checked with Bioanalyzer 2100 (Agilent) and Qubit 4 Fluorometer (Thermo Fisher Scientific). Libraries passed quality checks, pooled up to their concentration and sequenced on the Illumina NovaSeq 6000 platform using S1 and S4 flowcells at SciLife Laboratories in Stockholm.

##### c. Fourier Transform Infrared (FTIR) Spectroscopy

We used FTIR analysis for studying organic preservation in Çatalhöyük human skeletal remains, following recent work (*5*). We selected 206 samples covering different ages and sexes (prenatal (n=6), neonate (n=33), infant (n=19), child (n=54), adolescent (n=17) and adult (n=77) **(Table S1).** The samples were powdered in a pestle and mortar and convenient sieves were used to standardize the particle size of the powder to 50-100 µm. IR spectra of the samples were collected using a Universal ATR accessory (Specac Ltd., UK) equipped with a Perkin Elmer Spectrum 100 FTIR spectrometer (Perkin-Elmer Inc, Boston, MA, USA). In the ATR-FTIR spectroscopy technique, the atmospheric $CO_2$ and $H_2O$ absorption bands of the environmental air, and a background spectrum were recorded before the sample spectra were collected. This spectrum was mathematically subtracted from the sample spectra. This process was automatically repeated by the software for each sample spectra. Following the subtraction process, sieved sample powder was placed directly onto the diamond crystal platform with slight pressure to ensure that the samples touched the crystal evenly. The scanning process was performed without drying in the 4000 to 450 $cm^{-1}$ spectral range at room temperature, with 64 scans collected at a resolution of 4 $cm^{-1}$ for each spectrum. The spectrum was collected three times independently for each sample. The average spectra of these three replicates were used for further analysis and data manipulation was performed using *OPUS 5.5* software (Bruker Optics, Reinstetten, Germany).

##### d. Radiocarbon dating

C14 radiocarbon dates of 126 samples are taken from (*6*) **(Tables S1)**. All dates were calibrated using IntCal20.

### 2. Data preprocessing and quality control

##### a. Mapping, trimming, filtering

Residual adapter sequences were removed from raw FASTQ files for each library using the *Adapter Removal* (*7*) (v 2.3.1) software. The removal process included options such as '--*qualitybase 33 --gzip --trimns*' and required a minimum 11 bp overlap between pairs --*collapse --minalignmentlength 11*'. The merged reads were aligned to the human reference genome (hs37d5) using *BWA aln/samse*(*8*) (v 0.7.15) with parameters *'-n 0.01, -o 2'* and seed disabled *'-l 16500'*. Multiple libraries from the same individual were merged using the *SAMtools*(*9*) (v 1.9) tool *'merge'* function, and PCR duplicates with identical start and end coordinates were removed using the *'FilterUniqueSAMCons.py'*(*4*) script. Finally, reads with >10% mismatches to the human reference genome, mapping quality <30, and base pair count <35 were discarded.

Average genome coverage was calculated using the '*genomeCoverageBed'* script from *bedtools2*(*10*), only including reads with a mapping quality >30. To avoid biases, published ancient genome data was remapped and filtered using the same preprocessing steps.

We created trimmed versions of the BAM files by removing 10 bases from end of the reads by using the *'trimBam'* command of *bamUtil* (*11*) (v 1.0.14). Note that all libraries were double-stranded and non-UDG-treated. These trimmed BAM files were then used for genotyping and estimating mtDNA contamination.

#### b. Testing for contamination and quality control

To assess genome authenticity, we employed four methods on each set of libraries, after filtering out reads with minimum base quality and mapping quality <30. First, we analyzed postmortem deamination patterns resulting from cytosine deamination in all samples using *PMDtools* (v 0.60) (*12*) with the *'–deamination'* parameter. Second, we used the *contamMix* (v 1.0-10) (*13*) approach, which considers the rate of consensus mitochondrial sequence mismatches based on a reference panel of 311 mitochondrial genomes. For *contamMix*, consensus mitochondrial sequences were generated using *ANGSD* (v 0.941) (*14*) with the options *'--doFasta 2 -doCounts 1 -minQ 30 -minMapQ 30 -setMinDepth 3 -rMT'*-. Third, we used Schmutzi (v 1.5.7) (*15*), which calculates the probability of authenticity based on the deamination patterns on the consensus mitochondrial sequence, using the information from read lengths and postmortem deamination. Finally, in male individuals, we assessed contamination based on the X chromosome using *ANGSD* with the options *'-i BAMFILE -r X:5000000-154900000 -doCounts 1 -iCounts 1 -minMapQ 30 -minQ 30'*. The probability of X chromosome heterozygosity was subsequently estimated using the R script *'contamination.R'* and reference files from the *ANGSD* (*14*) package.

Five samples with >0.001x coverage were removed from the dataset because of possible contamination (**Table S16**). One of those samples had very low (<2%) PMD scores, and the rest had relatively high (≥0.06) contamination estimates in Schmutzi (classified as "suspicious") and had relatively low (~0.001x) coverages.

For ten samples, *contamMix* values were <0.8 but mtDNA coverages of these samples were also <3x, at which level *contamMix* estimates are known to be unreliable. As no other contamination signal was observed based on other measurements, we did not remove these from downstream analyses (**Table S16**).

We also removed a subset libraries from published Çatalhöyük libraries (*16*), which we identified as showing possible contamination signals upon reanalysis. We note that in previously published Çatalhöyük samples, *contamMix* had been calculated based on transversion (using '--*transverOnly'* command). However, in this study, *contamMix* was calculated including both transition and transversion after trimming 10 bp from two sides. In terms of the results of this calculation, one library from cth747, four libraries from cth728 and three libraries from cth217 were excluded in the analysis to be consistent with the calculations of other samples studied in this work. We used the remaining libraries from these individuals in downstream analyses.

#### c. Merging identical libraries

Two pairs of genomes were found to be genetically identical given *READv2* and *NgsRelatev2* results (see section "**Genetic kinship estimation**" below) (**Figure S24; Table S16**). These genomes could belong to twins or bones of the same person; as we could not reach a definitive conclusion, we decided to merge the pairs into a single genome each. Consequently, the total number of individuals analyzed for relatedness decreased to 131. Merged versions of these individuals were used for all genetic analyses described in this study.

### 3. Genomic imputation

We employed the *GLIMPSE2* (*17*) imputation and phasing tool to impute ancient genomic data and produce diploid phased genomes. First, we calculated genotype probabilities using the BCFtools (*9*) (v 1.18) '*mpileup'* command with parameters *'-I -E -a 'FORMAT/DP' -T -q 30 -Q 30'*, followed by BCFtools (v 1.18) *'call -Aim -C alleles'*(*18*). As our reference panel, we utilized the 1000 Genomes Project (*19*) dataset including biallelic variants, following the *GLIMPSE2* tutorial (odelaneau.github.io/GLIMPSE).

We proceeded by generating imputation regions using '*GLIMPSE2_chunk*' and converting the reference panel to binary format using *'GLIMPSE2_split_reference' script.* Input data included haplotype reference panels, genetic maps, and previously created imputation regions. Finally, we imputed each chunk using the *'GLIMPSE2_phase'* script with default parameters and merged imputed chunks from the same chromosome using the '*GLIMPSE2_ligate'* script.

We performed imputation on all ancient genomes using shotgun genomes with a coverage >0.1x and capture genomes with a target coverage (per targeted SNP) >1x.

Imputation on 0.1x-0.25x genomes has been previously used (*20*) but can potentially lead to low accuracy genotypes (*21*). For this reason, we mainly report results using imputed genomes with either >0.25x or >0.3x original coverage (depending on the tool). At the same time, we performed downsampling experiments to measure any bias that may be introduced by using 0.1x-0.25x genomes on downstream results (e.g. kinship, IBD sharing, or ROH calling). If low coverage imputation did not introduce large biases, we used results from the >0.1x set as additional evidence (see below). In ROH estimation, we did notice biases and therefore only report results from 0.3x genomes.

We applied '*gp>0.99'* genotype probability filtering to imputation results before using them for kinship estimation and ROH analyses by using *BCFtools* (*9*) (v 1.18). No genotype probability filtering was applied for *ancIBD,* following the recommendation in the original publication (*22*).

### 4. Molecular sex assignment

#### a. $K_x$ and $K_y$ approaches

Here, we present two approaches ($K_X$ and $K_Y$) to improve biological sex determination in low-coverage genomes. $K_X$ is a statistic reflecting the copy number of the X chromosome in the individual (relative to autosomes), while $K_Y$ is a statistic indicating whether the sample has a Y chromosome or not. In high-coverage non-contaminated samples, $K_X$ is expected to be 1 for females and to be 0.5 for males. $K_Y$ is expected to be 0 for females and to be 1 for males. The statistics are similar to the $R_X$ (*23*) statistic but are calculated taking into account unmappable regions of chromosomes and mapping ambiguity (see below).

To calculate $K_X$, we first apply a mapping quality filter *'-q 30'* to input BAM files using *SAMtools* (v 1.9) (*9*) and calculate the number of reads on each chromosome using the *SAMtools* (*9*) '*idxstats*' function. This number is divided by the chromosome length to obtain the number of reads per 1 base length for chromosome *i* ($R_{UL,i}$). The chromosome length here does not indicate the total length of the chromosome, as in the $R_X$ method (*23*), but the ungapped length of the chromosome:

$$R_{UL,i} = \frac{Mapped\ reads\ to\ chr\ i}{Ungapped\ length\ of\ chr\ i}$$

Secondly, the standard error for $R_{UL}$ is calculated as follows:

$$SD = \sqrt{\frac{\sum_{i=1}^{22} \left(R_{UL_i} - \underline{R_{UL}}\right)^2}{22-1}} \qquad SE = \frac{SD}{\sqrt{22}}$$

Thirdly, the 95% confidence interval for $R_{UL}$ is calculated as $\underline{R_{UL}} \pm 2.08 \cdot SE$. Because the degrees of freedom for 22 chromosomes is 21, the value to be used for the 95% confidence interval is 2.08.

The $R_{UL}$ ratio calculated from autosomal chromosomes is multiplied by the ungapped length of the X chromosome to calculate the expected number of reads on the X chromosome, $E_{XR}$, for an individual with two copies of the X chromosome. The upper and lower 95% bounds for $E_{XR}$ are calculated as follows:

$$Upper\ E_{XR} = Upper\ R_{UL} \cdot Ungapped\ Length\ of\ chrX \qquad Lower\ E_{XR} = Lower\ R_{UL} \cdot Ungapped\ Length\ of\ chrX$$

The $K_X$ statistic is calculated by dividing the number of reads on the X chromosome observed from the sample by the expected number of reads on the X chromosome. The 95% upper and lower bounds of $K_x$ is calculated as follows:

$$Upper\ K_x = \frac{chrX\ Reads}{Lower\ E_{XR}} \text{ and } Lower\ K_x = \frac{chrX\ Reads}{Upper\ E_{XR}}$$

Our $K_Y$ statistic depends on Y chromosome coverage. To explain our motivation in introducing this, we will briefly summarise chrY nucleotide sequence classes and composition. Although the total length of chrY is ~60 Mb, its ungapped length is only ~26 Mb (in hg19, hg38), since it does not include the heterochromatic region. The Y chromosome consists of the nucleotide sequence classes: Pseudoautosomal region (PAR), X-transposed, X-degenerated, and ampliconic. PAR contains sequences that crossover with the X chromosome. The X-transposed region contains sequences that are 99% identical to the X chromosome, the X-degenerate region consists of distinctive sequences, and the ampliconic region contains of sequences 99.9% identical to other parts of the Y-chromosome (*24*). We note that reads from PAR, X-transposed, and amplionic sequences can map to multiple locations in the genome. These regions have low mapping quality, and the applied quality filter eliminates reads with poor mapping quality (**Figure S20)**.

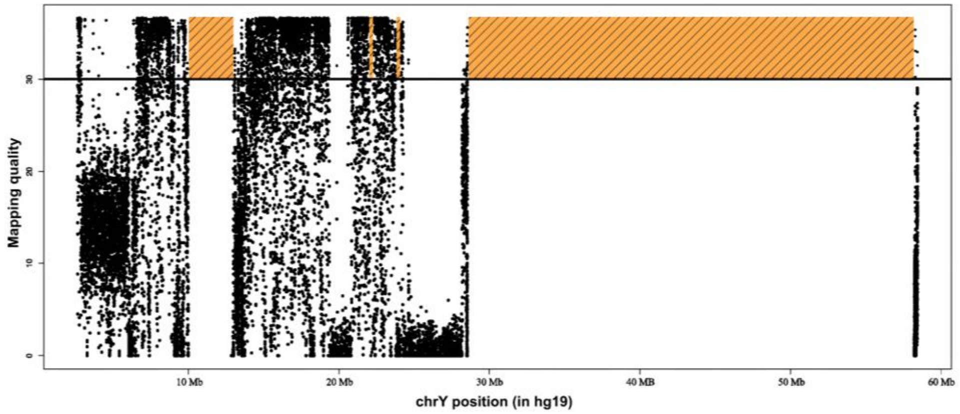


**Figure S20:** Mapping quality averages of a high-coverage ancient human genome per 100 reads by position on the Y chromosome. The x-axis shows the positions on the Y chromosome, and y-axis shows the mapping quality.

To accurately estimate the number of reads expected on Y chromosome (chrY), it is insufficient to rely solely on the ungapped length. This is because nucleotide sequences with high intrachromosomal and interchromosomal similarity often fail to pass the quality filter, resulting in an absence of reads in those regions. Therefore, we first prepared a BED file specifying the positions of mappable (relatively distinctive) sequences on chrY using *BLAT* (*25*). We used the *'intersectBED'* function of *BEDTools* (*10*) (v 2-2.31.0) for subsetting all chrY-mapped reads that fall in mappable regions. Because of the haploid nature of chrY chromosome, we divided $R_{UL}$ by two and then multiplied this by the length of the mappable chrY chromosome (from the BED file) to calculate the expected number of reads on chrY for an individual with one copy of the Y chromosome, $E_{YR}$, and the 95% confidence interval is calculated using the following formulas.

$$E_{YR} = \frac{R_{UL}}{2} \cdot BED\ Length\ of\ chrY$$

$$Upper\ E_{YR} = \frac{Upper\ R_{UL}}{2} \cdot BED\ Length\ of\ chrY$$

$$Lower\ E_{YR} = \frac{Lower\ R_{UL}}{2} \cdot BED\ Length\ of\ chrY$$

$K_Y$ is also calculated as same way with $K_X$.

$$Upper\ K_y = \frac{chrY\ Reads\ intersected\ with\ BED}{Lower\ E_{YR}} \quad and\ Lower\ K_y = \frac{chrY\ Reads\ intersected\ with\ BED}{Upper\ E_{YR}}$$

$K_X$ can be used alone for sex determination, but when used in combination with $K_Y$ it can also detect whether someone has a sex chromosome-related syndrome such as Klinfelter or Turner syndrome. Using both statistics in combination also improves detection accuracy.

Our script for calculating $K_X$ and $K_Y$, as well as the $R_X$ and $R_Y$ statistics is called *'SexDetermine.R'* and is available at https://github.com/mskilic/SexDetermineR).

#### b. Comparison of $K_x$ and $R_x$ approaches

The 95% confidence interval is calculated for $R_X$ as:

$R_{x_i} = \frac{chrX\ Reads}{Total\ Length\ of\ chrX} \cdot \frac{1}{R_{TL_i}}$ , and the confidence interval $= \underline{R_x} \pm 1.96 \cdot SE$

We compared the standard deviations of the $R_{TL}$ statistic ($\frac{Reads}{Total\ Length}$) used as above in $R_X$ and the $R_{UL}$ statistic ($\frac{Reads}{Ungapped\ Length}$) used in $K_X$. Across 131 Çatalhöyük individuals, we found that the coefficient of variation of $R_{UL}$ is 58% less than the coefficient of variation of $R_{TL}$ (**Table S17**), indicating that $K_x$ can be a more powerful approach. The distributions of $R_{TL}$ and $R_{UL}$ values of one Çatalhöyük sample (cch245) exemplify the difference between $R_{TL}$ and $R_{UL}$ (**Figure S21**).

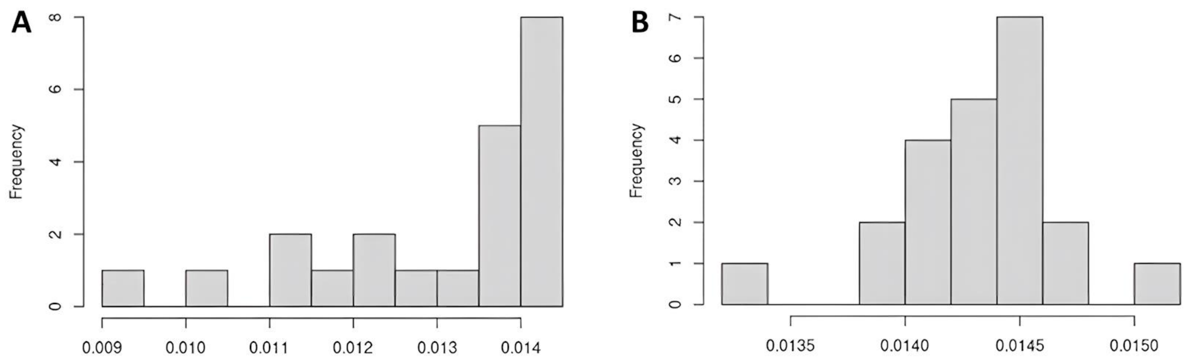


**Figure S21:** The relative depth (*R*) statistics for genomic libraries of individual cch245, comparing the distributions of **A)** $R_{TL}$ and **B)** $R_{UL}$, demonstrating lower variation in the latter.

It can be seen that $R_{UL}$ has a more normal distribution than $R_{TL}$. Considering the dramatic decrease in the standard deviation and the normalization in the distribution, using "ungapped length" instead of the "total length" is expected to render the estimations more accurate in sex determination.

We note that, even if $R_{TL}$ is perfectly normal distributed, since $\frac{1}{R_{TL}}$ is used to calculate the confidence interval, this makes $R_X$ an Inverse Gaussian distribution. Therefore the confidence interval should not be calculated using $\underline{R_x} \pm 1.96 \cdot SE$ (the value of 1.96 here is not strictly appropriate).

#### c. PMD filtering for sex assignment

Because we analysed the molecular sex of libraries with ultra-low endogenous proportions, we used reads filtered by *PMDtools* (*12*) with the *'--threshold 1'* parameter for sex assignment, following Mittnik and colleagues (*23*). We assigned sex only using genomes with >3000 total reads remaining after PMD filtering.

##### d. Turner individual

We identified an individual with Turner syndrome using SexDetermine.R script. This neonate (sample ID: cch148, Unit ID: 10370) represents the earliest known case of mosaic Turner syndrome (45,X0/46,XX) karyotype, dating to the early levels of the Late period. To our knowledge, she is only the second instance documented in ancient DNA literature. The genetic sample includes three sequenced libraries, all of which were assigned to Turner syndrome. Additionally, we ran the SexDetermine.R script on the previously published Iron Age individual identified as having Turner syndrome (*26*), and our script confirms this assignment. The results for our three libraries and the previously published individual are summarized in **(Table S18).**

#### 5. Uniparental haplogroup determination

##### a. Y-chromosome haplogroups

First, we performed genotype calling on the Y-chromosome using *BCFtools* (*9*) (v 1.11) *'mpileup'* function and annotated the Y-SNP names by merging them with the *'ISOGG YBrowse Y-SNP index'*, which contains more than two million Y-SNP entries. (https://ybrowse.org/gbrowse2/gff/snps_hg38.vcf.gz; accessed 2023-11-30), after lifting coordinates from hg38 to hg19 using *CrossMap* (*27*). Then, we identified derived and ancestral genotypes at these SNPs. Subsequently, we utilized *Clade-finder* (https://github.com/hprovyn/clade-finder) to determine Y-chromosome haplogroups by using these derived and ancestral Y-SNP names as input, along with the '*YFull YTree'* (v 11.04.0) (https://github.com/YFullTeam/YTree/blob/master/ytree/tree_11.04.0.zip) as phylogenetic tree. We also repeated the analysis excluding C to T and G to A transitions, which are significantly affected by postmortem damage (as our libraries were not UDG-treated).

##### b. mtDNA haplogroups

First, we performed genotype calling on mitochondrial DNA using *BCFtools (v 1.18) 'mpileup'* function, and using both *'rCRS'* (*28*) and '*RSRS'* (*29*) as reference sequences. We then filtered the results to include only homozygote positions (1/1 or 0/0) in the VCF. Subsequently, we utilized *Haplogrep 3*(*30*) *'classify'* function to determine mtDNA haplogroups using phylogenetic trees, with *'--tree phylotree-rcrs@17.2'* and *'--tree phylotree-rsrs@17.1'* options. We assigned the haplogroup based on the following criteria:

- If using both references, i.e '*rCRS'* (*28*) and *'RSRS'* (*29*), yielded the same result, we assigned that haplogroup to the individual.
- If using both references were consistent but one provided a more specific subclade, we assigned the more specific subclade.
- If using one reference was inconclusive but the other provided a result that matched our existing haplogroup pool (i.e., the list of haplogroups unambiguously called from other Çatalhöyük individuals [**Table S1**]), we assigned that haplogroup.

- If using one reference was inconclusive and the other provided a result that did not match our haplogroup pool, we did *not* assign a haplogroup.
- If using the two references provided different subclades but agreed on a higher-level clade, we assigned the clade they both agreed on.

### 6. Genotyping

#### a. Genome-wide SNP panels

We prepared four SNP panels for different population genetics analyses.

<u>SNP panel 1:</u> The 1000 Genomes (*19*) sub-Saharan African dataset with 4.7M autosomal SNP positions, a relatively unbiased set (*31*) for demographic inference in Eurasia, was used in demographic analyses: outgroup $f_3$-statistics including more than one site, $f_4$-statistics, qpAdm, qpWave, and MDS.

<u>SNP panel 2:</u> We used an extended version of SNP panel 1 to test kinship; here, ascertainment bias is not an issue, and instead we aimed to increase statistical power by using the maximum number of variants. All SNP filtering was performed as described in Koptekin et al. (2023) (*31*), except SNP positions with minor allele frequency (MAF) >1% in any of the five European (Finnish, CEPH, British, Iberian, Tuscany) populations in phase 3 of the 1000 Genomes (*19*) project were added to the SNP panel. After filtering, 6,939,179 autosomal and 282,640 chrX SNP positions have remained. This SNP panel was used for kinship estimations and within-Çatalhöyük analyses with outgroup $f_3$-statistics.

<u>SNP panel 3:</u> Human Origins (HO) Dataset (*32*, *33*) (v.54.1) with 605,775 autosomal SNP positions were called on 1021 present-day Western Eurasian and 249 ancient published genomes, merged with Çatalhöyük samples. This SNP panel was used for PCA analysis.

<u>SNP panel 4:</u> The 1240K Capture Array dataset (*34*) (v.54.1) with 1,121,751 autosomal SNP positions was called on imputed Çatalhöyük genomes to compute runs of homozygosity (ROHs) using *hapROH* (*35*) and haplotype-sharing using *ancIBD* (*22*).

#### b. Pseudo-haploid genotyping

We created pseudo-haploid datasets by randomly selecting one allele for each targeted SNP position. This process involved using the *pileupCaller* (v 1.5.3.1) (https://github.com/stschiff/sequenceTools) on the output from *SAMtools 'mpileup'* (v 1.9) function with minimum base and mapping quality 30 applied to trimmed BAM files. Genotyping was conducted independently for all SNP panels.

#### c. Diploid genotyping

We called diploid genotypes using SNP panel 2 from the imputed dataset using the *'--extract'* option of *PLINK2* (*36*) (v 2.00a4.5) for kinship analysis.

### 7. Population genetic analysis

#### a. Çatalhöyük genomes used in population genetic analyses

We chose a set of Çatalhöyük individuals with >0.03x coverage, and that did not include closely related individuals (if pairs of individuals were identified as second-degree or closer in kinship analyses, we retained the higher coverage individual). We thus made a list of 67 genomes used in various analyses, such as MDS and PCA (**Table S1)**.

We also split this set into three periods, Early (n=10), Middle (n=30), and Late (n=23). This comprised n=63 genomes in total; one individual from the "Final" period and 3 unstratified individuals were not included.

#### b. $f_3$-statistics

$f_3$-statistics were calculated in the form of $f_3$*(Outgroup; Ind1, Ind2)* using *qp3Pop* (v.651) software from *ADMIXTOOLS*(*37*) (v.7.0.2) package. Two different sets were calculated: the first set included pairwise $f_3$-statistics across all possible pairs of Çatalhöyük individuals and calculated with the SNP panel 2 (Methods). The second set was composed of genomes of unrelated individuals (identified as relatedness lower than second-degree in kinship analyses; **Table S2**) from Late Upper Pleistocene and Early Holocene SW Asia and SE Europe, namely Pınarbaşı, Aşıklı, Barcın, Boncuklu, Çatalhöyük, Tepecik-Çiftlik, Musular, Çayönü, Levant HG, Levant N, Iran HG, Iran N and calculated with the SNP panel 1 (Methods) (**Table S19)**.

Within-population genetic diversities were calculated in the form of ($1 - f_3$). The 1000 Genomes (*19*) phase 3 Yoruba population (n=108) was used as an outgroup. Any pair with <2,000 overlapping SNPs was not included.

#### c. Testing change in pairwise $f_3$ diversity over time

The first pairwise $f_3$ set (862 pairs) was used to test genetic diversity changes within Çatalhöyük using the 63 genome set split into three periods (see above). We separated pairwise $f_3$ values into three Çatalhöyük periods. We calculated the significance of the mean diversity differences between any two periods using random permutations of individual IDs. We performed 1,000 permutations and compared the observed mean diversity differences with the expected distribution to calculate p-values. None of the differences were significant (p>0.05) (**Figure S3A**).

#### d. Multidimensional Scaling (MDS)

The second $f_3$ set was used to construct a dissimilarity matrix of pairwise genetic difference ($1 - f_3$) values and summarized using MDS. MDS dimensions were calculated with the *'cmdscale'* function in the *R* (*38*) *'stats'* package, and the first two dimensions were visualized. Only individuals with coverage >0.03x were used in MDS, except for published Iran HG and Levant HG genomes (**Table S19)**. These latter genomes all had coverages <0.03x, but we still wanted to include them in the analysis to better distinguish between regional gene pools. For this reason, we retained individuals as long as they had >2,000 overlapping SNPs with other individuals in the dataset.

#### e. Principal Components Analysis (PCA)

PCA was calculated by projecting ancient individuals onto principal components calculated on genome-wide polymorphism data of 54 Western Eurasian present-day populations (1021 individuals) from the Human Origins SNP Panel (*32*, *39*) **(Table S19)**. Software *smartpca* (v 18140) from package *EIGENSOFT* (*40*) (v 8.0.0) was used to compute principal components, with the *'shrinkmode: YES'* and *'lsqproject: YES'* options. Only individuals with coverage >0.03x were used in PCA (the same set used in MDS except for Iran HG and Levant HG samples, which had low coverages).

#### f. Admixture modelling with *qpAdm*

We modeled the admixture history of Anatolia Pottery Neolithic genomes from C and W Anatolia (Çatalhöyük, Tepecik-Çiftlik, Musular, and Barcın). Past work had shown that these genomes could be explained as admixed between Pre-Pottery Neolithic (PPN) C Anatolia (Boncuklu and Aşıklı) and Levant and Zagros sources (*16*, *31*), while Altınışık, Kazancı and colleagues (*41*) had shown that U Mesopotamia could be a possible source of eastern/southern ancestry in C Anatolia. We thus tried a simple 2-way admixture model with C Anatolia PPN (Boncuklu and Aşıklı) and Upper Mesopotamia PPN (Çayönü) as sources to explain admixture Anatolia Pottery Neolithic genomes. Here we used the 63 Çatalhöyük genomes split into three periods. We performed modelling using the *qpAdm* (v 1520) software from the *ADMIXTOOLS*(*37*) (v 7.0.2) package, using the *'allsnps: YES'* option, suitable for low-coverage data. Two different reference sets, differentially related to left populations, were selected **(Table S19)**.

- First set: Mbuti, Ust_Ishim, Kostenki14.SG, MA1, Han, Papuan, Dai, Chukchi, Mixe, CHG, WHG, AfontovaGora3, and Anatolia HG.
- Mbuti, Ust_Ishim, Kostenki14.SG, MA1, Han, Papuan, Dai, Chukchi, Mixe, Iran Ganj Dareh N, WHG, AfontovaGora3, and Anatolia HG. Only shotgun sequences except AfontovaGora3, and Anatolia HG.

The sets were modified from (*31*). We preferred shotgun-sequenced genomes where possible to avoid technical bias (*31*).

If a model had p-value > 0.01 (based on qpAdm Z-scores), and admixture coefficient sum of source 1 and source 2 is = 1, it was classified as feasible (cannot be rejected). Otherwise the model was rejected.

#### g. Testing change in pairwise haplotype diversity over time

We estimated haplotype-based diversity within Çatalhöyük as an alternative measure to $f_3$ diversity for testing possible temporal changes in the gene pool. Haplotype-based diversity is expected to be more sensitive to gene flow from genetically closely related populations as we show in population genetic simulations (see section 'Population genetic simulation results and discussion').

Haplotype-based diversities within each period were measured using n=46 imputed genomes from Early, Middle and Late periods (with >0.1x coverage) (excluding 3 that were from the Final period or unstratified). Imputed variant sites missing in at least one genome were filtered out. The phased genotype data were analyzed in windows of 300 SNPs, which correspond to ~10 Kbs. For each SNP window, we generated a distance matrix of haplotype similarities. If two chromosomes compared had the same alleles across all SNPs in a window, they were assigned a match, while if they had at least one difference they were considered a mismatch (irrespective of the distance between haplotypes).

We then calculated the total number of windows divided by the total number of pairwise compared windows within each period, using a custom *R* script. Note that each individual was represented by both of its autosomal chromosomes.

The change in haplotype diversities between periods was tested by a random permutation of individuals (see section 'Testing change in pairwise genetic diversity over time' above).

#### h. $f_4$-statistics

All $f_4$-statistics were calculated using *qpDstat* (v 980) software from the *ADMIXTOOLS*(*37*) (v 7.0.2) package. The 1000 Genomes(*19*) phase 3 Yoruba population (n=108) is used as an outgroup.

Four different settings were used for $f_4$-statistics.

- $f_4$-Statistics1: This was used to investigate possible genetic outliers of non-Anatolian descent. We performed $f_4$-statistics in the form of $f_4$*(Outgroup, SurrPop; AnatInd, AnatPop)*, where *SurrPop,* contemporary or earlier populations from surrounding regions (Levant N, Iran N, CHG, and Balkan HG) were tested against any Anatolian Neolithic individual *AnatInd* (from Barcın, Boncuklu, Aşıklı, Çatalhöyük, Çayönü, Musular) versus the population from its own village, *AnatPop* (e.g. Barcın1 tested against the pool of all other Barcın individuals). This aimed to determine genetic outliers, where a surrounding population was significantly choosing specific individuals over the pool of other individuals from the same village.
- $f_4$-Statistics2: This was used to investigate possible genetic outliers of Anatolian descent in each village. We performed $f_4$-statistics in the form of $f_4$*(Outgroup, AnatInd; AnatPop, AnatSurrPop),* where each Anatolian Neolithic genome, *AnatInd,* was tested against its own population (*AnatPop*) or other populations from Anatolia or surrounding regions (*AnatSurrPop)*. We thus determined if an Anatolian individual was significantly choosing other populations instead of its own.
- $f_4$-Statistics3: This was used to investigate possible genetic differentiation between three Çatalhöyük periods. We performed $f_4$-statistics in the form of $f_4$*(Outgroup, $Period_1Ind$; $Period_1Pop$, $Period_2Pop$)*, where each Çatalhöyük genome, *$Period_1Ind$,* was tested against its own period (*$Period_1Pop$*) or other two Çatalhöyük populations (*$Period_2Pop$)*. We thus

determined if a Çatalhöyük individual was significantly choosing other periods instead of its own.
- $f_4$-Statistics4: This was used to investigate if any of the Çatalhöyük periods have higher affinity to other Anatolian Neolithic populations or if other Anatolian Neolithic or surrounding populations have higher affinity to any of the Çatalhöyük periods. We performed $f_4$-tests in the form of $f_4(Outgroup, Period_1; Period_2, AnatSurrPop)$ and $f_4(Outgroup, AnatSurrPop; Period_1, Period_2)$.

In the first three $f_4$-statistics and *qpWave* analyses (see below), the composition of *AnatPop* (genomes from an Anatolian site) or *Period₁Pop (*genomes from a Çatalhöyük period) changed in each test, where the tested individual was removed while all other individuals remained. However, we did not include two genetic outlier genomes in their village populations: cay008 from Çayönü (*41*) and Ash033 from Aşıklı (*16*). $f_4$-Statistics4 was computed on entire populations. In order to avoid technical bias we used shotgun-sequenced genomes for all $f_4$-statistics.

All statistists with |Z| > 2 were considered nominally significant.

#### i. qpWave analysis

Possible genetic differentiation within Anatolian Neolithic settlements, namely Aşıklı, Boncuklu, Barcın, Çatalhöyük, Çayönü, Tepecik-Çiftlik and Musular gene pools, was tested similar to analyses using $f_4$-statistics, comparing a genome from a site from all other genomes from the same site (the leave-one-out approach described above). We ran the *qpWave* (v 1520) software from the *ADMIXTOOLS*(*37*) (v 7.0.2) package with the *'allsnps: YES'* option (suitable for low-coverage data). We used the same outgroup sets used in qpAdm (see above). The null hypothesis (i.e. a population can be explained by one gene pool) was rejected for the tests that had p-value < 0.05 (**Table S5**) We preferred shotgun-sequenced genomes whenever possible to avoid technical bias (*31*).

#### j. Identifying putative outliers

Putative outliers were defined as individuals who were either;

- an outlier in the PCA plot (visually identified),
- an outlier in the MDS plot (visually identified),
- showed nominally significant (|Z| > 2) affinity to the other Anatolian and surrounding populations based on $f_4$-Statistics1, indicating higher affinity of an external genetic profile to that individual compared to its compatriots,
- showed nominally significant (|Z| > 2) affinity to the other Anatolian and surrounding populations based on $f_4$-Statistics2, indicating higher affinity of the individual to an external genetic profile than to its compatriots,
- had a p-value < 0.05 in qpWave analysis when tested against its own population.

The proportion of putative outliers were calculated as the total number of individuals who were identified as putative outliers in any of the five tests described above, divided by the total number of tested individuals. Since only shotgun-sequenced genomes were tested in $f_4$-statistics and qpWave, only shotgun individuals >0.03x were included in the outlier tests. Therefore, the capture-generated Barcın individual I1103, who was a PCA outlier, was not included in this calculation.

### 8. Coalescent simulations for interpreting diversity estimates

We performed population genetic simulations for three purposes: a) to test if haplotype-based diversity measures would be more sensitive to admixture than allele frequency-based measures (i.e., $f_3$), as may be theoretically expected, b) to evaluate the observed lack of significant temporal increase in diversity measures (**Figure S3**), and c) to assess the observed loss of homogeneity over time with $f_4$ statistics.

#### a. Population genetic simulation framework

We used *msprime* (*42*) (v 1.0) to simulate 100 Mbs of diploid sequences for two demographic scenarios; no gene flow versus continuous symmetric gene flow between sister populations A and B, which would be representing Çatalhöyük and a neighbouring gene pool (**Figure S22**). The divergence time between A and B was chosen as 50 generations ago, followed by the divergence of ancestral population AB and the outgroup O 150 generations ago. The recombination and mutation rates were both chosen as 1e-8, per generation. The migration rate between A and B for the second scenario was set to 0.01 per generation. The population sizes of ABO, O and AB were fixed at 3000, while A and B were given a bottleneck from 3000 to 1000 individuals, 5 generations subsequent to their split. Both scenarios had 100 replicates and 25 individuals were sampled from populations A and O for the calculation of $f_3$-based and haplotype-based diversities. Outgroup $f_3$ values between individuals within population A were calculated using *tskit* (*43*) (v 0.5.6), recording the median $1\text{-}f_3$ value per replicate. We followed the same approach used for real data (see section "Testing change in pairwise haplotype diversity over time") when calculating haplotype diversities from the simulated genotype data, written to vcf files with *tskit* (*43*) (v 0.5.6), and recorded the median diversity value of windows per file. In order to compare the effect size of gene flow when using $f_3$-based versus haplotype-based diversity estimates, we computed the Cohen's D effect size statistic for both cases.

To study temporal change in diversities measured using $f_3$ statistics, we used the same set-up except the outgroup split time of 3000 generations for a second set of simulations where we sampled from population A at three different time points: generations -30 (A1), -15 (A2) and 0 (A3). We had three scenarios where the continuous migration rate between A and B were 0, 0.01 and 0.1. Again sampling 25 individuals from populations A and O, we recorded the median $1\text{-}f_3$ values for A1, A2 and A3 to calculate the change in diversities through time. We also tested the effect of migration from a more distant population X, split from the ancestral population AB 300 generations ago, to determine the parameter values where the diversity increases from A1 to A3. For this, we checked combinations of higher $N_e$ values for both X and A (5,000 and 10,000) with a migration rate of 0.01.

Finally, to evaluate the change in homogeneity over time as measured using $f_4$ statistics, we used the same set-up used to test the change in temporal diversities with $f_3$, where we sampled from population A at three different time points. In addition to a scenario with no migration, we had scenarios where the migration rate between population A and B, and A and X, were set to be 0.01. For all period combinations, $f_4$ statistics of the form *$f_4$(O, Ay-individual, Ay-pop, Az-pop)*, where An-individual corresponds to an A individual from period y (i.e. A1), Ay-pop to population A at period y, and Az-pop to population A at period z different from y (i.e. A2 or A3), were calculated. We tested Ne values of 1,000 and 5,000 for A, the Ne of B and X were set to be the same as A.

#### b. Population genetic simulation results and discussion

We first compared the sensitivity of the two diversity measures, *1-$f_3$* and haplotype similarity, with simulated genomes. Using a simple demographic model in **Figure S22**, we compared the median pairwise diversity values obtained from sampling n=25 individuals per population, for n=100 replicates, under scenarios of no gene flow versus continuous gene flow from a closely related population. We found that the effect size (i.e., the impact of simulated migration) calculated when using haplotype similarity was about 2x higher than calculated using *1-$f_3$* (**Figure S23**), confirming our expectation.

We then asked to what extent the *1 - outgroup $f_3$* measure could capture the effect of short-term gene flow from a closely related population (one that split 50 generations ago in our simulations). For this, we sampled individuals from the target population in three different time points, and compared the median 1-$f_3$ values for n=100 replicates. Testing different effective population size values and rates of gene flow from a population recently split from the target, no increase in $f_3$-based diversity estimates was observed between subsequent time points (**Figure S24**). Hereupon, we tested gene flow from a more distant population, X (one that split 300 generations ago in our simulations). A change in diversities was observed only under scenarios where the target population had a high $N_e$ (e.g., 10,000), in other words, low drift (**Figure S25**). We, therefore, conclude that capturing gene flow through temporal diversity samples using *1-$f_3$* values can only be possible when the source of gene flow involves a distant population and the population size is large; otherwise the effect of drift swamps the gene flow effect.

Lastly, we investigated the conditions for loss of homogeneity over time as observed in the real data with gene flow using $f_4$-statistics. In the real data (**Figure 2**), the individuals from Early period had significantly higher affinity to their contemporaries than to other periods, but this was weaker or not observed in the Middle and Late periods. We could capture this pattern using scenarios with gene flow from a relatively distant population, X. When Ne of both X and A was set to be 5,000, the majority $f_4$-statistics of the form *$f_4$(O, A1-ind, A1-pop, A2/3-pop)* were negative, indicating a higher affinity of the individuals from the first period towards each other. However, an affinity of a similar extent was not observed for individuals from periods A2 and A3 **(Figure S26).**

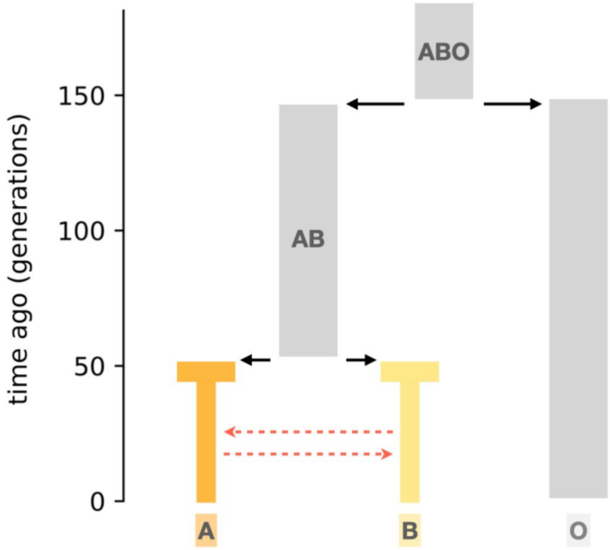


**Figure S22:** Migration model. The model was used to study the effect of gene flow in population A with (Ne=1000). We compared the sensitivities of allelic diversity (pairwise *1 - outgroup* $f_3$ within a population) and haplotype diversity to capture gene flow.

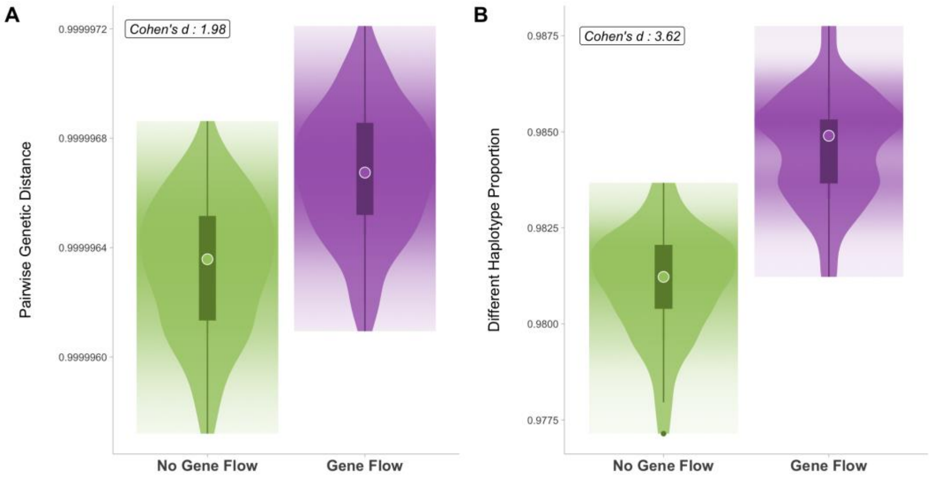


**Figure S23:** Sensitivities of allelic diversity and haplotype diversity to gene flow in a population genetic simulation. **A)** Allelic diversity calculated as pairwise *1 - outgroup* $f_3$ within a population. **B)** Genomic haplotype diversity is calculated as the proportion of pairs with different haplotypes. Cohen's D statistic was used to measure the shift in diversity caused by gene flow between the two scenarios, shown in the inset.

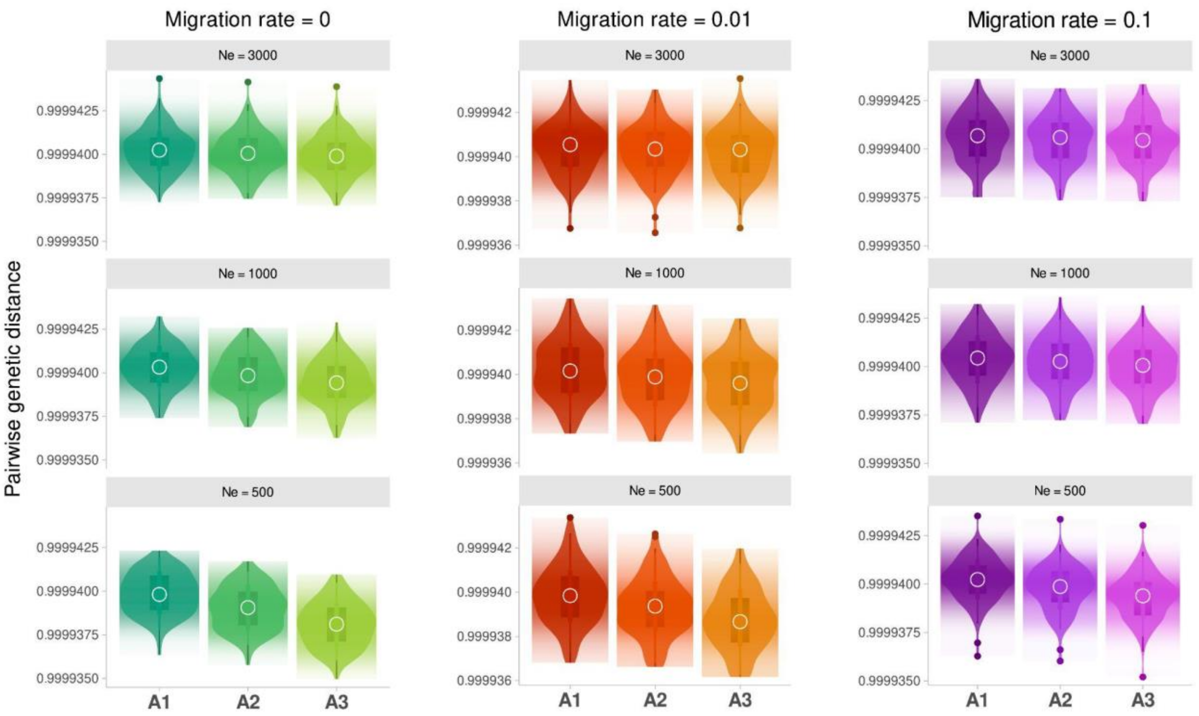


**Figure S24.** Change in temporal diversity within the simulated population A with continuous gene flow from a closely related population. The median *1 - outgroup* $f_3$ values were used to measure diversity per n=100 replicates. n=25 individuals were sampled from population A at different time points: generation -30 (A1), -15 (A2) and 0 (A3).

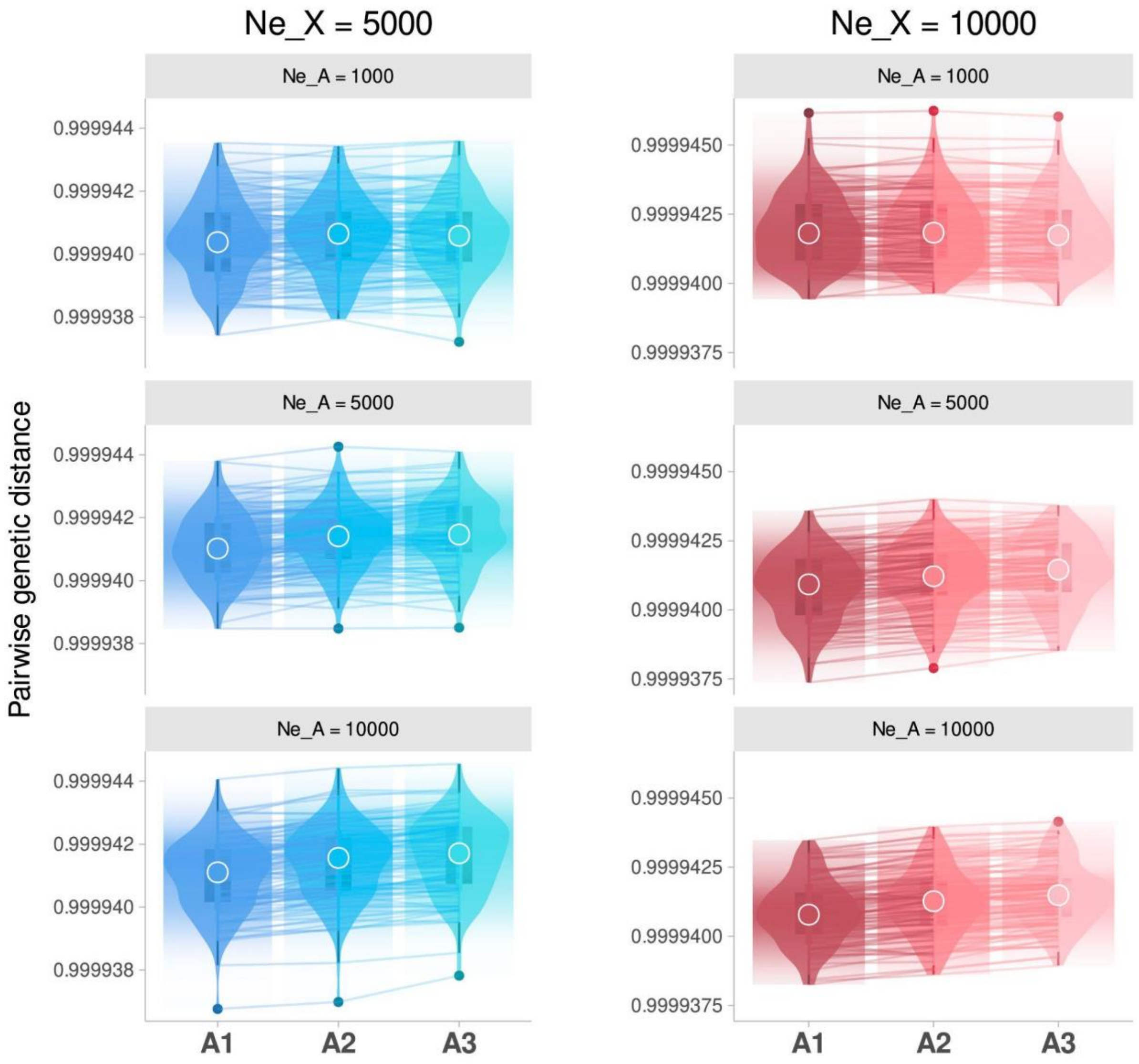


**Figure S25.** Change in temporal diversity within the simulated population A with continuous gene flow from a distant population X. The migration rate was set as 0.01. The median *1-$f_3$* values were used to measure diversity per n=100 replicates. n=25 individuals were sampled from population A at different time points: generation -30 (A1), -15 (A2) and 0 (A3). The lines connect the change in median *1-$f_3$* values per replicate.

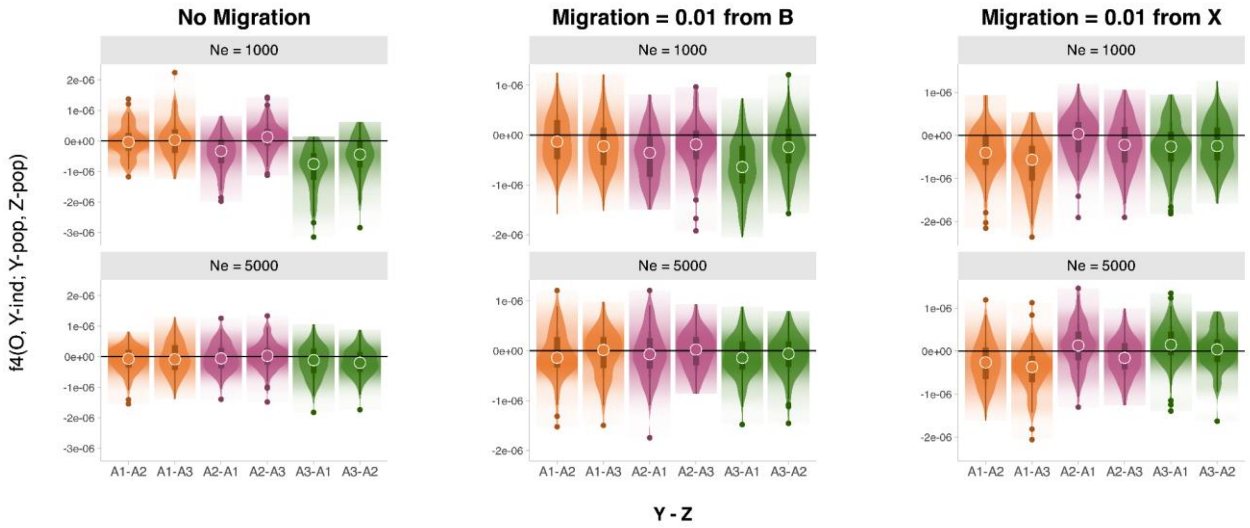


**Figure S26.** Change in homogeneity over time in population A with migration from a close population B and a distant population X, at different Ne values for A. Per n=100 replicates, n=25 individuals were sampled from population A at different time points: generation -30 (A1), -15 (A2) and 0 (A3). The $f_4$-statistics of the form $f_4$(O, Y-ind, Y-pop, Z-pop) where Y-ind corresponds to any individual from period A1, A2 or A3, Y-pop to the population from which the individual was sampled from, and Z-pop to the population A1, A2 or A3 different than Y. A negative $f_4$ value shows that the individual has higher affinity to its contemporaries than to other periods.

### 9. Detecting IBD segments

We investigated pairs sharing identical-by-descent (IBD) haplotype segments using the software *ancIBD* (*22*) (v 0.6). This algorithm uses a Hidden Markov Model to estimate long shared blocks (>8 cM) between two imputed genomes that are likely IBD, i.e., inherited from a relatively recent common ancestor. Our study utilized a subset of the imputed genome dataset containing 1240K SNPs. We included n=49 Çatalhöyük genomes >0.1x coverage for analysing third-degree or higher relatedness and n=18 Çatalhöyük genomes >0.25x coverage, along with n=21 published shotgun-sequenced genomes of >0.25x coverage and 1240K SNP capture data of >1x target SNP coverage for analysing distant relatedness between different Neolithic sites, following (*22*). The *'hapBLOCK_chroms'* function in *ancIBD* was employed with default parameters to estimate shared blocks >8 cM for each chromosome.

IBD connections among sites were estimated using IBD segments (>12 cM), as 8 cM sharing appeared too ubiquitous and difficult to interpret. First, we summed the total shared IBD segments (>12 cM) among sites. Since sample sizes were heterogeneous among sites, we normalized the total number of shared IBD segments by dividing them by the maximum number of shared IBD segments among pairs of individuals of the same length range (i.e. by normalizing sharing). We estimated IBD-sharing frequency between a pair of sites by dividing the proportion of IBD-sharing calculated among sites by the total number of possible pairs (e.g. between two different sites of n1 and n2 individuals there would be n1*n2 comparisons).

### 10. Genetic kinship assignment

We estimated genetic kinship coefficients (θ) between each pair of Çatalhöyük individuals with sufficient coverage using multiple tools simultaneously, an approach we have argued based on simulations is the most reliable (*44*). We thus estimated θ using both *READv2* (*45*) and *NGSRelate v2* (*46*). P0 values of *READv2* (*45*) were normalized to calculate $P0_{norm}$ using the median P0 values of pairs with >10,000 common SNPs for autosomes, and all pairs with >500 common SNPs for the X chromosome. The kinship coefficient (θ) was calculated as 1 - $P0_{norm}$. For autosomal relatedness, *READv2* (*45*) was computed with 49 imputed samples (with coverage >0.1x) and 82 non-imputed samples, using the new functionality of *READv2* that allows processing mixed diploid/pseudo-haploid data. For relatedness calculations based on the X chromosome, *READv2* was performed on non-imputed samples.

*NGSRelate v2* (*46*) was only computed with non-imputed samples. To increase the accuracy of background allele frequencies that *NGSRelate v2* uses for kinship estimation, we calculated background allele frequencies from a pool of 115 Late Upper Pleistocene/Early Holocene C, W, and SE Anatolian genomes (Pınarbaşı, Aşıklı, Boncuklu, Musular, Çatalhöyük, Tepecik-Çiftlik, Barcın, Çayönü) with coverages >0.05x (**Table S2**). A MAF filter >0.05 per SNP was applied to the dataset when running both methods. We noticed that using a lower MAF cutoff increases the number of SNPs used but introduces noise, as we observed in a number of trials.

#### a. Relatedness classification cutoffs

Five different relationship categories (identical, first-degree, second-degree, third-degree, unrelated) were assigned to pairs based on the arithmetic mean of the theoretical θ and used as cutoff values for relationship assignment (*44*). Individuals with θ>0.37500 are classified as identical, between 0.37500-0.18750 as first-degree, between 0.18750-0.09375 as second-degree, 0.09375-0.03125 as third-degree and <0.03125 as unrelated. Note that distant relatives beyond the third-degree would be assigned as unrelated in this classification scheme. However, even third-degree estimation is highly noisy when only low-coverage genomes are available and the main source of information is allelic mismatch (*44*). We therefore did not attempt to estimate beyond third-degree.

#### b. Autosomal relatedness estimation

##### i. The first round of kinship estimates

Relatedness was assigned to each pair of the 133 samples by *READv2* (*45*) and *NGSRelate v2* (*46*) with common SNPs >1000. Overall, there was sufficient concordance between the two tools, even though they use partly different sources of information (Spearman correlation coefficient r=0.38, p<0.001) (**Figure S27).** If both *READv2* and *NGSRelate v2* made the same estimation, this was assumed correct. If the estimates from the two tools were inconsistent, we assigned the pair to a fuzzy category (e.g., if the tools estimated first- and second-degree, we used ‘First- or second-degree’). Out of 6874 pairs, 890 (13%) were assigned to this category.

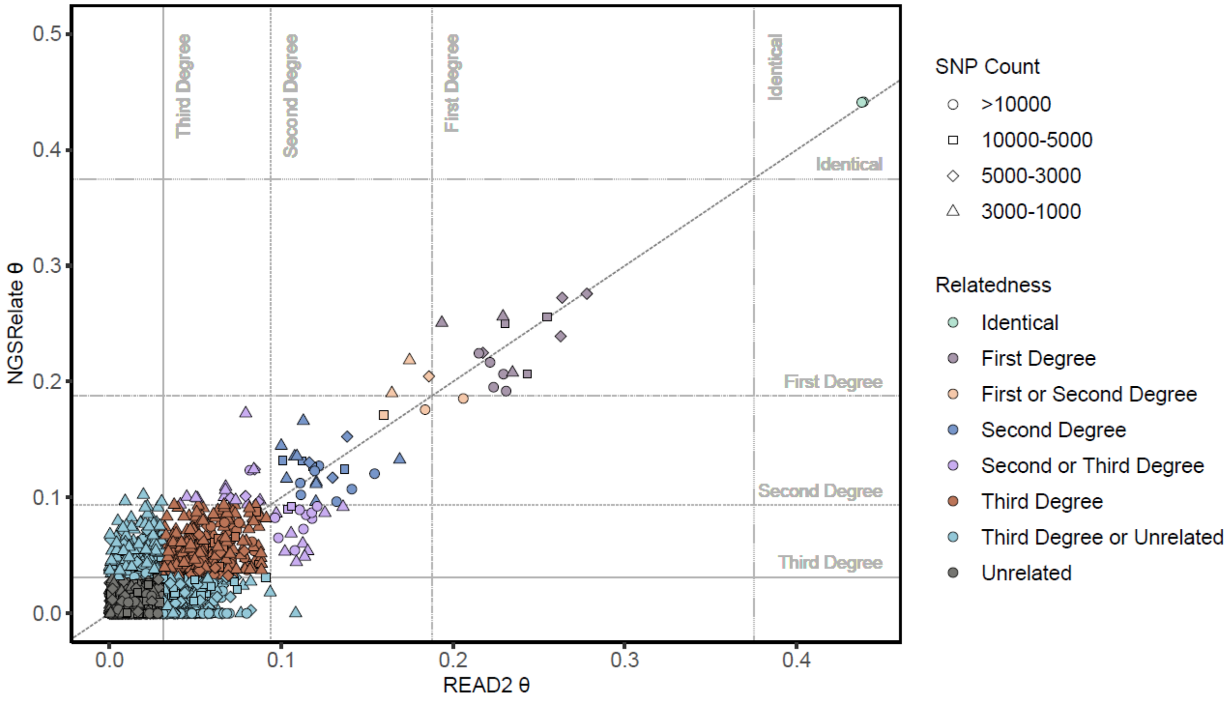


**Figure S27:** Comparison of *READv2* and *NGSRelate v2* θ estimates. Horizontal and vertical lines show the arithmetic mean of theoretical θ cutoffs used for relatedness assignments and the gray solid diagonal line is x=y line. Only pairs with θ>0 were shown for clarity (negative θ estimates can be encountered among unrelated pairs due to background noise). Note that the two identical pairs were merged in later analyses.

#### ii. Downsampling experiments to test the effect of imputation on kinship estimates

A series of downsampling experiments described below were conducted to observe the effect of imputation (see section ‘Genomic imputation’) on relatedness estimations, whereby we estimated kinship between original genotypes and downsampled-and-imputed genotypes. We used *READv2* and normalized P0 values with median P0 values from the original non-imputed dataset.

We prepared four different datasets to observe the effect of imputation on imputed/non-imputed pairs, especially when the non-imputed pair had very low coverage. Dataset 1 comprised the 49 Çatalhöyük genomes (non-imputed) with coverage >0.1x. In Dataset 2, the same 49 samples were imputed and pseudo-haploidised by a custom *Python* script named *‘haploidize.py’* (https://github.com/MerveNurGuler/Haploidize-VCF). In Dataset 3, non-imputed versions of the same genomes were downsampled to 0.01x. In Dataset 4, these were downsampled to 0.001x.

Dataset 1 and Dataset 2 were combined and filtered to retain only pairs where one individual was imputed and the other non-imputed (e.g., original cch251 and imputed cch102, or original cch102 and imputed cch251). This combined dataset is used as the reference set.

Dataset 2 and Dataset 3, as well as Dataset 2 and Dataset 4, were combined similarly and named as the 'imputed vs. 0.01x' set and the 'imputed vs. 0.001x' set, respectively.

Results of the reference set were compared with the 'imputed vs. 0.01x' set (**Figure S28A**) and the 'imputed vs. 0.001x' set (**Figure S28B**) by computing the Spearman r between θ values and testing significance using Mantel tests. The macro average F1 score was calculated from the confusion matrix using the *'ml_test'* function of the *'mltest'*(*47*) library in *R* for the kinship category assignments.

In both cases, θ values estimated in different datasets were well correlated, with r=0.67, p<0.001 for the 'imputed vs. 0.01x' set, and r=0.321, p<0.001 for the 'imputed vs. 0.001x' set. Meanwhile, the F1 score of the 'imputed vs. 0.01x' set was also high (F1=0.924), but the F1 score of the 'imputed vs. 0.001x' set was modest (F1=0.514).

Studying **Figure S28B** further**,** we concluded that even though relatedness estimates from the 'imputed vs. 0.001x' set (i.e., when one individual is imputed but the other ultra-low coverage) were not overall reliable, we could still identify first- or second-degree related pairs with sufficient confidence. Hence, we used evidence from imputed vs. ultra-low-coverage (down to 0.001x) pairs only when they indicated first or second-degree relatedness.

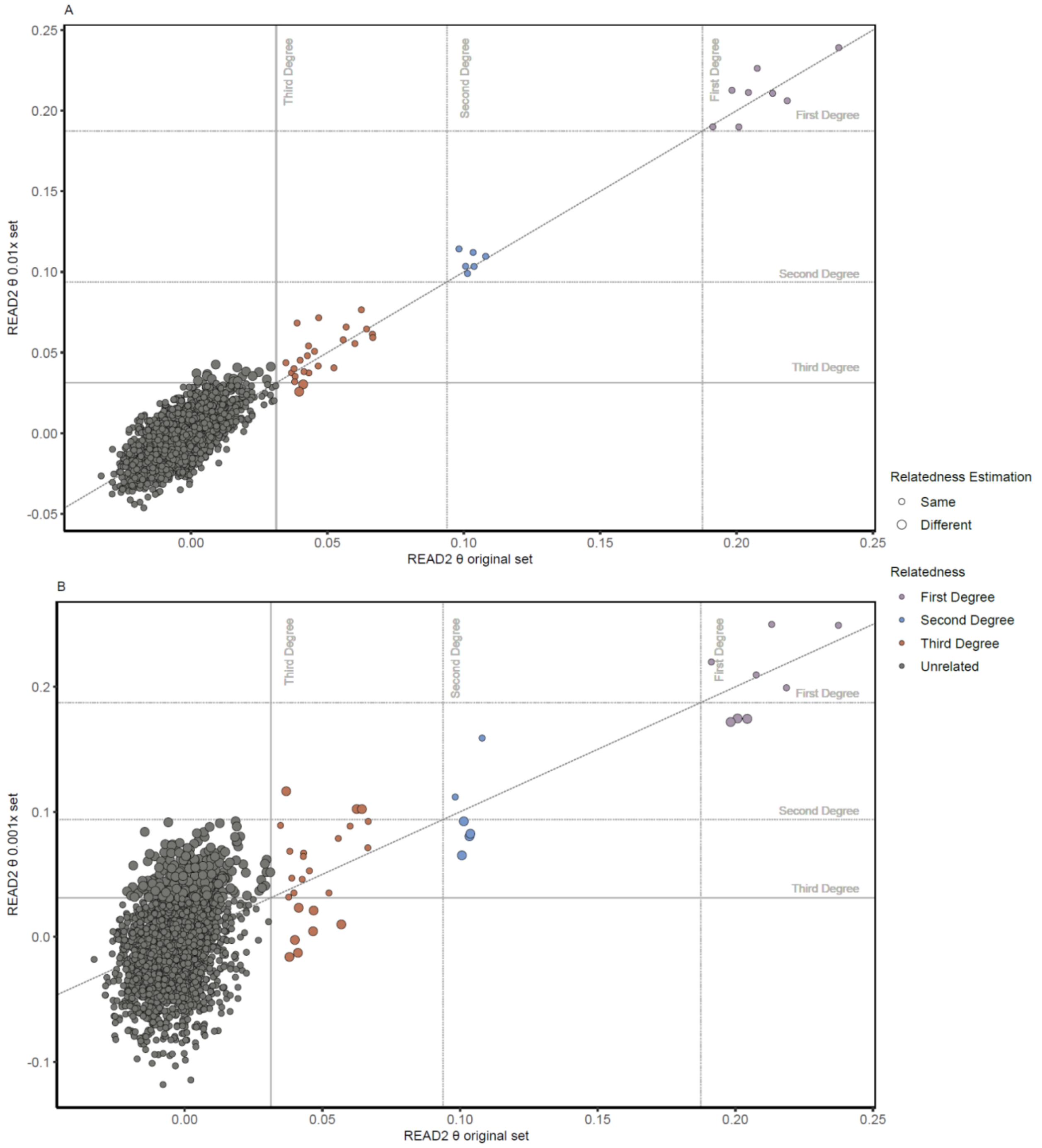


**Figure S28:** Comparison of *READv2* θ values between **A)** the reference set and the 0.01x set and **B)** the reference set and the 0.001x set. Horizontal and vertical lines show the arithmetic mean of the theoretical θ cutoff values used for relatedness assignment and the gray solid diagonal line is the x=y line. Relatedness types were colored based on the results of the original set.

Lastly, we studied the effect of using diploid or pseudo-haploid calls in kinship estimates using *READv2* (*45*), which accepts both diploid and pseudo-haploid data (including pairs where one is diploid and the other pseudohaploid). For this we prepared a dataset that included all 49 genomes with coverage >0.1x, represented as imputed genotypes, and the remaining 82 genomes with coverage <0.1x as pseudo-haploid genotypes. Two datasets were prepared; in the first one, imputed data was pseudo-haplodised and relatedness was estimated by the standard *READv2*, and in the second one, imputed data was kept in a diploid state and relatedness was estimated by the modified *READv2* script. Both θ values (r=0.866, p<0.001) and relatedness assignments (macro average F1=0.902) were highly consistent (**Figure S29**. Consequently, the dataset with diploid imputed data is used for final relatedness estimation with *READv2*.

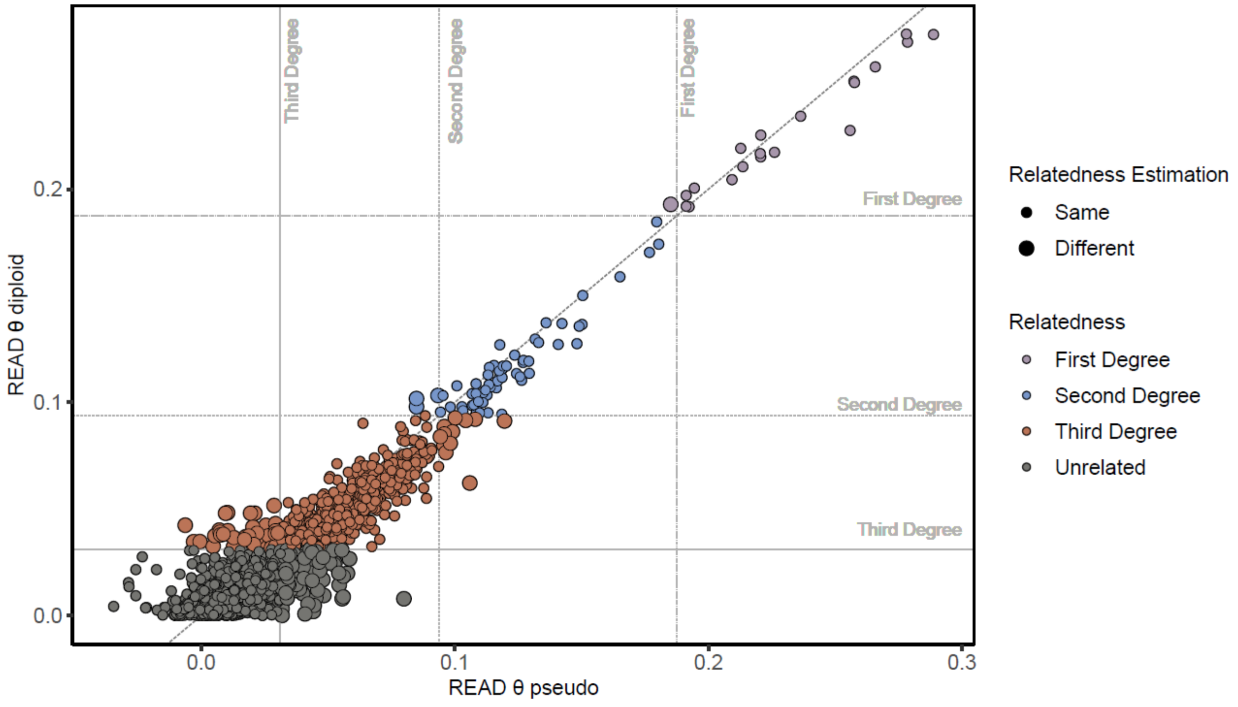


**Figure S29:** Comparison of *READv2* θ estimates involving diploid and pseudo-haploid calls of the same pairs. Horizontal and vertical lines show the arithmetic mean of theoretical θ cutoff values used for relatedness assignment and the gray solid diagonal line is x=y line. Relatedness types were colored based on the results of the diploid dataset. Only pairs with θ>0 were shown for clarity (negative θ estimates can be encountered among unrelated pairs due to background noise).

#### iii. Close relatedness estimation using *ancIBD* results

We estimated shared haplotype segments between the 49 imputed samples with coverage >0.1x using *ancIBD* (*22*). Third-degree or higher relationships were assigned based on plotting the total sum and the total number of shared segments >12 cM and detecting clusterings of the related individuals, as described in (*22*) **(Table S8)**. A possible grandparent-grandchild relationship was also detected in the second-degree clustering (**Figure S30**).

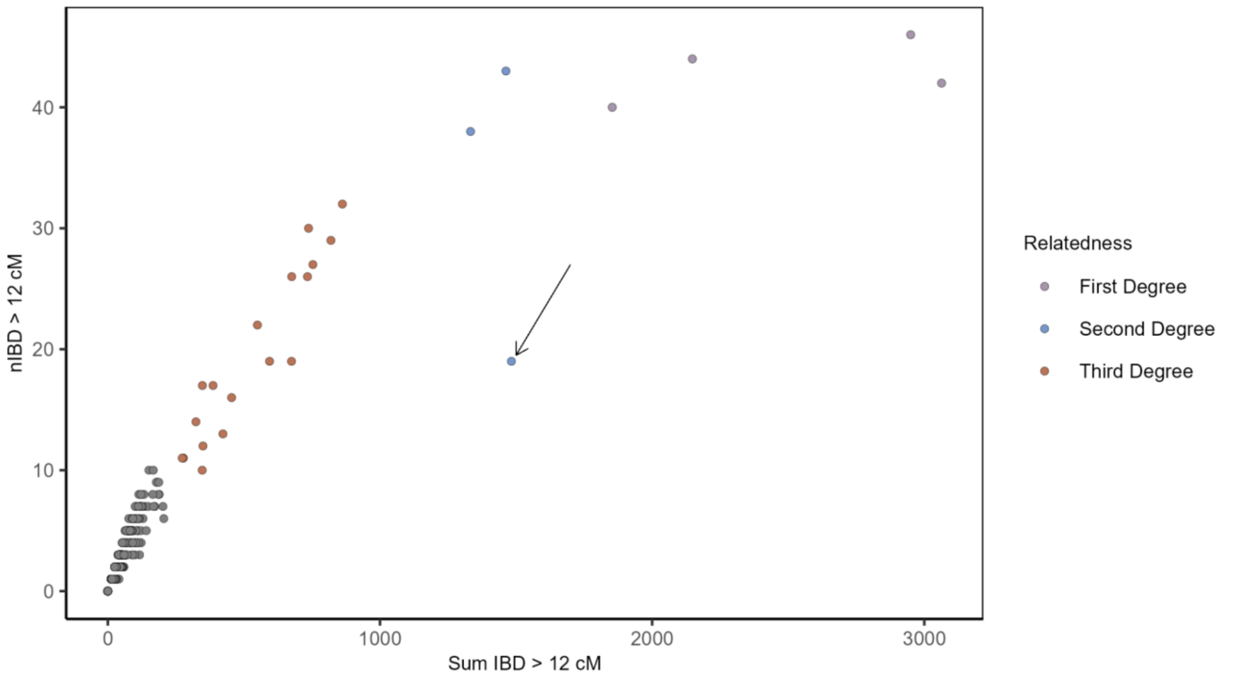


**Figure S30:** The sum vs. number (nIBD) of shared IBD segments larger than 12 cM. The arrow shows a possible grandparent-grandchild relation, evaluated based on simulation results presented in (*22*).

#### iv. The final round of relatedness assignments

The final relatedness assignment of each pair was made based on the following set of rules, chosen based on the above downsampling experiments and our previous work (*44*). We aimed to use the maximum amount of information while minimizing false negatives or false positives, both of which could impact downstream analyses.

**A) Data without imputation:** If neither pair could be imputed, we ran only two analyses: *READv2* (*45*) with pseudo-haploid genotypes and *NGSRelate v2* (*46*) with genotype likelihoods and population allele frequencies (i.e., the standard input of the two programs).

<u>Case A1:</u> The number of common SNPs used in both *READv2* and *NGSRelate v2* is >1000;

i. If both tools had consistent θ estimates (e.g., both in the first-degree range or unrelated range), we assumed this estimate correct.
ii. If the tools returned different results but the assigned relatedness categories were consecutive (e.g., one tool assigning first and the other second-degree), we used fuzzy categories (e.g., ‘First or second-degree).
iii. If the tools returned different results and the assigned relatedness categories were non-consecutive (e.g., second-degree and unrelated), the results were ignored and the pair assigned “NA” for their kinship category.

Case A2: The number of common SNPs used in both *READv2* and *NGSRelate v2* is <1000;

i. Results are ignored, and the pair was assigned “NA” to their kinship category.

**B) Data with imputation:** If one or both genomes in a pair could be imputed, we used three analyses: *READv2* using only non-imputed pseudo-haploid genotypes and *NGSRelate v2* using genotype likelihoods (as above), and also *READv2* using imputed diploid genotypes for one or both genomes (depending on their imputation status).

Case B1: Coverage of the non-imputed genome is >0.01x and the number of common SNPs used in all three analyses is >1000;

i. If all three tests returned consistent θ estimates, we assumed this estimate was correct.
ii. If the θ estimates differed among the tools, we assumed the *READv2* assignment using imputed data to be correct.

Case B2: Coverage of the non-imputed genome is <0.01x and the number of common SNPs of all three tests is >1000;

i. If all three tests returned consistent θ estimates, we assumed this estimate was correct.
ii. If the tools returned different results but the assigned relatedness categories were consecutive and third-degree or closer (e.g., one tool assigning second and the other third-degree), we used fuzzy categories (e.g., ‘second or third-degree’).
iii. If the tools returned different results but the assigned relatedness categories were either ‘third-degree’ or ‘unrelated’, we ignored the *READv2* results based on imputed genotypes and assigned relatedness based on *READv2* on non-imputed genotypes (as in case A1).
iv. If the tools returned different results and the assigned relatedness categories were non-consecutive, the results were ignored and the pair assigned ‘NA’.

Case B3**:** Coverage of the non-imputed individual is <0.01x and the number of common SNPs used in imputed *READv2* is >1000, but the number of common SNPs used in non-imputed *READv2* or *NGSRelate v2* is <1000;

i. If all three tests returned consistent θ estimates, we assumed this estimate was correct.
ii. If results were different but consecutive, and relatedness was third-degree or closer, we used fuzzy categories (e.g., ‘second or third-degree’).
iii. If the tools returned different results and the assigned relatedness categories were non-consecutive, the results were ignored and the pair assigned ‘NA’.
iv. If the tools returned different results and the assigned relatedness categories were ‘third-degree’ or ‘unrelated’, the results were ignored and the pair assigned ‘NA’.

Case B4**:** If both pairs were imputed, we also compared ancIBD results with READv2 results using imputed diploid genotypes.

i. If relatedness estimations are same, we assumed this estimate was correct.

ii. If results were different but consecutive, we used fuzzy categories (e.g., 'second or third-degree').
iii. If the tools gave different results and the assigned relatedness categories were non-consecutive, ancIBD results were ignored and the results were ignored and the pair assigned 'NA'.

<u>Case C:</u> In addition to the genetic data we also checked first-degree connections and observed two cases where the pedigree information is inconsistent with relatedness calculations.

i. The pairs cch131-cch3208 and cch3208-cch331 were first-degree related. All these individuals were subadults, so they had to be siblings. Therefore, cch131-cch331 pairs also had to be siblings; however, relatedness of this pair was originally assigned as 'second-degree', based on rule described in Case B1ii. This relationship was updated to 'first-degree'.
ii. The pairs cch124-cth842 and cth728-cth842 were first-degree subadult siblings, but the relatedness of cch124-cth728 pair was assigned as 'NA' based on rule described in Case A2ii. This relationship was updated to 'first-degree'.

##### v. SNP count threshold based on false positives

We tested kinship among all pairs of individuals irrespective of period. Because the Early and Late periods are ≥200 years in between, close kinship is not expected between pairs from these two periods. Among 685 pairs tested, none were first- or second-degree, while 14 (2%) were assigned third-degree relatedness. All of those pairs likely false positive pairs had overlapping SNP counts <3000. Therefore we further filtered the dataset for pairs with >3000 common SNPs (involving either pseudo-haploid genotypes or imputed genotypes).

#### c. Relatedness estimation based on X chromosome

We estimated the kinship coefficient θ using only X chromosome SNPs to provide further insight into kinship types. Here, we included only pairs with the number of overlapping SNPs >200 and used both *READv2* (*45*) and *NGSRelate* v2 (*46*). These estimates are noisier than autosomal estimates due to the smaller number of SNPs used but can still provide useful information.

#### d. Estimating variation in kinship estimates

We used pedigree simulations (see section 'Simulation of pedigrees') for evaluating the variation in θ estimates. Specifically, we were motivated to determine how frequently siblings could have divergent θ values with an avuncular (second-degree) relative (i.e., $\theta_{sib1\text{-}auntX}$ and $\theta_{sib2\text{-}auntX}$ being different). On average they should have the same θ, but in reality, we might observe substantial variation due to random recombination and noise in θ estimates.

To address this, we defined a pedigree containing four siblings and their avuncular relative in a def file and passed this to *PedSim* with the *'-d'* option and used the simulated founder individuals (see section '**Simulation of founders**' below). The output VCF file was first haploidised with a custom *Python* script named *'haploidize.py'* (https://github.com/MerveNurGuler/Haploidize-VCF) and then with the *PLINK2*(*36*) (v 2.00a4.5) software converted into *PLINK*'s TPED/TFAM format using the *'--maf 0.05 --recode transpose --chr 1-22'* options. After applying MAF and chromosome filters, individuals in the pedigree contained 480,466 overlapping autosomal SNPs. Then, we ran *READ* (*48*) software on 75 replicates of this pedigree to estimate normalized P0 and assigned pairs to relatedness categories. We thus calculated the chance of observing different relatedness category assignments between the four siblings and their avuncular relative. Across the trials, we randomly chose either 10,000, 5,000 and 2,000 autosomal SNPs and repeated the analysis.

Our results in **Figure S31** show that misassignment of an avuncular relationship to first- or third-degree was not uncommon, especially under low SNP numbers, in line with our earlier observations (*44*). This observation was critical for evaluating the genetic evidence for pedigree reconstruction.

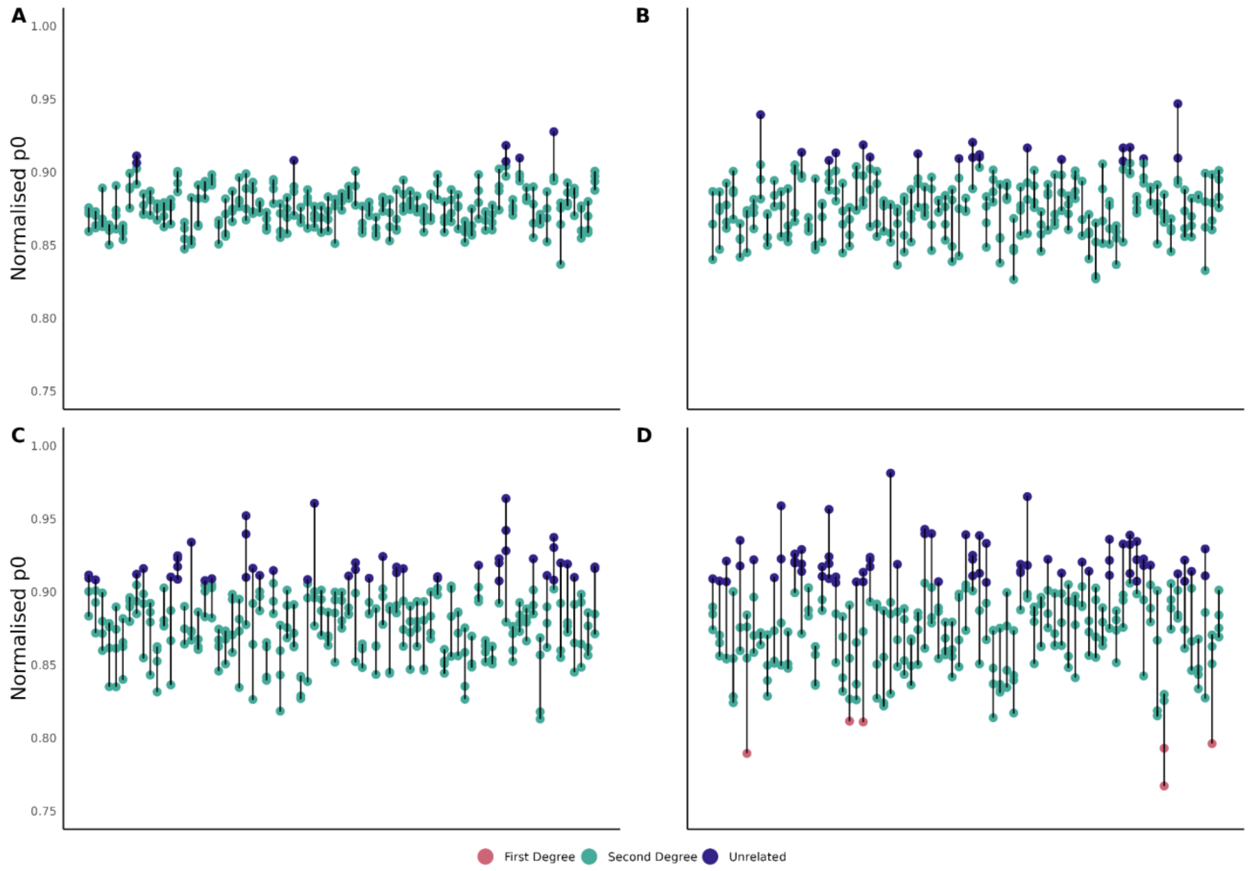


**Figure S31:** The kinship category assignments of four simulated siblings and their avuncular relative using *READ*. Pairs share **A)** 480,466, **B)** 10,000, **C)** 5,000 and **D)** 2,000 overlapping autosomal SNPs. The colours indicate the category assigned by *READ*.

### 11. Runs of Homozygosity (ROH)

We used *hapROH* (*35*), a method designed to call ROH blocks in low-coverage aDNA data by leveraging linkage disequilibrium from a reference haplotype panel. We ran the *hapROH* software with *'e_model="haploid", p_model="EigenstratUnpacked", n_ref=2504, random_allele=True, readcounts=False, delete=False, logfile=True, combine=True'* parameters. This method is applicable to shotgun genomes with >0.3x coverage, and it can successfully recover ROH blocks >4 centimorgan (cM) on the 1240K SNP set. When offspring have close parental relatedness in their history, the program can detect long ROH at even lower coverage. Here, we called ROH across 16 newly produced Çatalhöyük genomes with >240K SNPs (mean c.500k). We also called ROH among published genomes from Neolithic Anatolia with similar coverage (**Table S2**).

### 12. Genealogy simulations for measuring inbreeding under random mating

#### a. Simulation framework

Our goal here was to estimate the accumulation of inbreeding under different population sizes and breeding behaviour patterns within a certain time frame, under random mating. For this, we developed a *Python* script that takes breeding female and male numbers and the number of generations as parameters and simulates a randomly mating population, while recording genealogical relationships over multiple generations.

- Our simulations begin by creating a male and a female founder pool based on user-specified female and male numbers. This may be 1:1 or can be skewed.
- The first generation is formed randomly from these pools, keeping population size constant and assuming non-overlapping generations.
- Subsequent generations are formed from the previous generation with the same logic.

To make the simulation more realistic, we controlled the mating process using additional parameters:

- We allow the user to define the mean number of children per couple and the maximum number of children per individual. Each mating pair's offspring number (i.e., number of full siblings in the next generation) is drawn randomly from a Poisson distribution. The mean number of children per couple parameter of the Poisson was set to 2 in our simulations. Also, for each individual, we determined a limit for the number of offspring, a random value drawn from the Poisson distribution with a mean specified in the mean maximum number of children per individual parameter, set to seven in our simulations.
- We manipulated to exclude sibling matings. We did this because sibling matings will be highly rare in reality, but if they happen frequently in simulations under small population size, they could have a disproportionate impact on simulation inbreeding estimates.

The resulting genealogical relationships are stored in a data frame to trace lineage and calculate the pedigree-based inbreeding coefficient, $F_{PED}$, for any individual.

$F_{PED}$ for last-generation individuals are calculated following Ballou's description (*49*), backtracing the lineages and counting the loops. The code for this calculation is available at https://github.com/BusraKatircioglu/inbreedingSimulation.

Due to the exponential increase in the algorithm's time complexity with generation depth, we set a generation limit for backtracking. We trace back up to the 15th generation, except for the loop contributed by the first common ancestor of the mother and father lineages. We checked the effect of this limit and concluded that for population sizes >1000, this limit does not change the $F_{PED}$

distribution (**Figure S30**). For lower population sizes (e.g., 300) our algorithm is slightly underestimating $F_{PED}$.

#### b. Implementation for comparison with observed data

We ran our script according to the parameters in **Table S20**. For each set of parameters we performed 5 trials, and in each trial we randomly sampled 16 individuals from the last generation 1000 times. We then calculated the mean, median or maximum $F_{PED}$ values for these samples, and plotted the distributions to compare with the $F_{ROH}$ estimates from the 16 Çatalhöyük genomes. We note that previous has shown that $F_{ROH}$ and $F_{PED}$ are comparable statistics(*50*). We also note that the possible underestimation of $F_{PED}$ in our simulations due to the generation limit (**Figure S32)** would not alter our main conclusion, that either a large breeding population or active inbreeding avoidance would be necessary for the observed $F_{ROH}$.

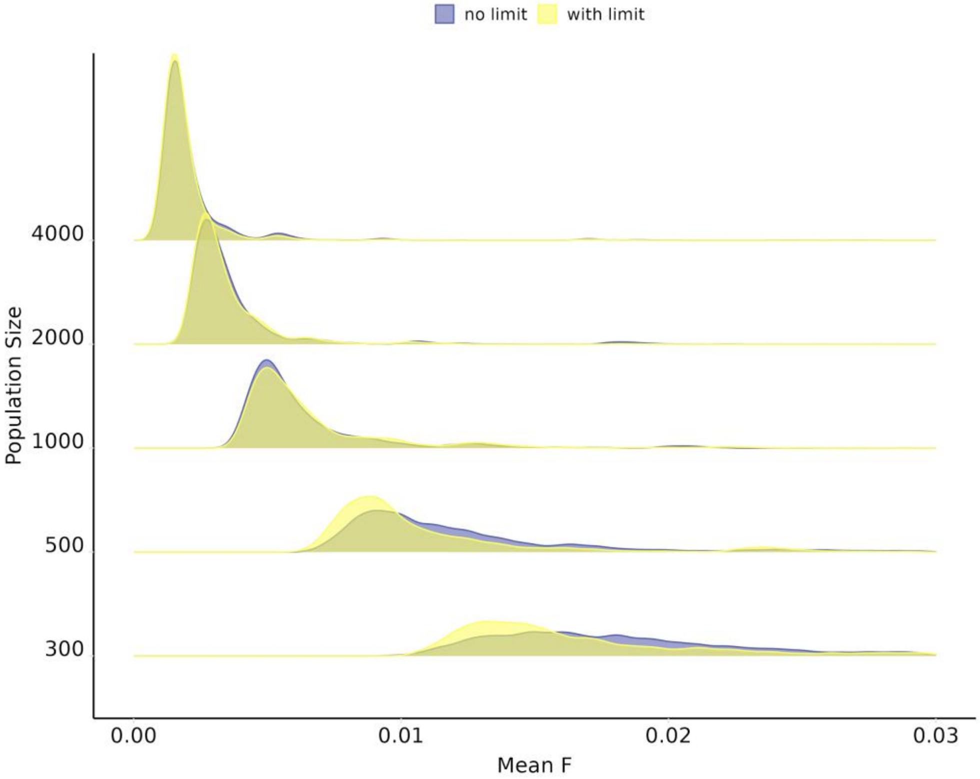


**Figure S32:** Effect of generation limit on simulated inbreeding coefficient distributions. We ran the algorithm 5 times for each distribution with the same parameters, taking the 1000 subsets with 16 elements each time and calculating the mean of each.

### 13. Uniparental haplogroup homogeneity / diversity analyses

#### a. Determining major haplogroups

Here our goal was to use mtDNA and chrY haplogroup data to infer maternal and paternal connections, respectively. However, due to limited data coverage, some individuals who actually should share the same mtDNA or chrY haplogroup subclade (such as siblings or parent-offspring pairs) might actually be assigned to either a more specific or more general haplogroups, thus confounding our analysis. Additionally, there is a possibility of incorrect assignment to a different subclade due to post-mortem DNA damage (C to T and G to A transitions).

To address this, we decided to use the first two characters of mtDNA and chrY haplogroups (ISOGG format) when calculating haplogroup homogeneity. mtDNA haplogroup H and its subclades were reclassified under the upper clade haplogroup HV. Additionally, samples with single-character haplogroups were excluded from the analysis to ensure consistency.

We note that using high level clades will naturally lead to some pairs being assigned the same haplogroup even though they carry different mtDNA or chrY haplotypes. Still, because our haplogroup-based analyses involve comparisons among groups, we do not expect this to bias our results.

#### b. Haplogroup homogeneity calculations

To calculate haplogroup homogeneity (or its inverse, diversity) within a set of individuals, we collected all pairs of samples that have an assigned haplogroup. We then filtered these pairs based on the specific criteria being inspected. For **Figure 2E**, in Çatalhöyük, we selected pairs from the same period, while in Gurgy (see below), we did not apply any filters. Relatives were *not* excluded in either case.

For **Figure 5A**, we selected pairs from the same period and filtered them by those within the same building and between different buildings. We calculated homogeneity as the ratio of pairs in which both individuals have the same major haplogroup to the total number of pairs. Diversity was calculated as 1 minus homogeneity.

When calculating homogeneity within buildings, we computed it separately for each building, and then averaged these homogeneity values among buildings from the same period. Directly calculating the homogeneity from individual pairs within the same building in the same period would introduce bias due to the varying number of individuals in different buildings. For calculating homogeneity between buildings, we calculated average the homogeneity among pairs between each pair of buildings, and then averaged these values to obtain the overall homogeneity measure.

#### c. Haplogroup assignment in the Gurgy dataset

We downloaded haplogroup information for Gurgy, a Middle Neolithic settlement from France (*51*), to compare mtDNA and chrY haplogroup homogeneity. For Y-chromosome haplogroups, we used data from Supplementary Table 7 ("YC assignment"). For mtDNA haplogroups, we referred to Supplementary Table 5 ("Mitodata") from the same article (*51*).

### 14. Simulations of matrilocal, patrilocal and mixed residence

#### a. Simulation of founders

We simulated the genomes of 250 founder individuals using *stdpopsim* (*52*, *53*) software to create pedigrees for two different analyses. We utilized the *msprime* (*42*) engine in the *'HomSap'* mode from the stdpopsim library to simulate genotypes of founder individuals. We used the *'HapMapII-GRCh37'*(*54*) with the '-g' option as an empirical recombination map. We thus produced 500 haploid genomes using a multi-population demographic history model of ancient Eurasia (*55*) with the *'-d AncientEurasia-9K19 0 500'* option. This is a simplified representation of the European Neolithic Linearbandkeramik (LBK) population of Anatolian descent (*56*), and thus a reasonable proxy for Çatalhöyük.

Next, we converted the succinct tree sequence output from *stdpopsim* into VCF format using the *tskit* (*43*) library *'vcf'* command with the *'--ploidy 2'* option. We selected 1 million random SNP positions with a custom bash script. These positions were further used to extract reference bases from the human reference genome (hs37d5) using the *'getfasta'* command of *BEDtools* (*10*) (v2.27.1). We estimated the transition/transversion ratio from the 1000 Genomes (*19*) Dataset v3 Tuscany (TSI) population to assign alternative alleles to the reference positions. With this information, we stochastically generated alternative alleles for each position in our dataset, employing a customized *R* script. This approach allowed us to replicate genetic variation according to the observed rates in the TSI population, resulting in a realistic distribution of allele frequencies in our simulated dataset.

#### b. Simulation of pedigrees

We used the VCF with simulated founder individuals, which is phased and contains no missing sites, to create pedigrees with the *PedSim* (*57*) pedigree simulator using the *'-i'* option. Additionally, we provided the randomly assigned sexes of the input VCF samples with the *'--sexes'* option. To create a map file, we interpolated the positions in the VCF file containing 250 simulated founder individuals using the '*filter_vcf.py'* script from the *adna_tools Python* package (https://github.com/CompEvoMetu/adna_tools). We then used this map file in the pedigree simulation step with the '-m' option. We employed the crossover interference model (*58*) provided by *PedSim* software to create more realistic genomes, using the *'--intf'* option while running *PedSim*. Additionally, we used the *'--keep_phase, --founder_ids, --fam, and --miss_rate 0 --X X'* parameters when running *PedSim*, along with a def file containing information on pedigree architecture specified by the *'--d'* option.

We then defined three pedigree scenarios described in **Figure S33** in def files and passed this to *PedSim* with the *'-d'* option. The output VCF file was first haploidized with a custom *Python* script and used in residence simulations (see section "Residence simulations type simulations for autosome vs chrX diversity").

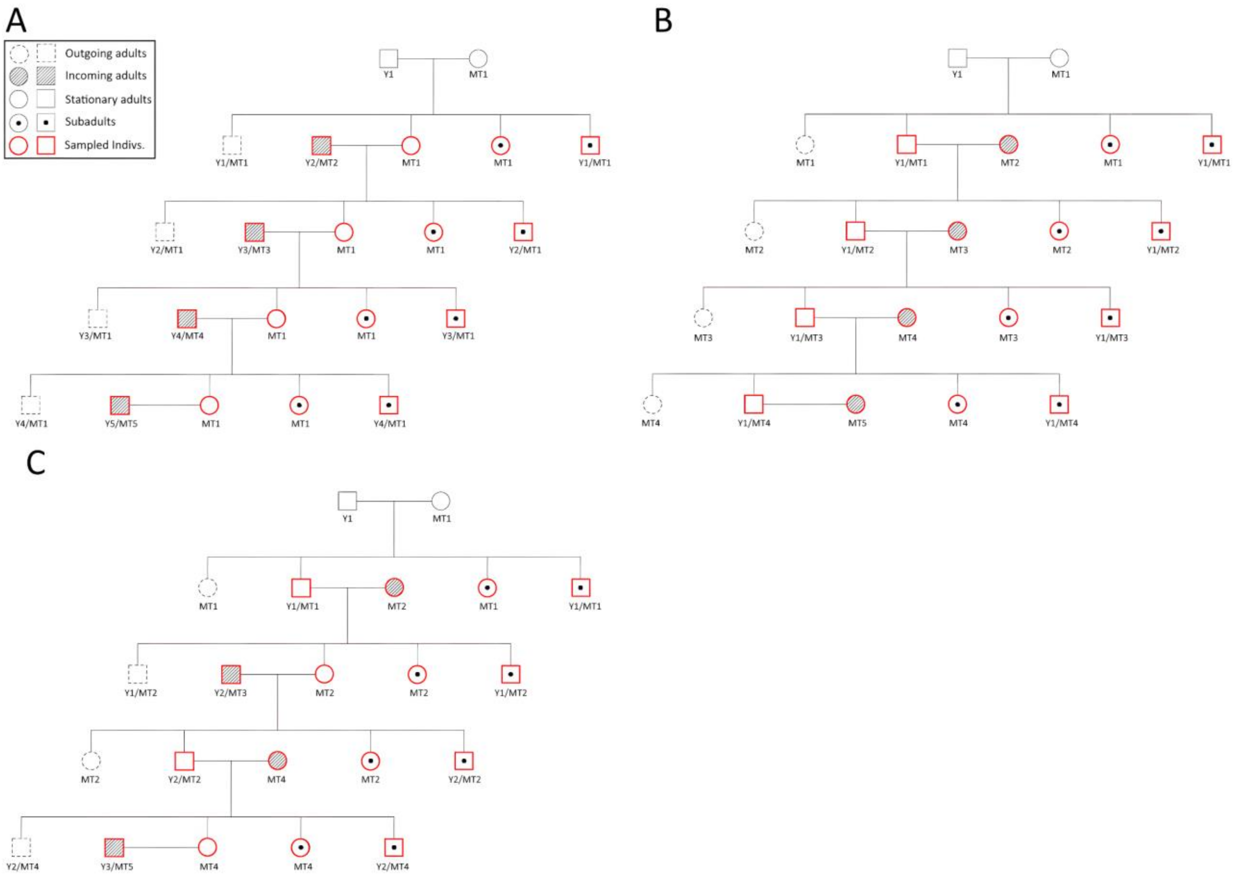


**Figure S33:** Three residence type scenarios simulated: **A)** matrilocal residence, **B)** patrilocal residence, **C)** mixed residence. The individuals sampled in the simulations are shown in red. Outgoing and incoming adults refer to individuals leaving or joining the pedigree. Note that in the simulations we assume a single biological family occupying a building here.

#### c. Residence type simulations for haplogroup diversity

In these simulations we created multi-generational families occupying buildings (one family per building), which followed matrilocal, patrilocal or mixed residence practices.

We used the Çatalhöyük mtDNA and chrY gene pools as starting point, and *not* the *PedSim* genomic simulation results described above. For this, we selected unique haplogroups for all genetically sampled Çatalhöyük buildings for either uniparental marker. We created founder gene pools using these sets (using unique haplogroups per building ensured that some buildings were not overrepresented).

We then created 100 pedigrees, representing the multi-generational inhabitants of Çatalhöyük buildings, and randomly assigned a mtDNA and chrY haplogroup as a founder to the initial generation of these buildings. For subsequent generations, haplogroups were inherited from the biological parents. Depending on the residence type, we shuffled adults between buildings in each generation (**Figure S30**). For example, in matrilocal residence, while adult females stayed in the building, adult males moved between buildings to mate. The subadults remained in the building (representing individuals who died before reproductive age).

We generated 100 simulation datasets for each residence type (patrilocal, matrilocal, mixed) with the following configurations:

- 2 adults and 2 subadults per generation, with each building occupied for 4 generations.
- 2 adults and 2 subadults per generation, with each building occupied for 6 generations.
- 2 adults and 4 subadults per generation, with each building occupied for 4 generations.
- 2 adults and 4 subadults per generation, with each building occupied for 6 generations.

We thus created 400 total simulations per residence type. We then calculated the proportion of pairs with the same haplogroup within buildings for both mtDNA and chrY haplogroups in each simulation, and returned the mean of this proportion.

#### d. Residence type simulations for autosome vs chrX diversity

For the autosome and chrX theta difference, we used 100 simulation datasets (VCF) generated simulated by *PedSim* under three of the residence types (patrilocal, matrilocal, mixed). The simulations contained 15 multi-generational pedigrees, which again stand for inhabitants of Çatalhöyük buildings, with 2 adults and 4 subadults per generation. Each building was occupied for 6 generations. We haploidized these VCF files with a *Python* script, converted them to *PLINK* format (.bed, .bim, .fam), applied a >0.01 MAF filter, and separated the datasets into autosome and X chromosome genotypes with PLINK. Then we ran *READv2* on the data. We sampled pairs according to the following configurations:

- 2 adults and 2 subadults per generation, with each building occupied for 4 generations.
- 2 adults and 2 subadults per generation, with each building occupied for 6 generations.
- 2 adults and 4 subadults per generation, with each building occupied for 4 generations.
- 2 adults and 4 subadults per generation, with each building occupied for 6 generations.

We thus created 400 total simulations per residence type. We then calculated the difference between autosome theta and chrX theta for pairs within buildings for each simulation, and then took the mean of these differences.

#### e. Modelling Çatalhöyük residence dynamics using simulation results

We thus collected 400 simulation results under each residence type, each simulation providing three summary statistics: a) mean chrY homogeneity, b) mean mtDNA homogeneity, and c) mean autosomal vs. chrX theta differences. For each statistic, we averaged the results of the 400 simulations.

Meanwhile, for the empirical data, we calculated mtDNA and chrY haplogroup homogeneity within buildings for each period of Çatalhöyük and also averaged these values. We calculated the theta difference between autosome and X chromosome within buildings only for the Middle period, as there were few pairs with >250 overlapping SNPs on chrX in the other periods.

Then, for each summary statistic, we normalised the set of observed values and three simulation values between 0 and 1 to assign all statistics equal weight in the next step (the table shown in **Figure 5D**).

Finally, we calculated all possible (n=5151) weighted averages (across the percentile range) of the mean summary statistics and determined the Euclidean distance between each vector of weighted averages and the observed values:

$$Distance = \sqrt{\left( \sum (Observed - (w_1 * Patrilocal + w_2 * Matrilocal + w_3 * Mixed))^2 \right)}$$

where *Observed* is the normalized vector of the three summary statistics observed, while *Patrilocal*, *Matrilocal* and *Mixed* are the normalized vectors of the mean summary statistics estimated from the simulations, and $w_1$, $w_2$, $w_3$ are the percentile weights of the three residence type models. The results are shown in **Figure 5C**.

### 15. Burial and diet analyses

#### a. DNA preservation differences

We analyzed 411 samples in terms of endogenous aDNA proportion (**Table S1**). We converted human endogenous DNA proportion per individual (using merged libraries per individual) to logarithmic scale. We then compared these values across different categories: the 3 main periods (Early, Middle, Late), the areas of the settlement (North and South), buildings (49 buildings in total), burial location, deposition of burials, flexing of burials, age groups, and sex groups. We also performed similar analyses using data from other Neolithic settlements (Aşıklı, Çayönü, Gurgy). For the deposition of burials, there were 4 main groups: primary, primary disturbed, secondary, and tertiary burials. Primary burials were considered as an undisturbed state, and others were disturbed states. For age groups, all adults were grouped together and subadults included adolescent, child, infant, neonate, and prenatal age groups. We required a minimum of >3000 PMD-filtered reads for reliable sex assignment for a genome. For Aşıklı and Çayönü, unpublished genomes generated in screening experiments in Ankara were also used **(Table S21).**

#### b. Burial objects

Data on Çatalhöyük burial objects associated with primary deposited burials has been previously collected and published by Vasic and colleagues (*59*) (**Table S15**). Here we used this data to study the frequency of the presence of burial objects (versus absence), compared across age and sex groups. For the sex assignments, we used the $K_{XY}$ assignments and a total read filter of >3000 PMD-filtered reads (see section **'Molecular sex assignment'**).

#### c. Dietary isotopes

Bulk collagen isotope values of carbon (δ13C) and nitrogen (δ15N) values represent protein or plant consumption of individuals. High δ15N values indicate the trophic level from which the proteins were acquired, while variation in δ13C levels can inform on differences in plant consumption (*60*). We focused our analysis on stable dietary isotope values from neonates in the Late period of Çatalhöyük, which was the largest sample of subadults in a single period. Neonate dietary isotopes are expected to reflect duration of nursing, but may also be impacted by the diet of the mother or wet nurse, the diet of the mother during pregnancy, or catabolic processes in the case of malnutrition (*61*). We studied the diets of 13 neonates excavated from six different buildings (neonates from five of those buildings were genetically sampled). Only buildings 53, 58 and 65 contained genetically related pairs. That is also supported by the fact that we observed statistically significant differences among buildings (Kruskal-Wallis test $p<0.02$).

### 16. Simulations of within-building co-burial relatedness

A series of Monte Carlo simulation tests were performed to test whether the observed temporal decrease in within building co-burial relatedness frequencies, i.e. increasing frequency of unrelated co-burials (**Figure 3**), could be caused by technical factors such as differences in burial numbers and/or genetic sample sizes rather than being a genuine effect. For this, we simulated kinship frequencies under the null hypothesis of no difference among periods in co-burial kinship, given the observed sample sizes and age structure, and assuming a range of biological family sizes using each building.

We selected n=23 buildings for the simulations: buildings dated to specific Çatalhöyük periods and that had at least two burials used in kinship analyses **(Table S22**). Each building was assumed to be composed of groups that are genetically related to each other (simply referred to as “families”) as well as individuals who are genetically unrelated to the rest **(Figure S34)**. The total number of individuals per building (the number of “inhabitants”) was set as the number of burials excavated in that building. Briefly, we randomly chose the same number of individuals from the inhabitants as genetically sampled from that building, and recorded their relatedness frequency; we eventually compared it with the observed data.

Two different settings were used for the simulations. For the first setting, families were assumed to have a constant number of members (referred as family size) for each building. Family sizes of 2-6 were separately tested for this setting. For the second setting, family sizes were assumed to vary in each building, a random size was assigned to each building from an array of 2-6 for this setting. In total, 6 different simulations were performed (5 for constant size and 1 for variable size). The setup of the simulations is described below step-by-step with a schematic description **(Figure S34).** The Early period building 6 (B6) and the first setting with the family size of 4, is given as an example:

1. In total, 11 burials had been identified in B6, and therefore we assigned 11 inhabitants to B6 **(Figure S34A)**.

2. Families were formed inside each building by assigning the inhabitants to groups of certain number and the rest as unrelated. In our example with a family size of 4, in B6, we first assigned 8 individuals to 2 families, and the remaining 3 as unrelated **(Figure S34B)**.
3. We assigned different age categories (adult, subadult or adolescent) to the inhabitants based on the real number of burials within that age category. In B6, one individual was an adolescent, 2 adults, and the remaining 8 subadults (**Figure S34C)**.
4. A subset of these individuals were randomly selected for further analyses based on the number of burials used in genetic kinship analyses from the building. In B6, 6 subadults were thus randomly selected **(Figure S34D)**.

Steps 1-4 were repeated 100 times. A custom *Python* script *'samplecoburiedkin.py'* was used to compute these steps.

5. Relatedness for all possible pairs collected were listed, and some pairs were randomly selected based on the number of pairs that have estimated relatedness in observed kinship analysis (given some pairs in the kinship dataset not having sufficient data to call kinship). In B6, all pairs had sufficient data, and therefore we used all 15 subadult-subadult pairs.
6. We calculated the odds ratio (OR) for relatedness frequencies between Early-Middle, Middle-Late, and Early-Late period samples (from 2x2 tables of related/unrelated x Period1/Period2).
7. The OR values were calculated using all pairs, and also for only subadult-subadult pairs.

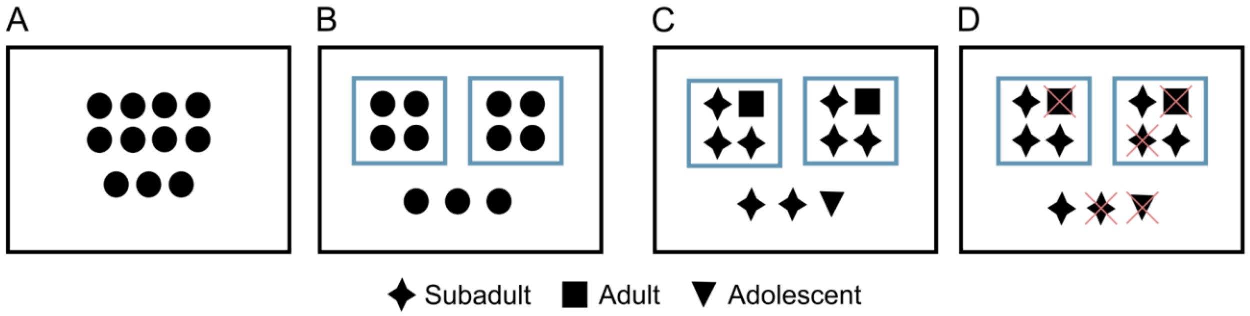


**Figure S34:** Schematic description of the simulation setup of the constant family size of 4 in building 6 (B6). **A)** Assignment of individuals to B6. **B)** Creating genetic families. **C)** Assignment of age categories following the real age distribution of B6 burials. **D)** Random sampling of individuals, as many as sampled for kinship analyses from B6. Blue rectangles show genetic families and red crosses show individuals removed after random sampling.

Steps 5 and 6 were repeated 1000 times. Finally, 100,000 (100 x 1000) OR values were obtained from simulations for each setting. A custom *Python* script *'randomizecoburiedkin.py'* was used to compute steps 5 and 6.

OR values for observed relatedness frequencies between Early-Middle, Middle-Late, and Early-Late periods were calculated for three different age-classifications; all-samples, subadult-subadult pairs including adolescents, and subadult-subadult pairs excluding adolescents.

The statistical significance of these observed OR values between any pair of periods was computed by comparing these with the corresponding simulated OR distributions (for the same period pair). A custom *Python* script *'calculateOR.py'* was used to compute the simulated OR distributions.

### 17. Metagenomic analysis

#### a. Microbial screening

The metagenomic screening of n=664 libraries was done by following the *aMeta* (*62*) pipeline with modifications explained below. The databases utilized for the *KrakenUniq* and *MALT* steps were consistent with those of aMeta, comprising non-redundant NT/NR records from June 2020. These databases encompass all microbial organisms (archaea, bacteria, viruses, fungi, protozoa, parasitic worms), the human reference genome, and a selection of complete eukaryotic genomes.

The first step includes pre-screening of present microbial species using *KrakenUniq*(*63*). *KrakenUniq* results for each library were filtered by 1000 unique *k*-mers and 200 unique reads by using in-house *R* script.

Following this, we created a database for *MALT* using the *malt-build* program in the *MALT*(*64*) software (v 0.6.2) by extracting a subset of the pre-built database containing only the unique species detected across all libraries by *KrakenUniq*. Then, using an in-house bash script, we generated an abundance matrix by dividing the number of reads assigned to each specific taxon within a library by each library's total number of assigned reads. The *KrakenUniq* abundance matrix for 664 libraries (including 28 libraries prepared from teeth) are summarized in **Table S23.** Subsequently, each library's FASTQ file was aligned against the *MALT* database using the *malt-run* program with the options '*-at SemiGlobal'* and '*-m BlastN'* in the *MALT* software (Huson et al., 2007). For each library, microbial reads uniquely assigned to respective taxa were extracted via the *MaltExtract* (https://github.com/rhuebler/MaltExtract) (v 1.7) program with *'--destackingOff, --downSampOff, -dupRemOff, -minPI 85.0, --maxReadLength 0 --minComp 0.0'* parameters. Finally, authentication metrics such as depth and breadth of coverage, PMD and edit distance profiles were calculated following the *aMeta* pipeline (*62*).

Overall, the detected microbes in the 664 libraries likely have an environmental rather than endogenous origin, particularly microbes that thrive in the soil. This result was expected, as almost all libraries (~95%, 636 out of 664) were prepared from petrous bone. As the inner part of the temporal bone, especially the cochlea lacks capillaries, petrous bone libraries are mostly devoid of human pathogens that circulate in the blood, unlike tooth libraries. Moreover, during the taphonomic process, soil can enter the ear canal, and soil microbial DNA can remain in the DNA extract. However, we did observe interesting patterns: Two petrous-derived libraries, cch253 and cch175, also contained reads with ancient profiles (i.e. with PMD) mapping to n=6 species of microbes with >10% breadth of coverage. In cch175, *Clostridium perfringens*, *Staphylococcus equorum*, and *Clostridium tetani* were detected with 3.29x, 0.58x, and 0.19x depths of coverage and 76%, 42%, and 16.5% breadths of coverage, respectively. In cch253, *Clostridium tetani*, *Alcaligenes faecalis*, *Clostridium butyricum* and *Clostridium perfringens* were detected with 1.43x, 0.67x, 0.17x and 0.11x depths of coverage and 68.5%, 46.7%, 14.8% and 10% of breadths of coverage, respectively. However, these microbes are highly unlikely to be associated with the deaths of these individuals given their ubiquity. Sequence statistics for all *MALT* alignments and authentication plots of authentic microbes can be found in **Table S24**.

#### b. Testing differences in bone preservation-associated microbes using random forests

Our finding of age-related organic preservation differences in skeletons (**Figure 6A-B**) may be attributable to differential burial treatment. We hypothesized that such treatment and/or preservation differences may be reflected in the metagenomic profiles.

To test such a possible association between organic preservation and microbial composition, selected libraries with divergent human endogenous DNA proportions and tested for differences in *k*-mer profiles and microbial abundance profiles between the groups. For this, we first chose 40 libraries that consisted of 10 adult (0.13x to 0.02x) and 10 subadult libraries (0.58x to 0.16x) with the highest endogenous DNA proportions, and 10 adult (0.0005x to 0.0003x) and 10 subadult libraries (0.00017x to 0.0001x) with the lowest proportions.

Before analyzing *k*-mer profiles, we filtered out reads mapped to the human reference genome from their respective FASTQ files to avoid confounding by human DNA. First, we retrieved IDs of reads that mapped to the human reference genome from unfiltered bam files using *SAMtools* (*9*) (v 1.18) with the '*view*' option. Then, we retrieved IDs of all sequenced reads from FASTQ files using *seqtk* (v 1.3) with the *-a* option. Using grep with the *'-v'* option in bash, we extracted IDs of reads that were not mapped to the human reference genome. Finally, we generated versions of each FASTQ file containing only non-human reads using *'seqtk'* with the *'subseq'* option.

*K*-mer profiles were calculated for each FASTQ file using *Jellyfish* (*65*) software (v 2.3.0) with a *k*-mer length of 15. An absolute abundance matrix was created by merging *k*-mer profiles for each library using an in-house bash script. Overall, a total of 536,476,361 unique *k*-mers were detected across the 40 libraries. Since the total number of unique kmers was unsuitable for analysis, we randomly selected 5 million *k*-mers using the '*powershuf.py*' (https://gitlab.com/aapjeisbaas/shuf) script. Subsequently, we calculated a relative abundance matrix by dividing the respective *k*-mer count with the total number of *k*-mers for each library using an in-house *R* script.

To test for the presence of distinctive *k*-mers that could be associated with human DNA preservation, we used a random forest approach (*66*). This method randomly selects features to create several decision trees to predict an outcome, then combines those predictions through a voting approach. This type of method enables to reduce variance and increase the prediction performance. In essence, each classification tree is trained on only a part of the data and the remaining part is used to calculate an out-of-bag (OOB) error rate. In this way, random forests internally divide the dataset as training and test data, and output a generalized OOB error rate. To rank the importance of the features, we used the mean decrease in Gini impurity (MDG) index that shows how each feature contributes to the overall accuracy of the model.

Libraries with human endogenous proportions >0.01 were classified as "High Proportion" (n=20), while those with proportions <0.0005 were classified as "Low Proportion" (n=20). Then, we used *randomforest* function with *ntree=1000* parameter on *k*-mer relative abundance matrix in randomforest *R* package (*67*).

The classification conducted by the random forest and the top 25 important *k*-mers are summarized in **Figure S35**. The overall OOB error rate was estimated around 5%. Only 2 out of 20 libraries with a high human proportion were misclassified as low proportion and only 1 out of 20 low human proportion libraries were misclassified as high proportion. The random forest approach identified *k*-mers that differ between high and low human proportion libraries. However, the top 25 *k*-mers with the highest MDG indexes had values close to zero. These low Gini scores suggest that individual *k*-mers have a weak signal for classification, i.e. no single *k*-mer is particularly powerful in distinguishing between libraries with high and low human endogenous DNA **(Table S25).**

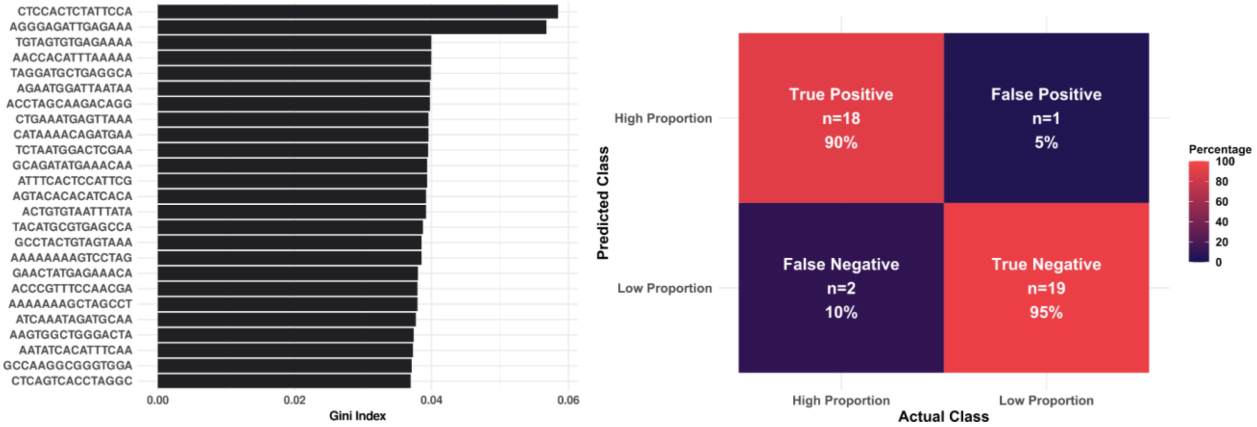


**Figure S35:** Random forest analysis conducted on the *k*-mer relative abundance matrix for distinguishing 20 high and 20 low human DNA proportion Çatalhöyük libraries. The top 25 important *k*-mers with the highest Gini indexes are summarized in the left panel, while the random forest group predictions are noted in the right panel.

To investigate differences in metagenomic profiles between high and low human proportion libraries, we used *KrakenUniq*(*63*) (v 1.0.4). We run Principal Component Analysis (PCA) using the *rda* function in the *R 'vegan'* (*68*) package based on the *KrakenUniq* Abundance Matrix. The PCA plot is summarized in **Figure S36**. In the PCA plot, high human proportion libraries clustered tightly, while low human proportion libraries were scattered.

We also performed a random forest analysis with the *KrakenUniq* abundance matrix **(Table S26).** The classification conducted by the random forest and the top 25 important microbial species are summarized in **Figure S37.** The overall OOB error rate was estimated at 25%. Out of the 20 libraries with a high human proportion, 7 were misclassified as low proportion, while 3 out of the 20 low human proportion libraries were classified as high proportion. The random forest approach identified microbial species that differ between high and low human proportion libraries. Moreover, PCA loadings towards high human proportion libraries are also compatible with the most important microbes found in the random forest analysis.

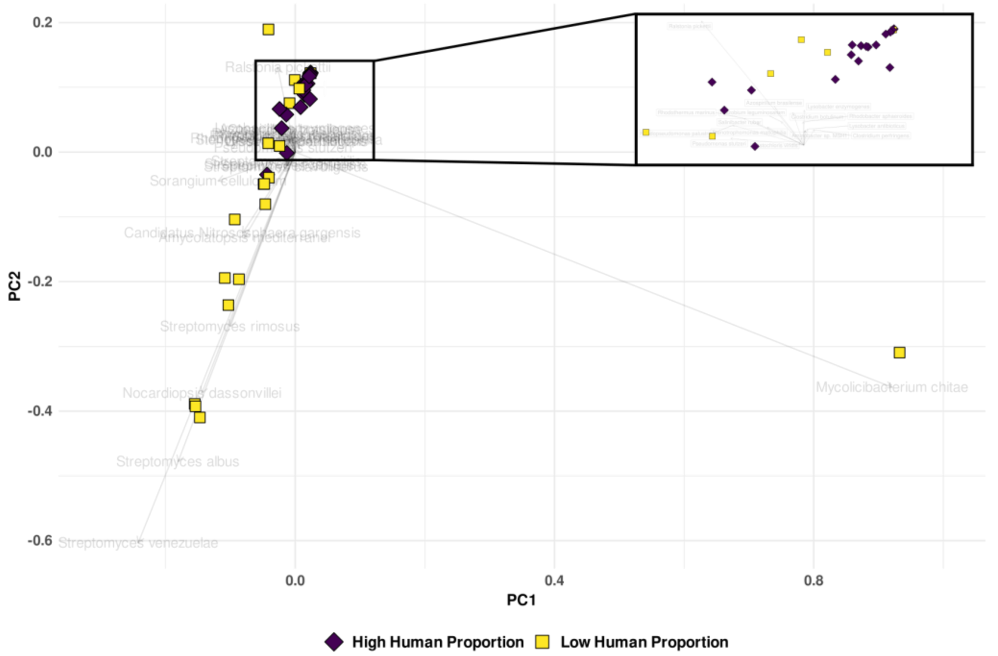


**Figure S36:** Principal Component Analysis (PCA) on the KrakenUniq abundance matrix includes the libraries with the highest (n=20) and lowest human proportion (n=20). The top right side of the figure zooms in on the high human proportion libraries. The top 25 loadings with the highest length are shown in the background.

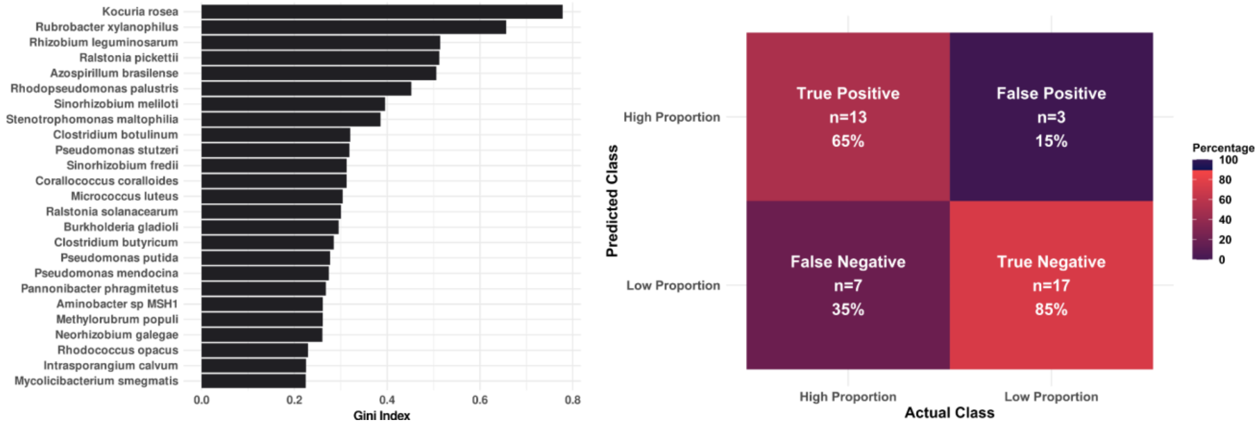


**Figure S37:** Random forest analysis was conducted on the *KrakenUniq* relative abundance matrix. The top 25 important microbial species with the highest Gini indexes are summarized in the left panel, while the random forest group predictions are noted in the right panel.

#### c. Testing aerobic versus anaerobic microbe abundance differences

Finally, we hypothesised that if low-preservation is indeed related to drying, defleshing or similar funerary treatments of the body (as we argue in the Main Text), such treatments would expose the body to higher levels of oxygen not available during decomposition in a grave, which in turn is mainly driven by anaerobic microbes. Indeed, an experiment with surface exposed and buried body parts reported environmental oxygen availability strongly shaped microbial communities that colonise the bone, with anaerobic gut bacteria more prevalent in deep buried body parts (*69*, *70*). We therefore tested whether aerobic microbes may be more prevalent in low-preserved libraries.

To do this, we first gathered oxygen tolerance information for the microbial species detected in the *KrakenUniq* analysis (see section '**Microbial screening**') from the BacDive database (https://bacdive.dsmz.de/) (*71*). We further collected all Çatalhöyük merged libraries (n=363) and sorted them into "High Proportion" (n=95) and "Low Proportion" sets (n=225), depending on whether the human proportion was >1% or <1%. After that, we filtered out the libraries containing only anaerobic or aerobic microbial species. We then calculated the overall proportion reads assigned to aerobic or anaerobic microbial species in each library. Interestingly, aerobic microbe proportions were >10-fold higher than anaerobic microbes among the "Low Proportion" libraries (median 0.102 vs. 0.009), while in "High Proportion" libraries anaerobic microbes were 20% higher (median 0.004 vs. 0.005). The results are summarized in **Figure S19** and **Table S27** below**.** We then tested these differences using both the paired t-test and Wilcoxon signed rank test, with the *t.test* and *wilcox.test* functions in the '*stats'* package in *R*, respectively, with the *'paired=T'* option. This supported the higher abundance of anaerobic versus aerobic microbes in "High Proportion" libraries (paired t-test p=0.356, Wilcoxon signed rank test p=0.082), as well as the higher abundance of aerobic versus anaerobic microbes in "Low Proportion" libraries (paired t-test p=3.90E-13, Wilcoxon signed rank test p=9.94E-12).

We also repeated these analyses by retaining only libraries that contained both anaerobic and aerobic microbial species. As a result, the sample sizes for "High Proportion" and "Low Proportion" libraries decreased by 93 and 131, respectively. Running the same analyses, we again found significant differences in abundance in "High Proportion" (paired t-test p=0.297, Wilcoxon signed rank test p=0.032) and "Low Proportion" libraries (paired t-test p=0.0042, Wilcoxon signed rank test p=0.0137) (**Figure S19**).

Our findings support the idea that microbial profiles may differ between highly and lowly preserved bone material. Moreover, the finding of a higher frequency of aerobic species in low-preserved bones is consistent with the notion of a longer duration of exposure of the bodies before burial in Çatalhöyük, likely as part of a specific funerary treatment. That said, we acknowledge that the observed correlations between human DNA preservation and microbial content might be driven not only by differences in decomposition dynamics, but also porosity differences between the two groups of bones (high- versus low-preserved) that allow different microbes to colonise the bones at later stages, including post-excavation. Understanding the exact mechanisms behind the observed correlations thus need further investigation, such as controlled body farm experiments.

19. A. Auton, G. R. Abecasis, D. M. Altshuler, R. M. Durbin, G. R. Abecasis, D. R. Bentley, A. Chakravarti, A. G. Clark, P. Donnelly, E. E. Eichler, P. Flicek, S. B. Gabriel, R. A. Gibbs, E. D. Green, M. E. Hurles, B. M. Knoppers, J. O. Korbel, E. S. Lander, C. Lee, H. Lehrach, E. R. Mardis, G. T. Marth, G. A. McVean, D. A. Nickerson, J. P. Schmidt, S. T. Sherry, J. Wang, R. K. Wilson, R. A. Gibbs, E. Boerwinkle, H. Doddapaneni, Y. Han, V. Korchina, C. Kovar, S. Lee, D. Muzny, J. G. Reid, Y. Zhu, J. Wang, Y. Chang, Q. Feng, X. Fang, X. Guo, M. Jian, H. Jiang, X. Jin, T. Lan, G. Li, J. Li, Y. Li, S. Liu, X. Liu, Y. Lu, X. Ma, M. Tang, B. Wang, G. Wang, H. Wu, R. Wu, X. Xu, Y. Yin, D. Zhang, W. Zhang, J. Zhao, M. Zhao, X. Zheng, E. S. Lander, D. M. Altshuler, S. B. Gabriel, N. Gupta, N. Gharani, L. H. Toji, N. P. Gerry, A. M. Resch, P. Flicek, J. Barker, L. Clarke, L. Gil, S. E. Hunt, G. Kelman, E. Kulesha, R. Leinonen, W. M. McLaren, R. Radhakrishnan, A. Roa, D. Smirnov, R. E. Smith, I. Streeter, A. Thormann, I. Toneva, B. Vaughan, X. Zheng-Bradley, D. R. Bentley, R. Grocock, S. Humphray, T. James, Z. Kingsbury, H. Lehrach, R. Sudbrak, M. W. Albrecht, V. S. Amstislavskiy, T. A. Borodina, M. Lienhard, F. Mertes, M. Sultan, B. Timmermann, M.-L. Yaspo, E. R. Mardis, R. K. Wilson, L. Fulton, R. Fulton, S. T. Sherry, V. Ananiev, Z. Belaia, D. Beloslyudtsev, N. Bouk, C. Chen, D. Church, R. Cohen, C. Cook, J. Garner, T. Hefferon, M. Kimelman, C. Liu, J. Lopez, P. Meric, C. O'Sullivan, Y. Ostapchuk, L. Phan, S. Ponomarov, V. Schneider, E. Shekhtman, K. Sirotkin, D. Slotta, H. Zhang, G. A. McVean, R. M. Durbin, S. Balasubramaniam, J. Burton, P. Danecek, T. M. Keane, A. Kolb-Kokocinski, S. McCarthy, J. Stalker, M. Quail, J. P. Schmidt, C. J. Davies, J. Gollub, T. Webster, B. Wong, Y. Zhan, A. Auton, C. L. Campbell, Y. Kong, A. Marcketta, R. A. Gibbs, F. Yu, L. Antunes, M. Bainbridge, D. Muzny, A. Sabo, Z. Huang, J. Wang, L. J. M. Coin, L. Fang, X. Guo, X. Jin, G. Li, Q. Li, Y. Li, Z. Li, H. Lin, B. Liu, R. Luo, H. Shao, Y. Xie, C. Ye, C. Yu, F. Zhang, H. Zheng, H. Zhu, C. Alkan, E. Dal, F. Kahveci, G. T. Marth, E. P. Garrison, D. Kural, W.-P. Lee, W. Fung Leong, M. Stromberg, A. N. Ward, J. Wu, M. Zhang, M. J. Daly, M. A. DePristo, R. E. Handsaker, D. M. Altshuler, E. Banks, G. Bhatia, G. del Angel, S. B. Gabriel, G. Genovese, N. Gupta, H. Li, S. Kashin, E. S. Lander, S. A. McCarroll, J. C. Nemesh, R. E. Poplin, S. C. Yoon, J. Lihm, V. Makarov, A. G. Clark, S. Gottipati, A. Keinan, J. L. Rodriguez-Flores, J. O. Korbel, T. Rausch, M. H. Fritz, A. M. Stütz, P. Flicek, K. Beal, L. Clarke, A. Datta, J. Herrero, W. M. McLaren, G. R. S. Ritchie, R. E. Smith, D. Zerbino, X. Zheng-Bradley, P. C. Sabeti, I. Shlyakhter, S. F. Schaffner, J. Vitti, D. N. Cooper, E. V. Ball, P. D. Stenson, D. R. Bentley, B. Barnes, M. Bauer, R. Keira Cheetham, A. Cox, M. Eberle, S. Humphray, S. Kahn, L. Murray, J. Peden, R. Shaw, E. E. Kenny, M. A. Batzer, M. K. Konkel, J. A. Walker, D. G. MacArthur, M. Lek, R. Sudbrak, V. S. Amstislavskiy, R. Herwig, E. R. Mardis, L. Ding, D. C. Koboldt, D. Larson, K. Ye, S. Gravel, The 1000 Genomes Project Consortium, Corresponding authors, Steering committee, Production group, Baylor College of Medicine, BGI-Shenzhen, Broad Institute of MIT and Harvard, Coriell Institute for Medical Research, E. B. I. European Molecular Biology Laboratory, Illumina, Max Planck Institute for Molecular Genetics, McDonnell Genome Institute at Washington University, US National Institutes of Health, University of Oxford, Wellcome Trust Sanger Institute, Analysis group, Affymetrix, Albert Einstein College of Medicine, Bilkent University, Boston College, Cold Spring Harbor Laboratory, Cornell University, European Molecular Biology Laboratory, Harvard University, Human Gene Mutation Database, Icahn School of Medicine at Mount Sinai, Louisiana State University, Massachusetts General Hospital, McGill University, N. National Eye Institute, A global reference for human genetic variation. *Nature* **526**, 68–74 (2015).
20. M. E. Allentoft, M. Sikora, A. Refoyo-Martínez, E. K. Irving-Pease, A. Fischer, W. Barrie, A. Ingason, J. Stenderup, K.-G. Sjögren, A. Pearson, B. Sousa da Mota, B. Schulz Paulsson, A. Halgren, R. Macleod, M. L. S. Jørkov, F. Demeter, L. Sørensen, P. O. Nielsen, R. A. Henriksen, T. Vimala, H. McColl, A. Margaryan, M. Ilardo, A. Vaughn, M. Fischer Mortensen, A. B. Nielsen, M. Ulfeldt Hede, N. N. Johannsen, P. Rasmussen, L. Vinner, G. Renaud, A. Stern, T. Z. T. Jensen, G. Scorrano, H. Schroeder, P. Lysdahl, A. D. Ramsøe, A. Skorobogatov, A. J. Schork, A. Rosengren, A. Ruter, A. Outram, A. A. Timoshenko, A. Buzhilova, A. Coppa, A. Zubova, A. M. Silva, A. J. Hansen, A. Gromov,

A. Logvin, A. B. Gotfredsen, B. Henning Nielsen, B. González-Rabanal, C. Lalueza-Fox, C. J. McKenzie, C. Gaunitz, C. Blasco, C. Liesau, C. Martinez-Labarga, D. V. Pozdnyakov, D. Cuenca-Solana, D. O. Lordkipanidze, D. En'shin, D. C. Salazar-García, T. D. Price, D. Borić, E. Kostyleva, E. V. Veselovskaya, E. R. Usmanova, E. Cappellini, E. Brinch Petersen, E. Kannegaard, F. Radina, F. Eylem Yediay, H. Duday, I. Gutiérrez-Zugasti, I. Merts, I. Potekhina, I. Shevnina, I. Altinkaya, J. Guilaine, J. Hansen, J. E. Aura Tortosa, J. Zilhão, J. Vega, K. Buck Pedersen, K. Tunia, L. Zhao, L. N. Mylnikova, L. Larsson, L. Metz, L. Yepiskoposyan, L. Pedersen, L. Sarti, L. Orlando, L. Slimak, L. Klassen, M. Blank, M. González-Morales, M. Silvestrini, M. Vretemark, M. S. Nesterova, M. Rykun, M. F. Rolfo, M. Szmyt, M. Przybyła, M. Calattini, M. Sablin, M. Dobisíková, M. Meldgaard, M. Johansen, N. Berezina, N. Card, N. A. Saveliev, O. Poshekhonova, O. Rickards, O. V. Lozovskaya, O. Gábor, O. C. Uldum, P. Aurino, P. Kosintsev, P. Courtaud, P. Ríos, P. Mortensen, P. Lotz, P. Persson, P. Bangsgaard, P. de Barros Damgaard, P. Vang Petersen, P. P. Martinez, P. Włodarczak, R. V. Smolyaninov, R. Maring, R. Menduiña, R. Badalyan, R. Iversen, R. Turin, S. Vasilyev, S. Wåhlin, S. Borutskaya, S. Skochina, S. A. Sørensen, S. H. Andersen, T. Jørgensen, Y. B. Serikov, V. I. Molodin, V. Smrcka, V. Merts, V. Appadurai, V. Moiseyev, Y. Magnusson, K. H. Kjær, N. Lynnerup, D. J. Lawson, P. H. Sudmant, S. Rasmussen, T. S. Korneliussen, R. Durbin, R. Nielsen, O. Delaneau, T. Werge, F. Racimo, K. Kristiansen, E. Willerslev, Population genomics of post-glacial western Eurasia. *Nature* **625**, 301–311 (2024).

21. B. Sousa da Mota, S. Rubinacci, D. I. Cruz Dávalos, C. E. G. Amorim, M. Sikora, N. N. Johannsen, M. H. Szmyt, P. Włodarczak, A. Szczepanek, M. M. Przybyła, H. Schroeder, M. E. Allentoft, E. Willerslev, A.-S. Malaspinas, O. Delaneau, Imputation of ancient human genomes. *Nat. Commun.* **14**, 3660 (2023).
22. H. Ringbauer, Y. Huang, A. Akbari, S. Mallick, I. Olalde, N. Patterson, D. Reich, Accurate detection of identity-by-descent segments in human ancient DNA. *Nat. Genet.*, 1–9 (2023).
23. A. Mittnik, C.-C. Wang, J. Svoboda, J. Krause, A Molecular Approach to the Sexing of the Triple Burial at the Upper Paleolithic Site of Dolní Věstonice. *PLOS ONE* **11**, e0163019 (2016).
24. H. Skaletsky, T. Kuroda-Kawaguchi, P. J. Minx, H. S. Cordum, L. Hillier, L. G. Brown, S. Repping, T. Pyntikova, J. Ali, T. Bieri, A. Chinwalla, A. Delehaunty, K. Delehaunty, H. Du, G. Fewell, L. Fulton, R. Fulton, T. Graves, S.-F. Hou, P. Latrielle, S. Leonard, E. Mardis, R. Maupin, J. McPherson, T. Miner, W. Nash, C. Nguyen, P. Ozersky, K. Pepin, S. Rock, T. Rohlfing, K. Scott, B. Schultz, C. Strong, A. Tin-Wollam, S.-P. Yang, R. H. Waterston, R. K. Wilson, S. Rozen, D. C. Page, The male-specific region of the human Y chromosome is a mosaic of discrete sequence classes. *Nature* **423**, 825–837 (2003).
25. W. J. Kent, BLAT—The BLAST-Like Alignment Tool. *Genome Res.* **12**, 656–664 (2002).
26. K. Anastasiadou, M. Silva, T. Booth, L. Speidel, T. Audsley, C. Barrington, J. Buckberry, D. Fernandes, B. Ford, M. Gibson, A. Gilardet, I. Glocke, K. Keefe, M. Kelly, M. Masters, J. McCabe, L. McIntyre, P. Ponce, S. Rowland, J. Ruiz Ventura, P. Swali, F. Tait, D. Walker, H. Webb, M. Williams, A. Witkin, M. Holst, L. Loe, I. Armit, R. Schulting, P. Skoglund, Detection of chromosomal aneuploidy in ancient genomes. *Commun. Biol.* **7**, 1–9 (2024).
27. H. Zhao, Z. Sun, J. Wang, H. Huang, J.-P. Kocher, L. Wang, CrossMap: a versatile tool for coordinate conversion between genome assemblies. *Bioinformatics* **30**, 1006–1007 (2014).
28. R. M. Andrews, I. Kubacka, P. F. Chinnery, R. N. Lightowlers, D. M. Turnbull, N. Howell, Reanalysis and revision of the Cambridge reference sequence for human mitochondrial DNA. *Nat. Genet.* **23**, 147–147 (1999).
29. D. M. Behar, M. van Oven, S. Rosset, M. Metspalu, E.-L. Loogväli, N. M. Silva, T. Kivisild, A. Torroni, R. Villems, A "Copernican" Reassessment of the Human Mitochondrial DNA Tree from its Root. *Am. J. Hum. Genet.* **90**, 675–684 (2012).
30. S. Schönherr, H. Weissensteiner, F. Kronenberg, L. Forer, Haplogrep 3 - an interactive haplogroup classification and analysis platform. *Nucleic Acids Res.* **51**, W263–W268 (2023).

31. D. Koptekin, E. Yüncü, R. Rodríguez-Varela, N. E. Altınışık, N. Psonis, N. Kashuba, S. Yorulmaz, R. George, D. D. Kazancı, D. Kaptan, K. Gürün, K. B. Vural, H. C. Gemici, D. Vassou, E. Daskalaki, C. Karamurat, V. K. Lagerholm, Ö. D. Erdal, E. Kırdök, A. Marangoni, A. Schachner, H. Üstündağ, R. Shengelia, L. Bitadze, M. Elashvili, E. Stravopodi, M. Özbaşaran, G. Duru, A. Nafplioti, C. B. Rose, T. Gencer, G. Darbyshire, A. Gavashelishvili, K. Pitskhelauri, Ö. Çevik, O. Vuruşkan, N. Kyparissi-Apostolika, A. M. Büyükkarakaya, U. Oğuzhanoğlu, S. Günel, E. Tabakaki, A. Aliev, A. Ibrahimov, V. Shadlinski, A. Sampson, G. M. Kılınç, Ç. Atakuman, A. Stamatakis, N. Poulakakis, Y. S. Erdal, P. Pavlidis, J. Storå, F. Özer, A. Götherström, M. Somel, Spatial and temporal heterogeneity in human mobility patterns in Holocene Southwest Asia and the East Mediterranean. *Curr. Biol.* **33**, 41-57.e15 (2023).
32. I. Lazaridis, N. Patterson, A. Mittnik, G. Renaud, S. Mallick, K. Kirsanow, P. H. Sudmant, J. G. Schraiber, S. Castellano, M. Lipson, B. Berger, C. Economou, R. Bollongino, Q. Fu, K. I. Bos, S. Nordenfelt, H. Li, C. de Filippo, K. Prüfer, S. Sawyer, C. Posth, W. Haak, F. Hallgren, E. Fornander, N. Rohland, D. Delsate, M. Francken, J.-M. Guinet, J. Wahl, G. Ayodo, H. A. Babiker, G. Bailliet, E. Balanovska, O. Balanovsky, R. Barrantes, G. Bedoya, H. Ben-Ami, J. Bene, F. Berrada, C. M. Bravi, F. Brisighelli, G. B. J. Busby, F. Cali, M. Churnosov, D. E. C. Cole, D. Corach, L. Damba, G. van Driem, S. Dryomov, J.-M. Dugoujon, S. A. Fedorova, I. Gallego Romero, M. Gubina, M. Hammer, B. M. Henn, T. Hervig, U. Hodoglugil, A. R. Jha, S. Karachanak-Yankova, R. Khusainova, E. Khusnutdinova, R. Kittles, T. Kivisild, W. Klitz, V. Kučinskas, A. Kushniarevich, L. Laredj, S. Litvinov, T. Loukidis, R. W. Mahley, B. Melegh, E. Metspalu, J. Molina, J. Mountain, K. Näkkäläjärvi, D. Nesheva, T. Nyambo, L. Osipova, J. Parik, F. Platonov, O. Posukh, V. Romano, F. Rothhammer, I. Rudan, R. Ruizbakiev, H. Sahakyan, A. Sajantila, A. Salas, E. B. Starikovskaya, A. Tarekegn, D. Toncheva, S. Turdikulova, I. Uktveryte, O. Utevska, R. Vasquez, M. Villena, M. Voevoda, C. A. Winkler, L. Yepiskoposyan, P. Zalloua, T. Zemunik, A. Cooper, C. Capelli, M. G. Thomas, A. Ruiz-Linares, S. A. Tishkoff, L. Singh, K. Thangaraj, R. Villems, D. Comas, R. Sukernik, M. Metspalu, M. Meyer, E. E. Eichler, J. Burger, M. Slatkin, S. Pääbo, J. Kelso, D. Reich, J. Krause, Ancient human genomes suggest three ancestral populations for present-day Europeans. *Nature* **513**, 409–413 (2014).
33. I. Lazaridis, D. Nadel, G. Rollefson, D. C. Merrett, N. Rohland, S. Mallick, D. Fernandes, M. Novak, B. Gamarra, K. Sirak, S. Connell, K. Stewardson, E. Harney, Q. Fu, G. Gonzalez-Fortes, E. R. Jones, S. A. Roodenberg, G. Lengyel, F. Bocquentin, B. Gasparian, J. M. Monge, M. Gregg, V. Eshed, A.-S. Mizrahi, C. Meiklejohn, F. Gerritsen, L. Bejenaru, M. Blüher, A. Campbell, G. Cavalleri, D. Comas, P. Froguel, E. Gilbert, S. M. Kerr, P. Kovacs, J. Krause, D. McGettigan, M. Merrigan, D. A. Merriwether, S. O'Reilly, M. B. Richards, O. Semino, M. Shamoon-Pour, G. Stefanescu, M. Stumvoll, A. Tönjes, A. Torroni, J. F. Wilson, L. Yengo, N. A. Hovhannisyan, N. Patterson, R. Pinhasi, D. Reich, Genomic insights into the origin of farming in the ancient Near East. *Nature* **536**, 419–424 (2016).
34. I. Mathieson, I. Lazaridis, N. Rohland, S. Mallick, N. Patterson, S. A. Roodenberg, E. Harney, K. Stewardson, D. Fernandes, M. Novak, K. Sirak, C. Gamba, E. R. Jones, B. Llamas, S. Dryomov, J. Pickrell, J. L. Arsuaga, J. M. B. de Castro, E. Carbonell, F. Gerritsen, A. Khokhlov, P. Kuznetsov, M. Lozano, H. Meller, O. Mochalov, V. Moiseyev, M. A. R. Guerra, J. Roodenberg, J. M. Vergès, J. Krause, A. Cooper, K. W. Alt, D. Brown, D. Anthony, C. Lalueza-Fox, W. Haak, R. Pinhasi, D. Reich, Genome-wide patterns of selection in 230 ancient Eurasians. *Nature* **528**, 499–503 (2015).
35. H. Ringbauer, J. Novembre, M. Steinrücken, Parental relatedness through time revealed by runs of homozygosity in ancient DNA. *Nat. Commun.* **12**, 5425 (2021).
36. C. C. Chang, C. C. Chow, L. C. Tellier, S. Vattikuti, S. M. Purcell, J. J. Lee, Second-generation PLINK: rising to the challenge of larger and richer datasets. *GigaScience* **4**, 7 (2015).
37. N. Patterson, P. Moorjani, Y. Luo, S. Mallick, N. Rohland, Y. Zhan, T. Genschoreck, T. Webster, D. Reich, Ancient Admixture in Human History. *Genetics* **192**, 1065–1093 (2012).
38. R: The R Project for Statistical Computing. https://www.r-project.org/.

39. O. Barge, H. Azizi Kharanaghi, F. Biglari, B. Moradi, M. Mashkour, M. Tengberg, C. Chataigner, Diffusion of Anatolian and Caucasian obsidian in the Zagros Mountains and the highlands of Iran: Elements of explanation in "least cost path" models. *Quat. Int.* **467**, 297–322 (2018).
40. N. Patterson, A. L. Price, D. Reich, Population Structure and Eigenanalysis. *PLOS Genet.* **2**, e190 (2006).
41. N. E. Altınışık, D. D. Kazancı, A. Aydoğan, H. C. Gemici, Ö. D. Erdal, S. Sarıaltun, K. B. Vural, D. Koptekin, K. Gürün, E. Sağlıcan, D. Fernandes, G. Çakan, M. M. Koruyucu, V. K. Lagerholm, C. Karamurat, M. Özkan, G. M. Kılınç, A. Sevkar, E. Sürer, A. Götherström, Ç. Atakuman, Y. S. Erdal, F. Özer, A. Erim Özdoğan, M. Somel, A genomic snapshot of demographic and cultural dynamism in Upper Mesopotamia during the Neolithic Transition. *Sci. Adv.* **8**, eabo3609 (2022).
42. F. Baumdicker, G. Bisschop, D. Goldstein, G. Gower, A. P. Ragsdale, G. Tsambos, S. Zhu, B. Eldon, E. C. Ellerman, J. G. Galloway, A. L. Gladstein, G. Gorjanc, B. Guo, B. Jeffery, W. W. Kretzschumar, K. Lohse, M. Matschiner, D. Nelson, N. S. Pope, C. D. Quinto-Cortés, M. F. Rodrigues, K. Saunack, T. Sellinger, K. Thornton, H. van Kemenade, A. W. Wohns, Y. Wong, S. Gravel, A. D. Kern, J. Koskela, P. L. Ralph, J. Kelleher, Efficient ancestry and mutation simulation with msprime 1.0. *Genetics* **220**, iyab229 (2022).
43. J. Kelleher, A. M. Etheridge, G. McVean, Efficient Coalescent Simulation and Genealogical Analysis for Large Sample Sizes. *PLOS Comput. Biol.* **12**, e1004842 (2016).
44. Ş. Aktürk, I. Mapelli, M. N. Güler, K. Gürün, B. Katırcıoğlu, K. B. Vural, E. Sağlıcan, M. Çetin, R. Yaka, E. Sürer, G. Atağ, S. S. Çokoğlu, A. Sevkar, N. E. Altınışık, D. Koptekin, M. Somel, Benchmarking kinship estimation tools for ancient genomes using pedigree simulations. *Mol. Ecol. Resour.* **24**, e13960 (2024).
45. Erkin Alaçamlı, Thijessen Naidoo, Şevval Aktürk, Merve N. Güler, Igor Mapelli, Kıvılcım Başak Vural, Mehmet Somel, Helena Malmström, Torsten Günther, READv2: Advanced and user-friendly detection of biological relatedness in archaeogenomics. *bioRxiv*, 2024.01.23.576660 (2024).
46. K. Hanghøj, I. Moltke, P. A. Andersen, A. Manica, T. S. Korneliussen, Fast and accurate relatedness estimation from high-throughput sequencing data in the presence of inbreeding. *GigaScience* **8**, giz034 (2019).
47. G. Dudnik, mltest: Classification Evaluation Metrics, version 1.0.1 (2018); https://cran.r-project.org/web/packages/mltest/index.html.
48. J. M. M. Kuhn, M. Jakobsson, T. Günther, Estimating genetic kin relationships in prehistoric populations. *PLOS ONE* **13**, e0195491 (2018).
49. Ballou: Calculating inbreeding coefficients from... - Google Scholar. https://scholar.google.com/scholar_lookup?hl=en&publication_year=1983&pages=509-520&author=J.+Ballou&title=Genetics+and+Conservation%3A+A+Reference+for+Managing+Wild+Animal+and+Plant+Populations.
50. R. McQuillan, A.-L. Leutenegger, R. Abdel-Rahman, C. S. Franklin, M. Pericic, L. Barac-Lauc, N. Smolej-Narancic, B. Janicijevic, O. Polasek, A. Tenesa, A. K. MacLeod, S. M. Farrington, P. Rudan, C. Hayward, V. Vitart, I. Rudan, S. H. Wild, M. G. Dunlop, A. F. Wright, H. Campbell, J. F. Wilson, Runs of Homozygosity in European Populations. *Am. J. Hum. Genet.* **83**, 359–372 (2008).
51. M. Rivollat, A. B. Rohrlach, H. Ringbauer, A. Childebayeva, F. Mendisco, R. Barquera, A. Szolek, M. Le Roy, H. Colleran, J. Tuke, F. Aron, M.-H. Pemonge, E. Späth, P. Télouk, L. Rey, G. Goude, V. Balter, J. Krause, S. Rottier, M.-F. Deguilloux, W. Haak, Extensive pedigrees reveal the social organization of a Neolithic community. *Nature* **620**, 600–606 (2023).
52. J. R. Adrion, C. B. Cole, N. Dukler, J. G. Galloway, A. L. Gladstein, G. Gower, C. C. Kyriazis, A. P. Ragsdale, G. Tsambos, F. Baumdicker, J. Carlson, R. A. Cartwright, A. Durvasula, I. Gronau, B. Y. Kim, P. McKenzie, P. W. Messer, E. Noskova, D. Ortega-Del Vecchyo, F. Racimo, T. J. Struck, S. Gravel, R. N. Gutenkunst, K. E. Lohmueller, P. L. Ralph, D. R. Schrider, A. Siepel, J. Kelleher, A. D.

Kern, A community-maintained standard library of population genetic models. *eLife* **9**, e54967 (2020).

53. M. E. Lauterbur, M. I. A. Cavassim, A. L. Gladstein, G. Gower, N. S. Pope, G. Tsambos, J. Adrion, S. Belsare, A. Biddanda, V. Caudill, J. Cury, I. Echevarria, B. C. Haller, A. R. Hasan, X. Huang, L. N. M. Iasi, E. Noskova, J. Obsteter, V. A. C. Pavinato, A. Pearson, D. Peede, M. F. Perez, M. F. Rodrigues, C. C. Smith, J. P. Spence, A. Teterina, S. Tittes, P. Unneberg, J. M. Vazquez, R. K. Waples, A. W. Wohns, Y. Wong, F. Baumdicker, R. A. Cartwright, G. Gorjanc, R. N. Gutenkunst, J. Kelleher, A. D. Kern, A. P. Ragsdale, P. L. Ralph, D. R. Schrider, I. Gronau, Expanding the stdpopsim species catalog, and lessons learned for realistic genome simulations. *eLife* **12**, RP84874 (2023).

54. K. A. Frazer, D. G. Ballinger, D. R. Cox, D. A. Hinds, L. L. Stuve, R. A. Gibbs, J. W. Belmont, A. Boudreau, P. Hardenbol, S. M. Leal, S. Pasternak, D. A. Wheeler, T. D. Willis, F. Yu, H. Yang, C. Zeng, Y. Gao, H. Hu, W. Hu, C. Li, W. Lin, S. Liu, H. Pan, X. Tang, J. Wang, W. Wang, J. Yu, B. Zhang, Q. Zhang, H. Zhao, H. Zhao, J. Zhou, S. B. Gabriel, R. Barry, B. Blumenstiel, A. Camargo, M. Defelice, M. Faggart, M. Goyette, S. Gupta, J. Moore, H. Nguyen, R. C. Onofrio, M. Parkin, J. Roy, E. Stahl, E. Winchester, L. Ziaugra, D. Altshuler, Y. Shen, Z. Yao, W. Huang, X. Chu, Y. He, L. Jin, Y. Liu, Y. Shen, W. Sun, H. Wang, Y. Wang, Y. Wang, X. Xiong, L. Xu, M. M. Y. Waye, S. K. W. Tsui, H. Xue, J. T.-F. Wong, L. M. Galver, J.-B. Fan, K. Gunderson, S. S. Murray, A. R. Oliphant, M. S. Chee, A. Montpetit, F. Chagnon, V. Ferretti, M. Leboeuf, J.-F. Olivier, M. S. Phillips, S. Roumy, C. Sallée, A. Verner, T. J. Hudson, P.-Y. Kwok, D. Cai, D. C. Koboldt, R. D. Miller, L. Pawlikowska, P. Taillon-Miller, M. Xiao, L.-C. Tsui, W. Mak, Y. Qiang Song, P. K. H. Tam, Y. Nakamura, T. Kawaguchi, T. Kitamoto, T. Morizono, A. Nagashima, Y. Ohnishi, A. Sekine, T. Tanaka, T. Tsunoda, P. Deloukas, C. P. Bird, M. Delgado, E. T. Dermitzakis, R. Gwilliam, S. Hunt, J. Morrison, D. Powell, B. E. Stranger, P. Whittaker, D. R. Bentley, M. J. Daly, P. I. W. de Bakker, J. Barrett, Y. R. Chretien, J. Maller, S. McCarroll, N. Patterson, I. Pe’er, A. Price, S. Purcell, D. J. Richter, P. Sabeti, R. Saxena, S. F. Schaffner, P. C. Sham, P. Varilly, D. Altshuler, L. D. Stein, L. Krishnan, A. Vernon Smith, M. K. Tello-Ruiz, G. A. Thorisson, A. Chakravarti, P. E. Chen, D. J. Cutler, C. S. Kashuk, S. Lin, G. R. Abecasis, W. Guan, Y. Li, H. M. Munro, Z. Steve Qin, D. J. Thomas, G. McVean, A. Auton, L. Bottolo, N. Cardin, S. Eyheramendy, C. Freeman, J. Marchini, S. Myers, C. Spencer, M. Stephens, P. Donnelly, L. R. Cardon, G. Clarke, D. M. Evans, A. P. Morris, B. S. Weir, T. Tsunoda, T. Johnson, J. C. Mullikin, S. T. Sherry, M. Feolo, A. Skol, H. Zhang, C. Zeng, H. Zhao, I. Matsuda, Y. Fukushima, D. R. Macer, E. Suda, C. N. Rotimi, C. A. Adebamowo, I. Ajayi, T. Aniagwu, P. A. Marshall, C. Nkwodimmah, C. D. M. Royal, M. F. Leppert, M. Dixon, A. Peiffer, R. Qiu, A. Kent, K. Kato, N. Niikawa, I. F. Adewole, B. M. Knoppers, M. W. Foster, E. Wright Clayton, J. Watkin, R. A. Gibbs, J. W. Belmont, D. Muzny, L. Nazareth, E. Sodergren, G. M. Weinstock, D. A. Wheeler, I. Yakub, S. B. Gabriel, R. C. Onofrio, D. J. Richter, L. Ziaugra, B. W. Birren, M. J. Daly, D. Altshuler, R. K. Wilson, L. L. Fulton, J. Rogers, J. Burton, N. P. Carter, C. M. Clee, M. Griffiths, M. C. Jones, K. McLay, R. W. Plumb, M. T. Ross, S. K. Sims, D. L. Willey, Z. Chen, H. Han, L. Kang, M. Godbout, J. C. Wallenburg, P. L’Archevêque, G. Bellemare, K. Saeki, H. Wang, D. An, H. Fu, Q. Li, Z. Wang, R. Wang, A. L. Holden, L. D. Brooks, J. E. McEwen, M. S. Guyer, V. Ota Wang, J. L. Peterson, M. Shi, J. Spiegel, L. M. Sung, L. F. Zacharia, F. S. Collins, K. Kennedy, R. Jamieson, The International HapMap Consortium, Genotyping centres: Perlegen Sciences, Baylor College of Medicine and ParAllele BioScience, Beijing Genomics Institute, Broad Institute of Harvard and Massachusetts Institute of Technology, Chinese National Human Genome Center at Beijing, Chinese National Human Genome Center at Shanghai, Chinese University of Hong Kong, Hong Kong University of Science and Technology, Illumina, McGill University and Génome Québec Innovation Centre, University of California at San Francisco and Washington University, University of Hong Kong, University of Tokyo and RIKEN, Wellcome Trust Sanger Institute, Analysis groups: Broad Institute, Cold Spring Harbor Laboratory, Johns Hopkins University School of Medicine, University of Michigan, University of Oxford, W. T. C. for H. G. University of Oxford, RIKEN, US

National Institutes of Health, US National Institutes of Health National Center for Biotechnology Information, Community engagement/public consultation and sample collection groups: Beijing Normal University and Beijing Genomics Institute, E. E. I. Health Sciences University of Hokkaido and Shinshu University, Howard University and University of Ibadan, University of Utah, legal and social issues: C. A. of S. S. Ethical, Genetic Interest Group, Kyoto University, Nagasaki University, University of Ibadan School of Medicine, University of Montréal, University of Oklahoma, Vanderbilt University, Wellcome Trust, SNP discovery: Baylor College of Medicine, Washington University, Scientific management: Chinese Academy of Sciences, Genome Canada, Génome Québec, C. Japanese Ministry of Education Sports, Science and Technology, Ministry of Science and Technology of the People's Republic of China, The Human Genetic Resource Administration of China, The SNP Consortium, A second generation human haplotype map of over 3.1 million SNPs. *Nature* **449**, 851–861 (2007).

55. J. Kamm, J. Terhorst, R. Durbin, Y. S. Song, Efficiently Inferring the Demographic History of Many Populations With Allele Count Data. *J. Am. Stat. Assoc.* **115**, 1472–1487 (2020).
56. G. M. Kılınç, A. Omrak, F. Özer, T. Günther, A. M. Büyükkarakaya, E. Bıçakçı, D. Baird, H. M. Dönertaş, A. Ghalichi, R. Yaka, D. Koptekin, S. C. Açan, P. Parvizi, M. Krzewińska, E. A. Daskalaki, E. Yüncü, N. D. Dağtaş, A. Fairbairn, J. Pearson, G. Mustafaoğlu, Y. S. Erdal, Y. G. Çakan, İ. Togan, M. Somel, J. Storå, M. Jakobsson, A. Götherström, The Demographic Development of the First Farmers in Anatolia. *Curr. Biol. CB* **26**, 2659–2666 (2016).
57. M. Caballero, D. N. Seidman, Y. Qiao, J. Sannerud, T. D. Dyer, D. M. Lehman, J. E. Curran, R. Duggirala, J. Blangero, S. Carmi, A. L. Williams, Crossover interference and sex-specific genetic maps shape identical by descent sharing in close relatives. *PLOS Genet.* **15**, e1007979 (2019).
58. C. L. Campbell, N. A. Furlotte, N. Eriksson, D. Hinds, A. Auton, Escape from crossover interference increases with maternal age. *Nat. Commun.* **6**, 6260 (2015).
59. M. Vasic, M. Siebrecht, C. Tsoraki, R. Veropoulidou, "Beads and pendants in life and death: insights into the production, use and deposition of ornamental technologies at Çatalhöyük" (2021), pp. 215–246.
60. J. A. Pearson, S. D. Haddow, S. W. Hillson, C. J. Knüsel, C. S. Larsen, J. W. Sadvari, Stable carbon and nitrogen isotope analysis and dietary reconstruction through the life course at Neolithic Çatalhöyük, Turkey. *J. Soc. Archaeol.* **15**, 210–232 (2015).
61. B. t. Fuller, J. l. Fuller, D. a. Harris, R. e. m. Hedges, Detection of breastfeeding and weaning in modern human infants with carbon and nitrogen stable isotope ratios. *Am. J. Phys. Anthropol.* **129**, 279–293 (2006).
62. Z. Pochon, N. Bergfeldt, E. Kırdök, M. Vicente, T. Naidoo, T. van der Valk, N. E. Altınışık, M. Krzewińska, L. Dalén, A. Götherström, C. Mirabello, P. Unneberg, N. Oskolkov, aMeta: an accurate and memory-efficient ancient metagenomic profiling workflow. *Genome Biol.* **24**, 242 (2023).
63. F. P. Breitwieser, D. N. Baker, S. L. Salzberg, KrakenUniq: confident and fast metagenomics classification using unique k-mer counts. *Genome Biol.* **19**, 198 (2018).
64. D. H. Huson, A. F. Auch, J. Qi, S. C. Schuster, MEGAN analysis of metagenomic data. *Genome Res.* **17**, 377–386 (2007).
65. G. Marçais, C. Kingsford, A fast, lock-free approach for efficient parallel counting of occurrences of k-mers. *Bioinformatics* **27**, 764–770 (2011).
66. L. Breiman, Random Forests. *Mach. Learn.* **45**, 5–32 (2001).
67. A. Liaw, M. Wiener, Classification and regression by randomForest. *R News* **2**, 18–22 (2002).
68. J. Oksanen, G. L. Simpson, F. G. Blanchet, R. Kindt, P. Legendre, P. R. Minchin, R. B. O'Hara, P. Solymos, M. H. H. Stevens, E. Szoecs, H. Wagner, M. Barbour, M. Bedward, B. Bolker, D. Borcard, G. Carvalho, M. Chirico, M. D. Caceres, S. Durand, H. B. A. Evangelista, R. FitzJohn, M. Friendly, B. Furneaux, G. Hannigan, M. O. Hill, L. Lahti, D. McGlinn, M.-H. Ouellette, E. R. Cunha, T. Smith, A.

Stier, C. J. F. T. Braak, J. Weedon, vegan: Community Ecology Package, version 2.6-6.1 (2024); https://cran.r-project.org/web/packages/vegan/index.html.

69. J. Adserias-Garriga, M. Hernández, N. M. Quijada, D. Rodríguez Lázaro, D. Steadman, J. Garcia-Gil, Daily thanatomicrobiome changes in soil as an approach of postmortem interval estimation: An ecological perspective. *Forensic Sci. Int.* **278**, 388–395 (2017).
70. A. L. Emmons, A. Z. Mundorff, K. M. Hoeland, J. Davoren, S. W. Keenan, D. O. Carter, S. R. Campagna, J. M. DeBruyn, Postmortem Skeletal Microbial Community Composition and Function in Buried Human Remains. *mSystems* **7**, e00041-22 (2022).
71. L. C. Reimer, J. Sardà Carbasse, J. Koblitz, C. Ebeling, A. Podstawka, J. Overmann, BacDive in 2022: the knowledge base for standardized bacterial and archaeal data. *Nucleic Acids Res.* **50**, D741–D746 (2022).
